# Discovery and characterization of stereodefined PMO-gapmers targeting tau

**DOI:** 10.1101/2024.05.09.591947

**Authors:** Kunihiko Kanatsu, Yoshinori Takahashi, Tetsuya Sakaguchi, Dae-Shik Kim, Miki Murota, Mingde Shan, Kazuki Fukami, Wataru Itano, Kenji Kikuta, Hikaru Yoshimura, Toshiki Kurokawa, Yuko Nagayama, Rena Ishikawa, Ryo Dairiki, Zhi Zhou, Kristen Sanders, Jacob Stupalski, So Yasui, Diana Liu, Farid Benayoud, Hui Fang, Enxuan Jing, Makoto Ogo, Francis G. Fang, John Wang, Hyeong-wook Choi

## Abstract

Antisense oligonucleotides (ASOs) are an important class of therapeutics to treat genetic diseases, and expansion of this modality to neurodegenerative disorders has been an active area of research. To realize chronic administration of ASO therapeutics to treat neurogenerative diseases, new chemical modifications improving activity and safety profile are still needed. Furthermore, it is highly desirable to develop a single stereopure ASO with defined activity and safety profile to avoid any efficacy and safety concerns due to the batch-to-batch variation in the composition of diastereomers. Herein, a stereopure PMO-gapmer was developed as a new construct to improve safety and stability by installing charge-neutral PMOs at the wing region and by fully controlling phosphorus stereochemistries. The developed stereopure PMO-gapmer construct was applied to the discovery of ASO candidates for the reduction of microtubule-associated protein tau (*MAPT*, tau). Sequence screening targeting *MAPT* followed by screening of optimal phosphorus stereochemistry identified stereopure development candidates. While evaluating the stereopure PMO-gapmers, we observed a dramatic difference in safety profile among stereoisomers in which only one phosphorus stereochemistry differs. These results further highlight the benefits of developing stereopure ASOs as safe and well-characterized candidates for clinical studies.

## INTRODUCTION

Antisense oligonucleotides (ASOs) are an important class of therapeutics to treat diseases with genetically validated targets (1). ASOs exhibit therapeutic activity by knocking down the target genes or modulating splicing events. For the knockdown of target genes, gapmer ASOs (or gapmers) inducing RNase H-mediated degradation of the target RNA have been employed. The gapmers have a central DNA gap region for engagement with RNase H, and 5’ and 3’-wing regions of modified nucleotides (e.g., 2’-MOE, LNA and cEt) to improve the binding affinity to target genes as well as to increase stability against exonucleases (Figure 1). Furthermore, phosphorothioate linkages are preferentially used as the internucleotide linkages throughout the gapmers to improve the stability and pharmacokinetics (PK) profile of ASOs. While conventional gapmer configurations have been successfully used in approved drugs, there are still concerns regarding the toxicity caused by nonselective protein binding associated with phosphorothioate linkages. Efforts have been made to mitigate safety risks by adding a certain number of phosphodiesters linkages, adding a chemically modified nucleotide at a specific position of the gap region, or more recently, introducing charge-neutral linkages (2, 3, 4, 5, 6, 7). Incorporation of charge-neutral linkages like phosphoryl guanidine in the wing and/or gap region of gapmers was demonstrated to improve ASO properties (8, 9). Phosphorodiamidate morpholino oligomers (PMO) are another class of clinically validated, chemical modified oligonucleotides with neutral linkages. Considering the general safety benefits of PMO modification (10) and its resistance to nucleases due to the unnatural phosphorodiamidate linkages, the employment of PMO modification at the wing region of gapmers could be beneficial. However, the use of PMOs has been limited to modulation of splicing events, and its incorporation to gapmer constructs has not yet been reported.

**Figure 1.**
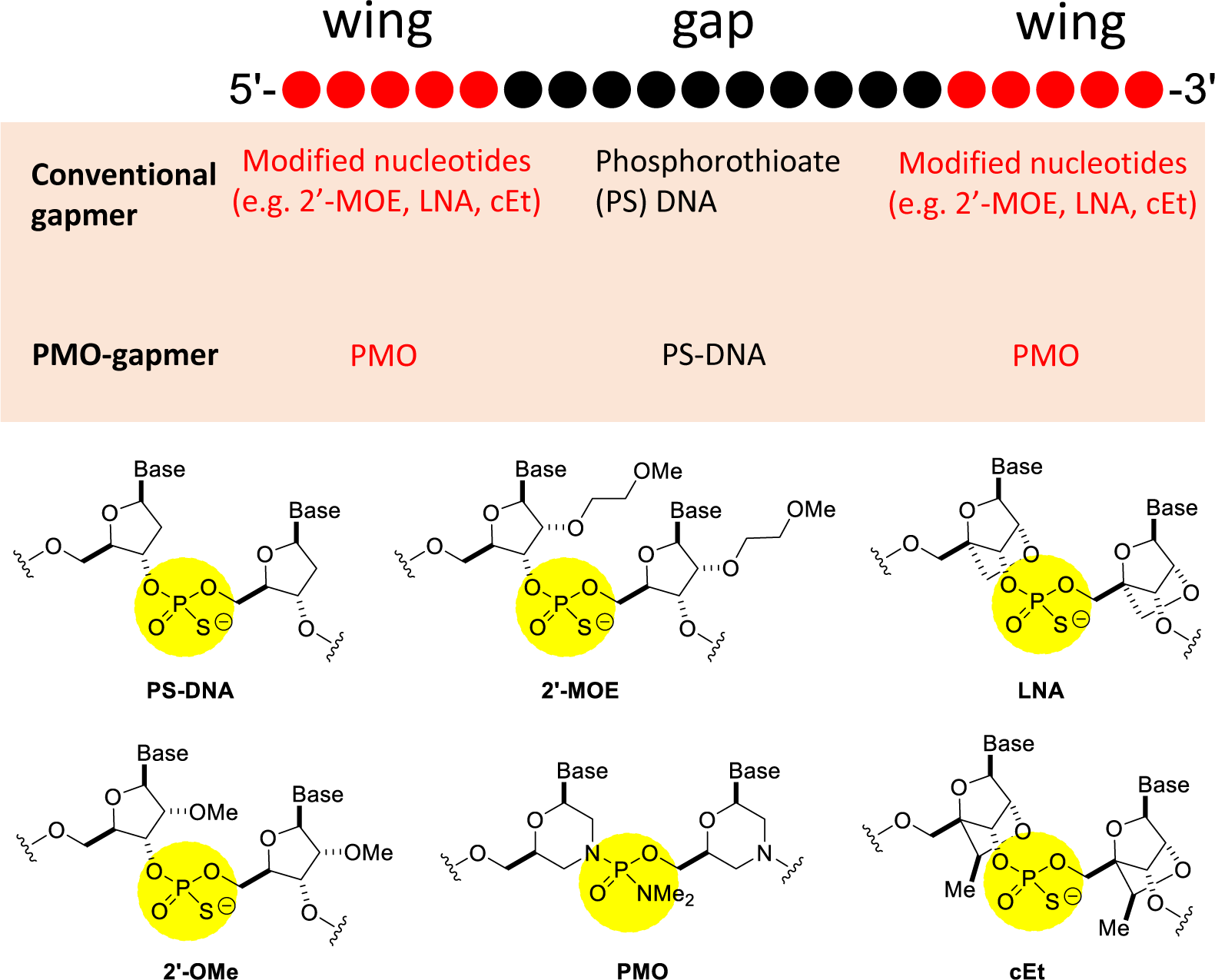
General structure of gapmer ASOs and configuration of PMO-gapmers were represented with a 5-10-5 construct (5’-wing of 5 modified nucleotides, DNA gap of 10-mer DNA and 3’-wing of 5 modified nucleotides). Red dots and black dots represent chemically modified nucleotides and DNA monomers, respectively. Commonly used chemical modifications are depicted and the chiral phosphorus atoms are marked with yellow circles.

Replacement of a phosphodiester linkage with a phosphorothioate or phosphorodiamidate linkage makes the phosphorus atom chiral and generates two possible diastereomers at each internucleotide linkage. It has been hypothesized that the phosphorus stereochemistry influences the activity, stability, and safety of ASOs and could be used for fine tuning of ASO properties (11, 12, 13, 14, 15, 16, 17). However, the investigation of this stereochemical hypothesis has been hampered by two related obstacles: 1) there existed the notion that issues of phosphorous stereochemistry were of secondary or even no importance, 2) The chemistry technologies to stereoselectively and practically construct single molecule ASOs were not available. Recently, advances in stereoselective phosphorylation chemistry have emerged to enable practical stereopure ASO synthesis. Such technologies include Stec’s oxathiaphospholane chemistry (18, 19, 20), Beaucage’s acyloxazaphospholidines (21), Wada’s oxazaphospholidines (22, 23, 24), and Baran’s PSI chemistry (25, 26, 27). Additionally, it was reported that the clinical failure of mongersen was traced to inconsistent biological activity exhibited in different clinical batches (28, 29, 30). The batch-to-batch variability was not discerned by the standard small molecule analytical release testing utilized at that time. Presumably, different compositions of mongersen’s 2^20^ possible phohphorothioate diastereomers was an uncontrolled variable in the “Good Manufacturing Practices (GMP)” production of the different clinical batches. The obvious implication is that a controlled synthesis and comparison of single molecule entities would be essential in order to scientifically explore and manage the safety and efficacy of ASOs.

Tau is an important target for tauopathies including Alzheimer’s disease (AD). Tau protein aggregates upon phosphorylation in pathological conditions to result in neurofibrillary tangles (NFT), which is a hallmark of AD pathology. The level of NFT correlates well with cognitive deficit and neuronal loss (31). In addition, there is strong genetic evidence between tau mutations and frontotemporal dementias (32, 33). DeVos *et. al.* reported that intracerebroventricular (ICV) infusion of a tau targeting ASO reduced already formed phosphorylated tau and prevented neuronal loss in PS19 mouse (34). Moreover, Ionis-MAPTRx (BIIB-080), an ASO targeting *MAPT*, reduced not only tau protein production but also the tau tangles measured by tau positron-emission tomography (PET) in clinical phase 1 study, providing further supporting evidence for the potential of a tau targeting ASO therapy (35).

Herein, a new gapmer construct, called PMO-gapmer, consisting of charge neutral PMO wing regions and a phosphorothioate DNA gap region is described. A synthesis of a stereodefined PMO-gapmer with complete control of phosphorus stereochemistry is established by combining a new method for controlling PMO stereochemistry with novel applications of phosphorothioate stereocontrol. A solution-phase synthetic method for stereodefined PMO-gapmers features a block coupling of 5’ and 3’ fragments and enables a convergent synthesis. A series of stereo-patterns in the DNA gap region were evaluated to select optimal stereochemistries in terms of in vivo activity, stability in brain homogenates and immunogenicity. ASO-486-R5 exhibits good activity, good stability in human brain homogenates, and no cytokine release in human peripheral blood mononuclear cells (PBMCs) and thus warrants further evaluation. Moreover, remarkable differences in efficacy and safety profile is reported for specific stereochemical patterns of PMO-gapmers. Specifically, a single phosphorous stereogenic center of ASO-409-R3 is found to impart lethality or safety in the same PMO-gapmer sequence.

## MATERIAL AND METHODS

### Synthesis of oligonucleotides

Stereorandom PMO-gapmers for this study were prepared by a solid-phase synthesis following the procedures described in Supplementary Information. The detailed procedures for the solution-phase synthesis of stereopure PMO-gapmers (ASO-486-R5-S, ASO-486-R5-R and ASO-486-R5-M) and ^31^P NMR and high-performance liquid chromatography (HPLC) profiles of selected compounds are provided in Supplementary Information. All other stereopure PMO-gapmers were prepared following the same procedure.

### In vitro screening

The SH-SY5Y cells (European Collection of Authenticated Cell Cultures) were cultured in Dulbecco’s Modified Eagle’s Medium/Nutrient Mixture F-12 (high-glucose) supplemented with Non-Essential Amino Acids and 10% fetal bovine serum (All materials were obtained Thermo Fisher Scientific, Waltham, MA, USA). Cultures were maintained at 37 °C in a 5% CO_2_ tissue culture incubator.

Cultured SH-SY5Y cells were transfected using Endo-Porter (Gene Tools, Philomath, OR, USA) with 10-300 nM antisense oligonucleotide. Two days post-transfection, RNA was isolated from the cell lysis using Maxwell® RSC simply RNA Cells/Tissue Kit (Promega, Madison, WI, USA) and cDNA was synthesized. Tau mRNA levels were measured by real-time quantitative PCR (qPCR) using TaqMan probes specific to Human *MAPT* (Assay ID Hs00902194_m1) and Human *GAPDH* (Assay ID HS99999905_m1). *GAPDH* mRNA levels were used as internal control for normalization. Results are presented as relative mRNA expression to the vehicle control.

### In vivo screening

Adult male human *MAPT* knock-in (hTau KI) mice (36) were used for in vivo evaluations. All surgical procedures were performed under butorphanol, medetomidine and midazolam anesthesia, and were approved by the Institutional Animal Care and Use Committee (IACUCC) and carried out in accordance with, as appropriate, the Animal Experimentation Regulations of Eisai Co., Ltd. ASOs (30-100 µg in 10 µL saline solution) were administered to mice by intracerebroventricular (ICV) bolus injection (n = 3-12 mice per group) into the left lateral ventricle. A control group was treated with saline with ICV injection. At necropsy, all surviving animals were weighed. Tissues were collected 3, 7, 21, or 56 days after oligonucleotide administration. RNA was extracted from hippocampus, striatum, and cortex and examined for human tau mRNA expression using real-time qPCR analysis. Human tau mRNA levels were measured using TaqMan probes specific to Human *MAPT* (Assay ID Hs00902194_m1) and were normalized by mouse Glyceraldehyde-3-Phoshate Dehydrogenase (*Gapdh*, Assay ID Mm99999915_g1). Results were calculated as normalized human tau mRNA percent inhibition of the vehicle control. The hemibrain were collected from animals treated with ASO-409-SSR2-M at 30 μg and 60 μg in 3-week study, and fixed in 10% neutral buffered formalin and paraffin-embedded sections were prepared. Tissue sections were stained with hematoxylin and eosin.

### In vitro stability in human brain homogenate

Human brain homogenate was obtained from BioIVT (Westbury, NY, USA). Gender mixed human brains (5 males and 5 females) were homogenized in 100 mM Tris HCl (pH 8) buffer including 1 mM magnesium acetate and antimycotic agents. The diluted homogenate (10 mg/mL) in the same buffer was incubated for 1 day at 37 °C. ASO solution (100 µM in 5 µL) was added to 500 µL of the pre-incubated homogenate and then incubated at 37 °C for 144 hours. The homogenate samples were sequentially collected at 0, 24, 48, 72, and 144 h during incubation. The collected samples were immediately mixed with Clarity OTX lysis-loading buffer (Phenomenex, Torrance, CA, USA) and stored at -80 °C. Solid-phase extraction was performed using Clarity OTX 100 mg, 96 well plate (Phenomenex) following manufacturer’s procedure. Samples were mixed with internal standard (IS, an in-house synthesized 18-mer oligonucleotide with different chemistry and sequence from ASO-486) solution and loaded onto clarity OTX plate equilibrated with 50 mM ammonium acetate (pH 4.5). After centrifugation, the plate was washed twice with 50 mM ammonium acetate (pH 4.5) in 50% acetonitrile. Analyte was eluted twice with 100 mM ammonium bicarbonate (pH 9.5) in 50% acetonitrile. The eluted solutions were dried and reconstituted with a solvent consisting of 10 mM triethylamine, 200 mM HFIP, 10% methanol, and 0.02% acetylacetone. The solution was analyzed by ion-pair liquid chromatography coupled with mass spectrometry (LC-MS) system consists of Nexera system (Shimadzu, Kyoto, Japan) and Q Exactive HF System (Thermo Fisher Scientific, Waltham, MA, USA). The peak area ratio (PAR) of each ASO-486 to IS was calculated. The residual percentages were calculated using percentage of PARs at 24, 48, 72, and 144 h of PAR at 0 h.

### Preliminary cytokine release assay using hPBMC

Human PBMCs were obtained from AllCells (donor ID 14078). Upon receiving the live cells, fresh hPBMCs were plated in 96 well plates with the seeding density at 5x10^5^ cells/well in 180 μL RMPI160 media supplemented with 10% FBS. Total 20µL ASOs and/or positive control (ODN2006, InvivoGen Cat# tlrl-2006-1) at 10x concentration were added to the cell plate to have final dosing concentrations at 0, 0.03, 0.1, 3, and 10 μM with 1 well PBMCs per ASO per treatment concentration. Cells were treated for 24 hours then cell plate was briefly spun, and supernatant was collected for MSD assay on different cytokine release in the media post-treatment. MSD V-plex proinflammatory panel 1 ELISA assay was performed following vendor protocol (MSD Cat# K15049D-1: Analytes include IFN-γ, IL-1β, IL-2, IL-4, IL-6, IL-8, IL-10, IL-12p70, IL-13, TNF-α,) using MESO QuickPlex SQ 120MM MSD reader.

### RNase H1 cleavage assay

The RNase H1 cleavage assay was based on published protocol (13). Human RNase H1 (Abcam, AB153634) was diluted to 10 ng/μL (10X) in buffer containing 100 mM Tris–HCl, 50 mM NaCl, 30% glycerol and 10 mM DTT at pH 7.4. RNA probe (5’-FAM-UGUCUAAGGGUCAUCUGCU-3’; IDTDNA Cat#: 438813816) was mixed in equal ratio with different ASOs at 0.33uM and annealed in reaction buffer containing 20 mM Tris–HCl, 50 mM NaCl, 10 mM MgCl_2_ and 10 mM DTT at pH 7.4. Either RNase buffer control or RNase H1 protein (1 ng) was added to each duplex solution and then was incubated at 37 °C for 10 min using a BioRad thermocycler. The reaction was stopped by adding 10 μL stop solution containing 8 M urea and 120 mM EDTA for every 20 μL of reaction mixture. Samples were heated to 95 °C for 5 min and separated on 15% TBE with urea polyacrilamide gel (BioRad Cat# 3450093) at 200 V for 60 min. The signal was detected using a LiCor Odyssey M Imager.

## RESULTS

### Synthesis of stereorandom PMO-gapmers for sequence screening

Stereorandom PMO-gapmers, for sequence screening, were prepared by the solid phase synthesis using a conventional chlorophosphoramidate chemistry for the construction of PMO wings and a phosphoramidite chemistry for the gap DNA region with phosphorothioate linkages. Since conventional PMO synthesis is conducted from 5’ to 3’ direction, unconventional 5’-phosphoramidite DNA monomers were used to match the synthetic direction (Supplementary Figure S1). For the connection of PMO-DNA junction and DNA-PMO junction at the 5’ or 3’-terminal of the gap region, a DNA monomer having 5’-chlorophosphoramidate and a PMO monomer having 5’-phosphoramidite were used, respectively. The developed solid phase synthesis was successfully implemented to synthesize selected 65 PMO-gapmers for in vitro and in vivo screening.

### Synthesis of stereodefined PMO-gapmers

Stereopure PMO wings were prepared by the previously reported synthetic protocols (Supplementary Figure S2) (37). Stereospecific coupling reactions with the stereopure activated PMO monomers were accomplished in the presence of a non-nucleophilic base in a polar aprotic solvent at room temperature: 1,2,2,6,6-pentamethylpioperidine (PMP) was selected as the optimal base and 1,3-dimethyl-2-imidazolidinone (DMI) as the best reaction solvent. The use of *N*-methylimidazole or *N*-ethylmorpholine, which were employed in conventional stereorandom PMO synthesis, caused epimerization or generated side products. Good solubility as well as faster reaction rate was observed in DMI solvent: DMI gave better coupling results than other tested solvents including acetonitrile, *N*-methylpyrrolidone (NMP) and methyl sulfoxide (DMSO). Under the optimized conditions, efficient stereospecific coupling reactions were achieved without epimerization. All the possible 16 PMO dinucleotides were synthesized by the developed method to confirm the integrity of phosphorus stereochemistry during the coupling reactions and determine the absolute stereochemistry (Supplementary Table S1). X-ray crystal structure of a TA dinucleotide was obtained to unambiguously determine the absolute stereochemistry of phosphorus atom and to deduce the absolute stereochemistry of activated A monomer given that the presumed mechanism of the coupling reaction would involve inversion of configuration at the phosphorus stereogenic center (37). The phosphorus stereochemistry of the rest of the dinucleotides and the activated monomers were then assigned based on the relative ^31^P NMR chemical shift comparison with those of PMO A dinucleotide and monomer (38, 39).

To assess the correlation of PMO phosphorus stereochemistry with binding affinity, A, T, C and G 10-mers having homogeneous phosphorus stereochemistry (i.e., all-*R*p and all-*S*p) were prepared from the stereopure monomers (Supplementary Figure S3). Tm measurements with the corresponding complimentary RNA 10-mer were conducted for each stereoisomer. All-*S*p 10-mers consistently exhibited higher Tm than the corresponding all-*R*p 10-mers thus demonstrating that *S*p phosphorodiamidate linkages generally have better binding affinity than the corresponding *R*p stereoisomers.

For the synthesis of stereodefined PMO-gapmers, P(V) chemistry was selected to ensure stereochemical integrity during the coupling reactions: the stereopure PMO synthesis was used for the stereopure PMO wings, and PSI chemistry developed by the Baran group was employed for the synthesis of DNA gap region with stereopure phosphorothioate linkages (25, 26, 27). The coupling reactions with PSI activated DNA monomers resulted in free phosphorothioates, which were not compatible with chlorophosphoramidate chemistry for PMO synthesis. Thus, the linear synthesis of stereodefined PMO-gapmers as for the stereorandom PMO-gapmers was not feasible. Instead, a convergent synthetic method featuring a block coupling of two oligomers, a 5’-fragment and a 3’-fragment, was developed (Figure 2A). As the coupling reaction between 3’-PSI activated monomer and 5’-OH showed better coupling efficiency in a model system, the block coupling of 5’-fragment (5 or 6-mer) having 3’-PSI activation with 3’-fragment (13 or 12-mer) having 5’-OH was selected as the key step. To make the key block coupling reaction more efficient in a practical scale, a solution phase synthesis was devised.

**Figure 2.**
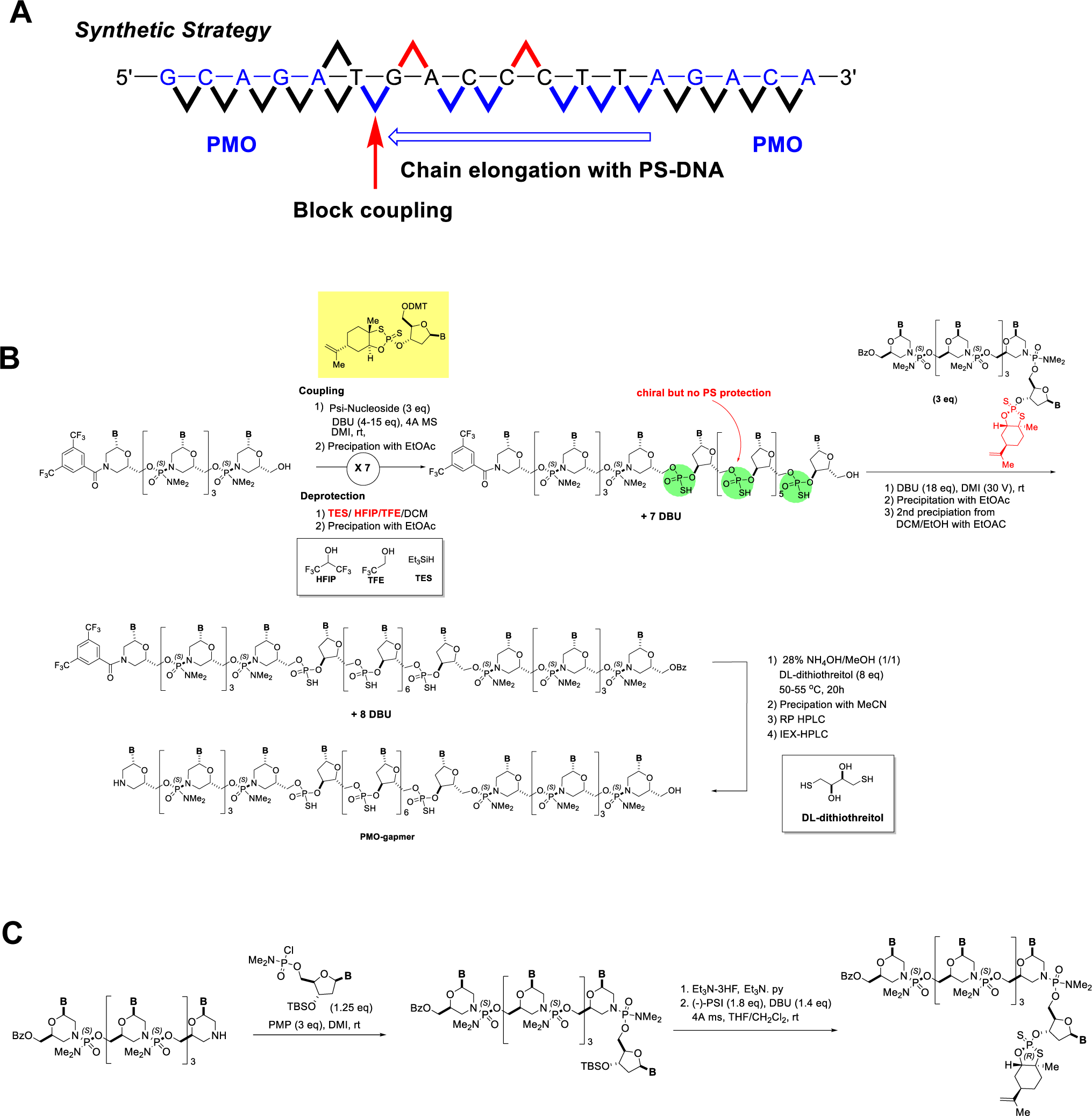
A convergent synthesis of stereopure PMO-gapmers via a block coupling of PSI activated 5’-fragment (6-mer) and 3’-fragment (12-mer). (A) Schematic representation of synthetic strategy for stereopure 5-8-5 PMO-gapmers using ASO-409-SSR2 as an example. (B) A synthetic scheme for the solution phase synthesis of stereopure PMO-gapmers. (C) Synthesis of PSI-activated 5’-fragment (6-mer).

The synthesis of a 3’-fragment commenced with the stereopure PMO synthesis (Figure 2B). After synthesis of the desired length of PMOs, all the protecting groups were removed with ammonium hydroxide, and the morpholine at the 3’-terminal was protected with bistrifluoromethyl benzoyl group. The stereopure PMO wings facilitated the precipitation of all the intermediates for the synthesis of 3’-fragment upon addition of less polar solvents to avoid column chromatographic purifications of the highly polar intermediates. The DNA gap region was elongated from the deprotected 3’-PMO wing with 3’-PSI activated DNA monomers to give the desired length of 3’-fragment. For the coupling reactions, 3 equiv of monomers in the presence of 4-15 equiv of 1,8-diazabicyclo[5.4.0]undec-7-ene (DBU) were used in DMI, which was the preferred solvent to compensate the low solubility of the growing oligomers. It is worth noting that the exclusion of water by azeotroping with toluene before the coupling reaction and by use of molecular sieves during the reaction was critical for successful coupling reactions. A mixture of triethylsilane, 1,1,1,3,3,3-hexafluoroisopropanol (HFIP), trifluoroethanol and dichloromethane was used for the mild deprotection of DMTr protecting group. Repeated precipitations after the coupling and deprotection reactions were employed to give the oligomeric intermediates in good purity.

The 5’-fragment was synthesized by coupling of a 5’-stereopure PMO wing with a 3’-tert-butyldimethylsilyl (TBS) protected DNA 5’-chlorophosphoramidate (Figure 2C). The resulting product was desilylated and activated with a selected PSI reagent to give the 5’-fragment for the block coupling. The key block coupling reaction was conducted by use of the 3’-fragment as the limiting reagent in the presence of 3 equiv of 5’-fragment and an excess amount of DBU in DMI at room temperature to give the coupling product in 30 – 80% yield. The obtained crude product was subjected to global deprotection with 28% NH_4_OH in methanol in the presence of DL-dithiothreitol at 50 – 55 °C. The resulting product was precipitated from the mixture by adding acetonitrile and ethyl acetate. The isolated solid was dissolved in water and filtered using Amicon ultracel 3K filter to remove unreacted substrate residues. Finally, the product was purified by reverse phase HPLC followed by ion-exchange chromatography, desalted with Amicon Ultracel 3K filter, and lyophilized to give a stereodefined PMO-gapmer.

The reaction progress was monitored by UPLC-MS, and ^31^P NMR was used to assess the chemical and stereochemical integrity of synthetic intermediates and stereodefined PMO-gapmers. The phosphorodiamidate and phosphorothioate linkages showed unique ^31^P NMR chemical shifts at 15-20 ppm and 55-60 ppm regions, respectively. Since the stereodefined PMO-gapmers have 7-9 phosphorodiamidate linkages and 8-10 phosphorothioates linkages, individual phosphorus peaks were well resolved. All the stereopure intermediates show different ^31^P NMR patterns from its stereoisomers. The stereodefined PMO-gapmers showed distinct ^31^P NMR and HPLC profiles to be used for identification. Under the HPLC conditions used for the analysis, certain stereodefined PMO-gapmers showed clear separation from the other stereoisomer having one different phosphorus stereochemistry (e.g., ASO-486-R5-S vs. ASO-486-R5-R, Supplementary Information).

### Selection of candidate sequences

Sequence screening for PMO-gapmers commenced with an *in silico* screening of oligonucleotides having 18-mer targeting both exonic and intronic regions of *MAPT* sequences homologous in cynomolgus monkey, rhesus monkey and human (Figure 3). The sequences having self-complementarity of more than 4 base pairs were filtered out. The identified sequences were further narrowed down by off-target search allowing up to 2 mismatches and by removing the sequences targeting disease causing and nerve system related genes by gene ontology (GO) terms.

**Figure 3.**
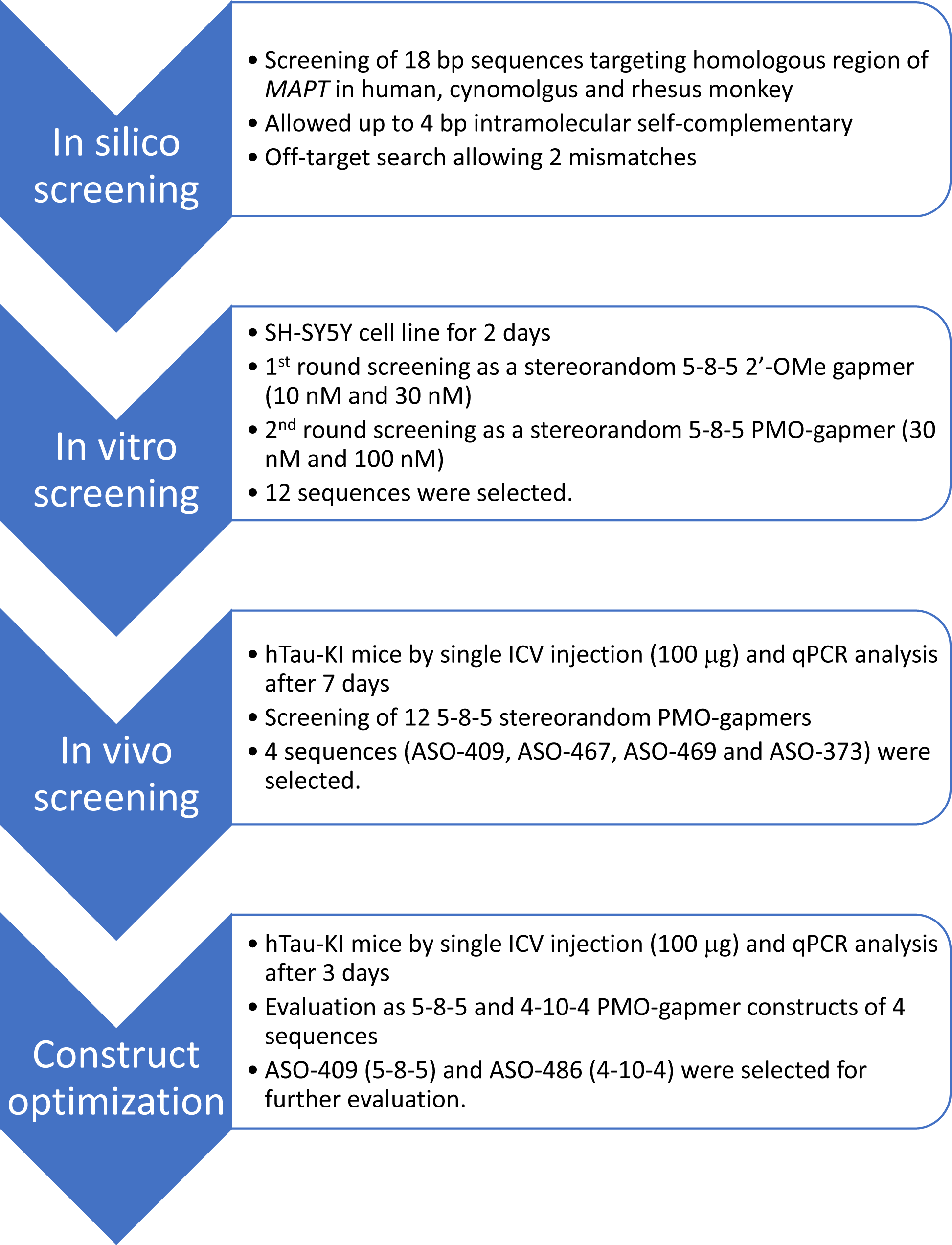
Schematic representation of screening flow.

Since a high throughput solid phase synthesis towards PMO-gapmers was not established yet, 5-8-5 2’-OMe gapmer was used as a surrogate of PMO-gapmer for the first-round sequence screening. The synthesized 2’-OMe gapmers were subjected to in vitro screening: human neuroblastoma (SH-SY5Y) cell line was treated with the gapmers in the presence of Lipofectamine^TM^ RANiMAX for 48 hr, and the knockdown results were analyzed by qPCR. The in vitro screening identified 65 active sequences after excluding the ones showing cytotoxicity. The sequences were prepared as a stereorandom 5-8-5 PMO-gapmer by solid phase synthesis and evaluated by in vitro assay using Endo-Porter as the trasfection agent (Supplementary Table S2 and Table S3). The 12 most active sequences were selected for in vivo evaluation in hTau knock-in mice (36) by single ICV injection. Four lead oligonucleotide sequences were identified by their prominent *MAPT* gene knockdown activity: ASO-409, ASO-467, ASO-469 and ASO-373 (Supplementary Figure S4). The 4 sequences were synthesized as two different PMO-gapmer configurations, 5-8-5 and 4-10-4 PMO-gapmers, and evaluated by in vivo study. Two most active compounds having similar sequences but different configurations, ASO-409 (5-8-5) and ASO-486 (4-10-4), were selected for further evaluations (Figure 4).

**Figure 4.**
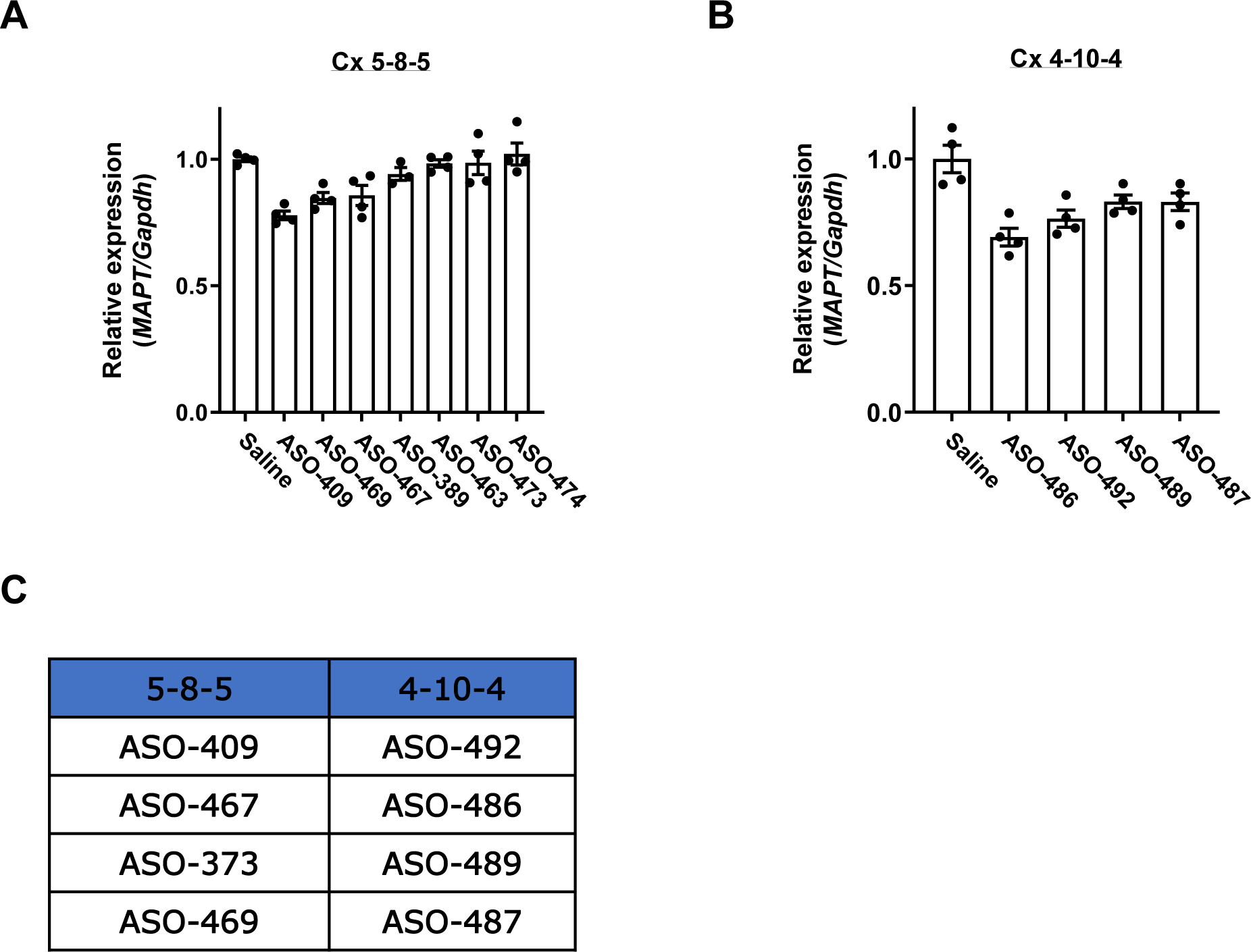
In vivo sequence screening of PMO-gapmer. (A and B) In vivo knockdown activities of stereorandom PMO-gapmers with 5-8-5 (A) and 4-10-4 (B) configuration. hTau KI mice were treated with 100 mg of ASO by ICV injection and *MAPT* mRNA in cortex was measured after 7 days. Error bars represent standard error of the mean. (C) Correlation of ASO sequence and configuration: e.g., ASO-409 and ASO-492 have the same sequence but different configuration of 5-8-5 and 4-10-4, respectively.

### Investigation of optimal phosphorus stereochemistry

Screening for optimal phosphorus stereochemical patterns was conducted with the selected two PMO-gapmer sequences (Figure 5 and Figure 6). The phosphorus stereochemistry at the PMO wing regions was fixed as *S*p to have better binding to the target sequences. Considering that *S*p phosphorothioate linkages show better stability against nucleases (40), more *S*p linkages with minimum number of *R*p linkages in the DNA gap region would be preferred to have longer duration of action. Thus, single *R*p walk in the DNA gap region was conducted to find the optimal position of the *R*p linkage in terms of knockdown activity. Also, *SSR* stereo-patterns at the gap region were evaluated since Verdine *et al.* reported that *SSR* stereotriad shows good activity (40).

**Figure 5.**
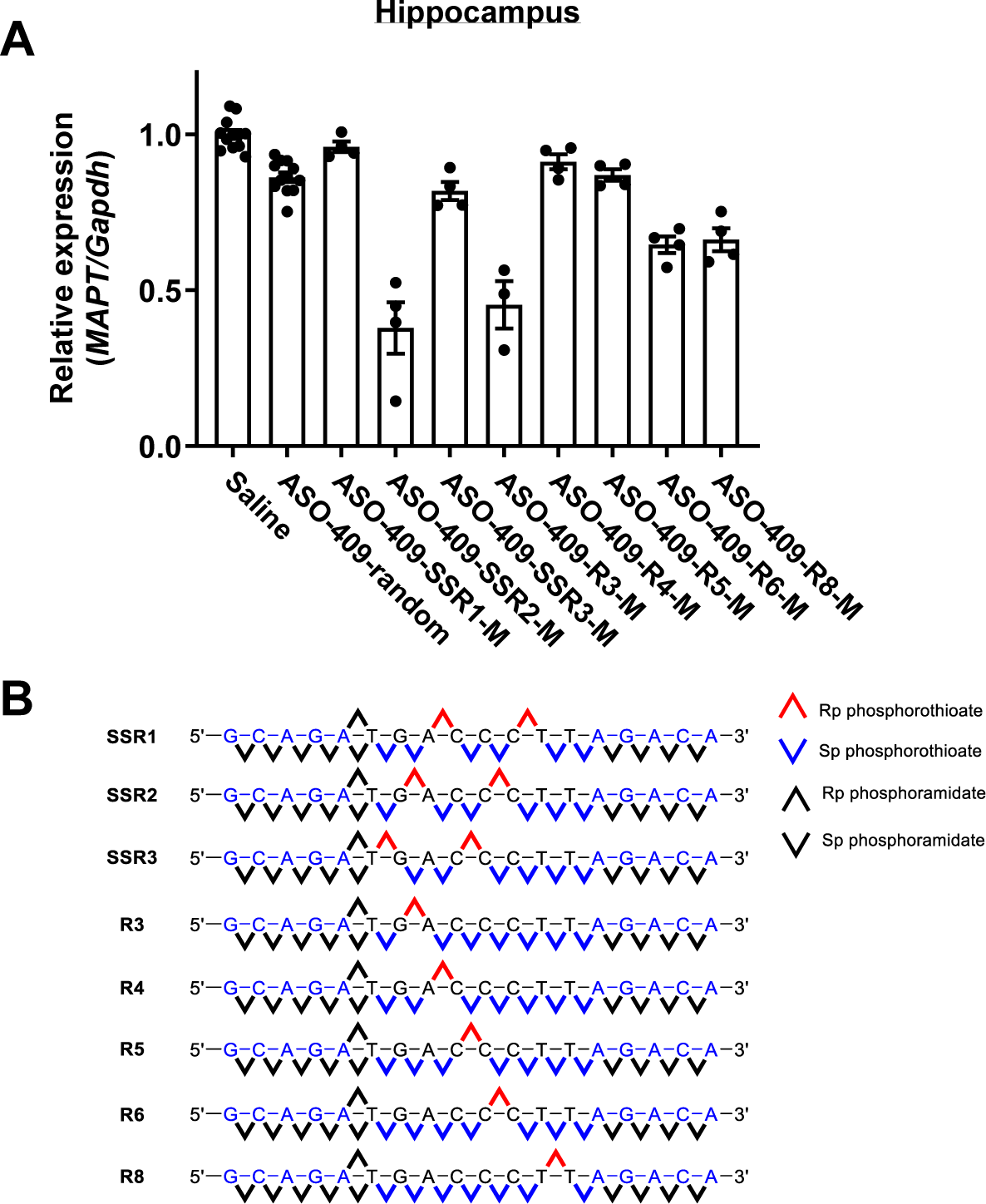
Screening of optimal stereochemistry at the DNA gap region for ASO-409. (A) Stereochemistry-dependent reduction of *MAPT* mRNA in hippocampus. The study was conducted in hTau KI mice by ICV injection of 100 mg dose. *MAPT* mRNA was measured after 3 days. Error bars represent standard error of the mean. (B) Schematic representation of chemical configuration and phosphorus stereochemistry of ASO-409 stereoisomers.

**Figure 6.**
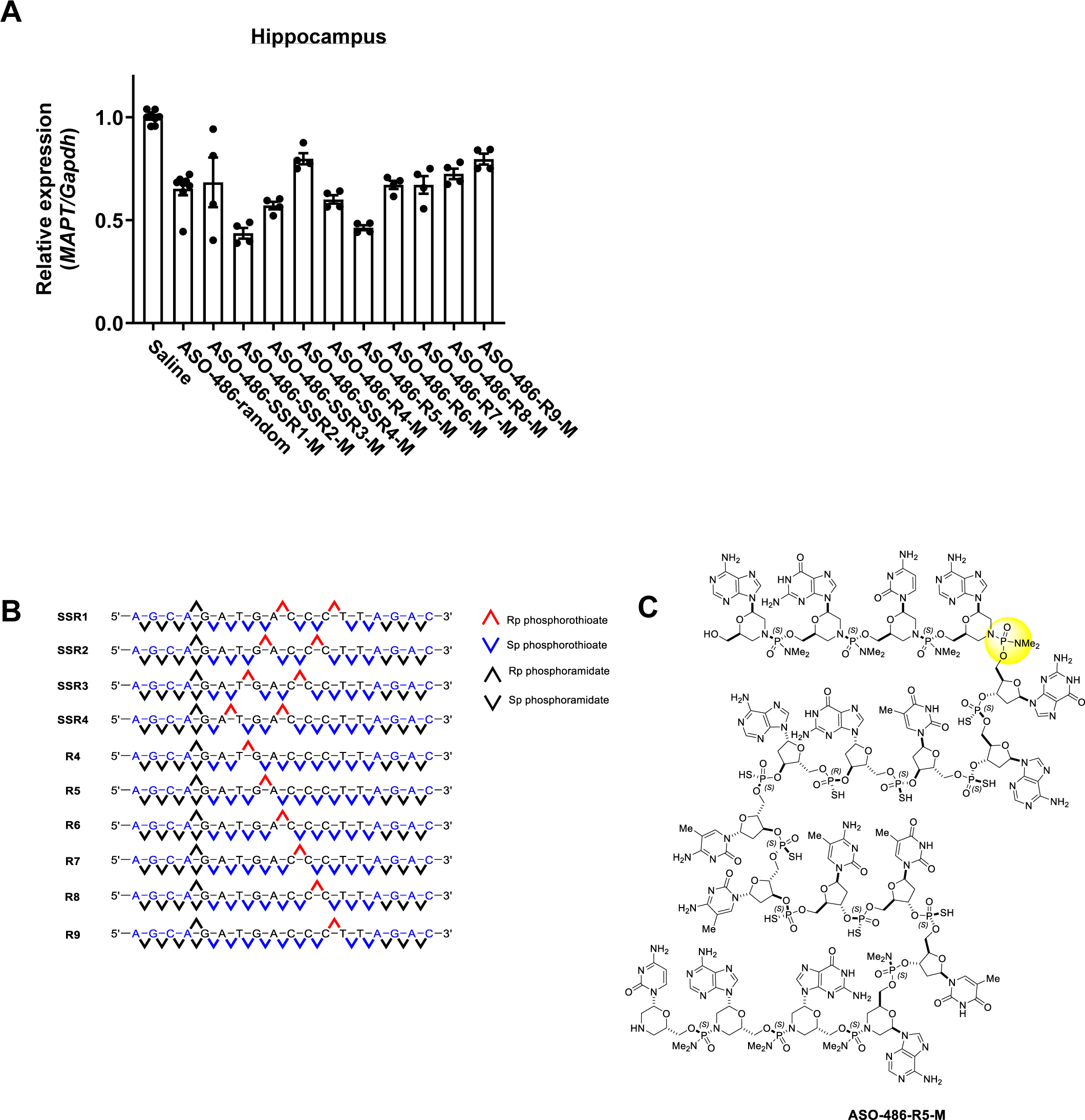
Screening of optimal stereochemistry at the DNA gap region for ASO-486. (A) Stereochemistry-dependent reduction of *MAPT* mRNA in hippocampus. The study was conducted in hTau KI mice by ICV injection of 100 mg dose. *MAPT* mRNA was measured after 3 days. Error bars represent standard error of the mean. (B)Schematic representation of chemical configuration and phosphorus stereochemistry of ASO-486 stereoisomers. (C) Chemical structure of stereopure ASO-486 was represented by ASO-486-R5-M. Phosphorus chirality at the 4^th^ position (yellow circle) was not controlled for this study.

For ASO-409, 3 PMO-gapmers having *SSR* patterns and 5 compounds having one *R*p phosphorothioate linkage in the DNA gap region were synthesized and evaluated by 3 day in vivo study with hTau KI mice (Figure 5). For synthetic convenience, the phosphorus stereochemistry of the phosphorodiamidate linkage at the 5’ PMO-DNA junction was not controlled and a mixture of two stereoisomers was used for in vivo study. The stereoisomers showed quite different in vivo activities: ASO-409-SSR2-M showed the best activity, which was ca. 5 fold better than that of stereorandom ASO-409. ASO-409-R3-M isomer showed similar activity with ASO-409-SSR2-M suggesting the Rp linkage at the position between 7th and 8th nucleotide (G-A) is important for the activity.

Similar stereochemical structure activity relationship (SSAR) studies were conducted with the ASO-486, a 4-10-4 PMO-gapmer sequence by synthesizing and characterizing different stereoisomers (Figure 6). Since ASO-486 has 10 nucleotides in the gap region, 4 PMO-gapmers having *SSR* patterns were designed and *R*p walk was conducted using 6 compounds having one *R*p linkage. Among them, ASO-486-SSR2-M showed the best activity, and ASO-486-R5-M showed similar activity with ASO-486-SSR2-M suggesting that the *R*p linkage at the position between 8th and 9th nucleotide (G-A) is important for the activity. The relative activity difference between ASO-486-SSR2-M/ASO-486-R5-M and ASO-486-random was smaller than that observed in ASO-409. The results suggested that the impact of stereochemistry would depend on the ASO sequences and ASO configuration.

### Impact of phosphorus stereochemistry on safety

The two most potent ASO-409 stereoisomers (ASO-409-SSR2-M and ASO-409-R3-M) from 3 day in vivo study were subjected to 2-month in vivo study with hTau KI mice at 30 and 60 μg dose to evaluate the duration of knockdown activity and preliminary safety after single ICV administration (4 animals/group). Mortality occurred in 3 animals in 60 μg ASO-409-SSR2-M group after 3 weeks (Figure 7A). One motality in 30 μg ASO-409-R3-M group was suspected to be incidental since all the other animals in the same group did not show any sign of toxicity. The mortality was not caused by the *MAPT* knockdown: ASO-409-R8-M group did not show any toxicity including mortality and body weight loss although ASO-409-R8-M showed similar *MAPT* knockdown activity with that of ASO-409-SSR2-M after 3 weeks (Figures 7B and 7C). To assess impact of the uncontrolled phosphorus chirality at the phosphorodiamidate linkage at 5’-PMO and DNA junction on the safety profile, fully stereocontrolled ASO-409-SSR2 and ASO-409-R3 were synthesized and subjected to 2-month in vivo single ICV dosing study at 60 μg dose (Figure 8). In this study, ASO-409-SSR2-M group showed similar results with the previous study. All the mice in the ASO-409-SSR2-S group also showed motality after 3 weeks, but only one mouse showed motality from ASO-409-SSR2-R group which differs in one phosphorus chirality. The single phosphorus stereochemistry exhibited more dramatic differences in safety profile in ASO-409-R3 groups. All the mice in ASO-409-R3-S group died in 6 weeks, but the mice in ASO-409-R3-R group did not show any mortality or body weight loss in 8 weeks.

**Figure 7.**
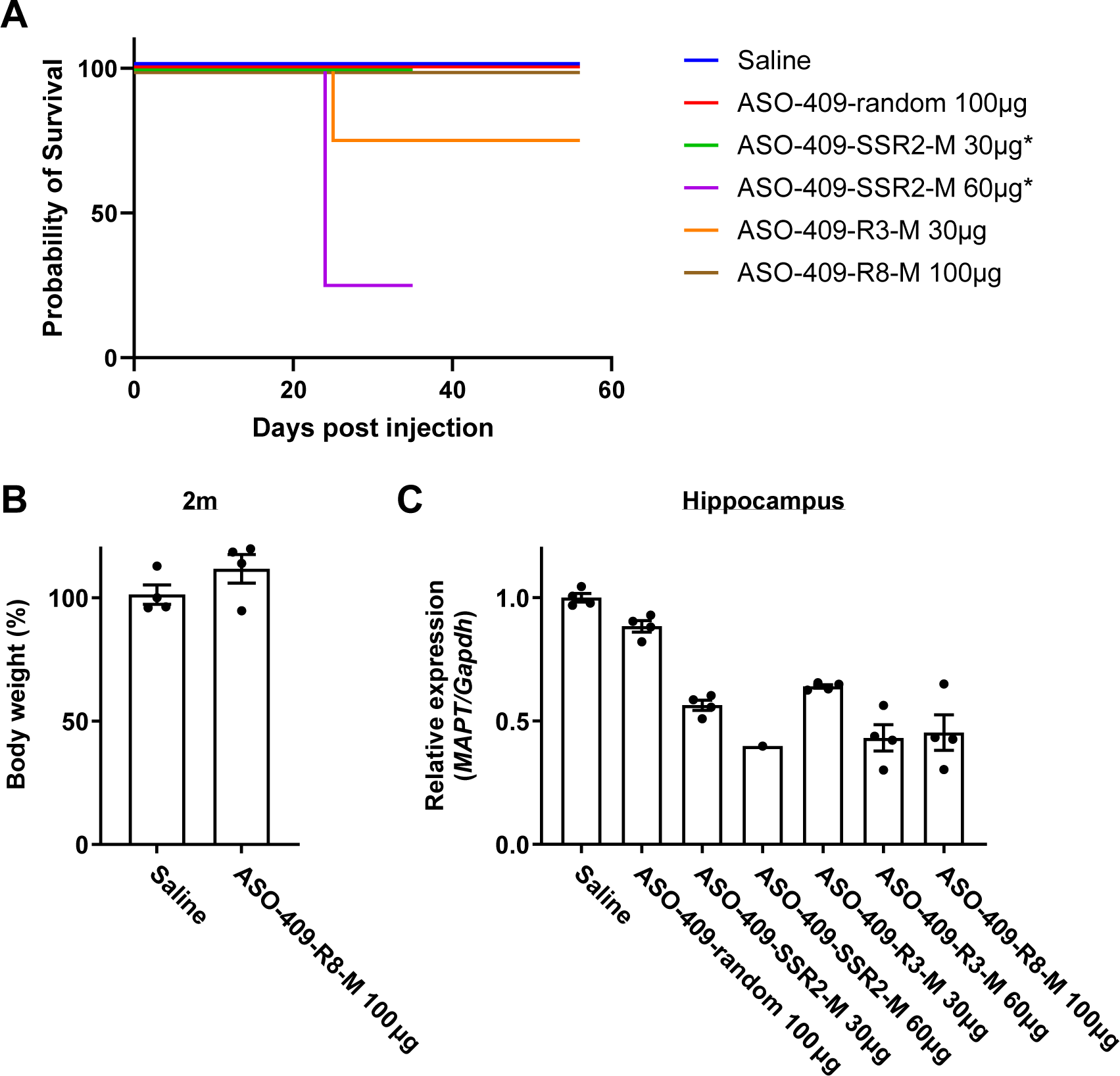
Assessment of long-term in vivo tolerability and knockdown activity for selected ASO-409 stereoisomers with hTau KI mice. (A) Survival curve after treatment with ASO-409-SSR2-M, ASO-409-R3-M, and ASO-409-R8-M. Asterisks represent the groups sacrificed at 35 days after injection due to moribund condition. (B) Body weight change 2 months after the treatment with ASO-409-R8-M. (C) In vivo knockdown of *MAPT* mRNA 3 weeks after treatment with ASO-409-SSR2-M, ASO-409-R3-M and ASO-409-R8-M. Error bars represent standard error of the mean.

**Figure 8.**
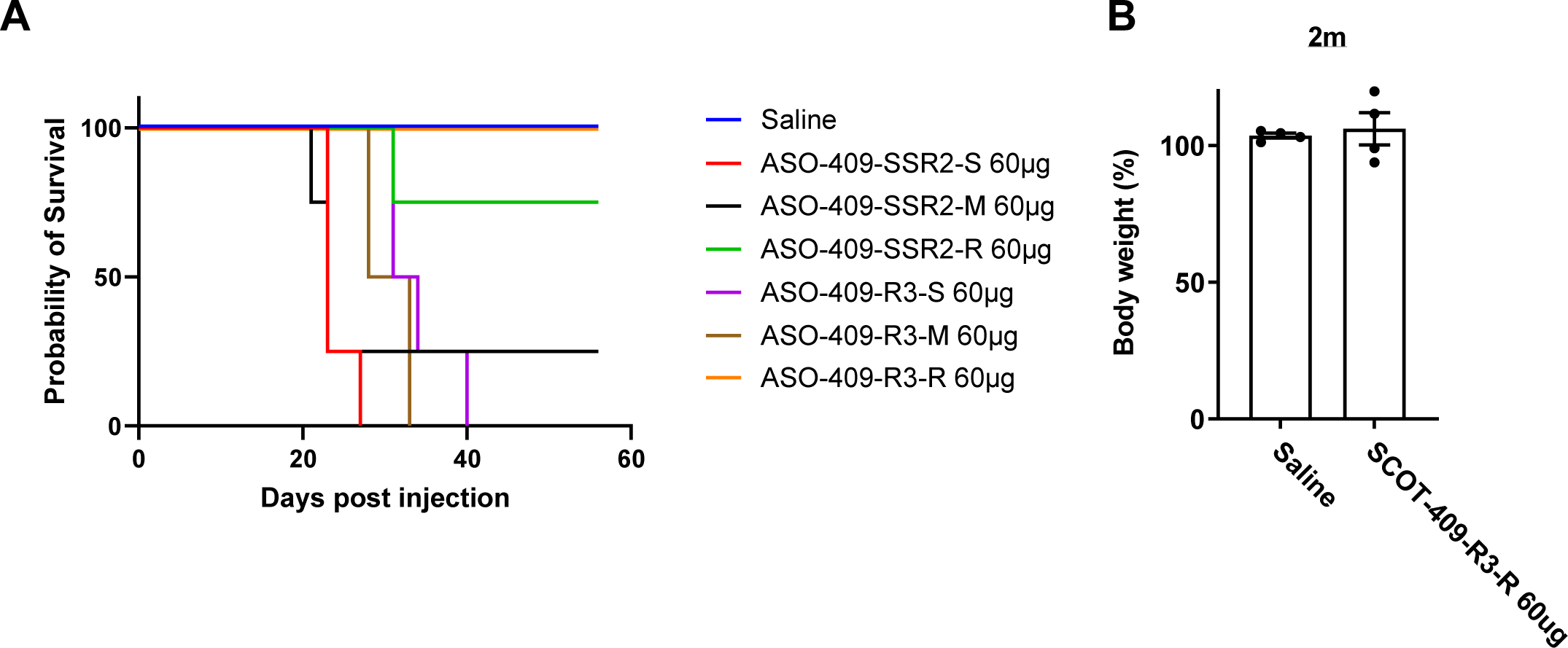
Assessment of long-term in vivo tolerability for completely stereodefined ASO-409-SSR2 and ASO-409-R3 with hTau KI mice. (A) Survival curve after treatment with completely stereodefined ASO-409-SSR2 and ASO-409-R3. (B) Body weight change 2 months after treatment with ASO-409-R3-R. Error bars represent standard error of the mean.

### Identification of ASO-486-R5 as a candidate

The two most potent stereoisomers of ASO-486 (ASO-486-SSR2-M and ASO-486-R5-M) from 3 day in vivo study was subjected to 2-month in vivo study with hTau KI mice at 30 μg and 60 μg dose to evaluate duration of knockdown activity and potential toxicity after single ICV administration (4 animals / group). ASO-486-R4-M, which showed weaker activity at 3 day in vivo study, as well as a stereorandom ASO-486 were tested in parallel at 100 μg dose for comparison (Figure 9). Gratifyingly, no mortality was observed after ASO-486 treatment. Furthermore, all the tested groups except ASO-486-SSR2-M 60 μg dose group did not show any body weight loss (Supplementary Figure S5). Both ASO-486-SSR2-M and ASO-486-R5-M showed dose dependent *MAPT* mRNA reduction lasting for 2 months. ASO-486-R5-M showed ca. 40% knockdown in cortex after 3 weeks, which was weaker than that of ASO-486-SSR2-M (ca. 60% knockdown). However, ASO-486-R5-M showed smaller decrease in the knockdown activity over time to show similar knockdown activity with ASO-486-SSR2-M after 2 months.

**Figure 9.**
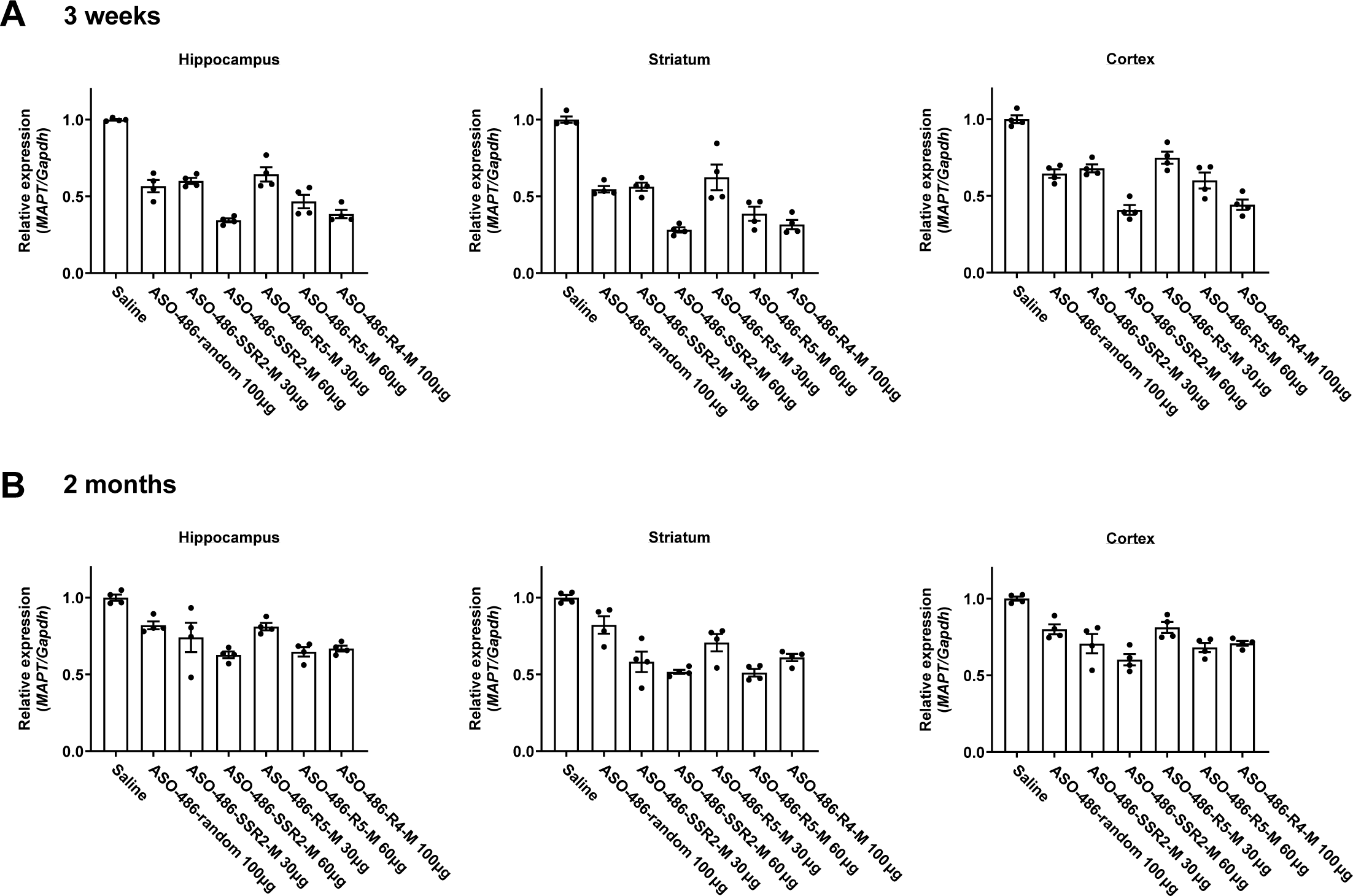
Assessment of long-term in vivo knockdown activity of ASO-486 stereoisomers with hTau KI mice. (A and B) Stereochemistry-dependent knockdown of *MAPT* mRNA in hippocampus, striatum, and cortex 3 weeks (A) and 2 months (B) after ASO treatment. Error bars represent standard error of the mean.

The stability of ASO-486 stereoisomers were evaluated in human brain homogenates (Figure 10). The residual percentage of stereorandom ASO-486 decreased time-dependently and was about 50% after 6 days incubation. As expected, all the stereodefined PMO-gapmers showed better stability profile than the stereorandom compound with up to 20% difference among different stereoisomers. ASO-486-R9-M showed the most noticeable degradation among stereodefined ASO-486 isomers (20% decrease), which was, however, still significantly better than that of the stereorandom compound. The compounds having one *R*p and two *R*p linkages (*SSR* isomers) did not show significant stability differences. The results suggested that the stability of PMO-gapmers would depend on not only the number of *R*p linkages but also the position of *R*p linkages.

**Figure 10.**
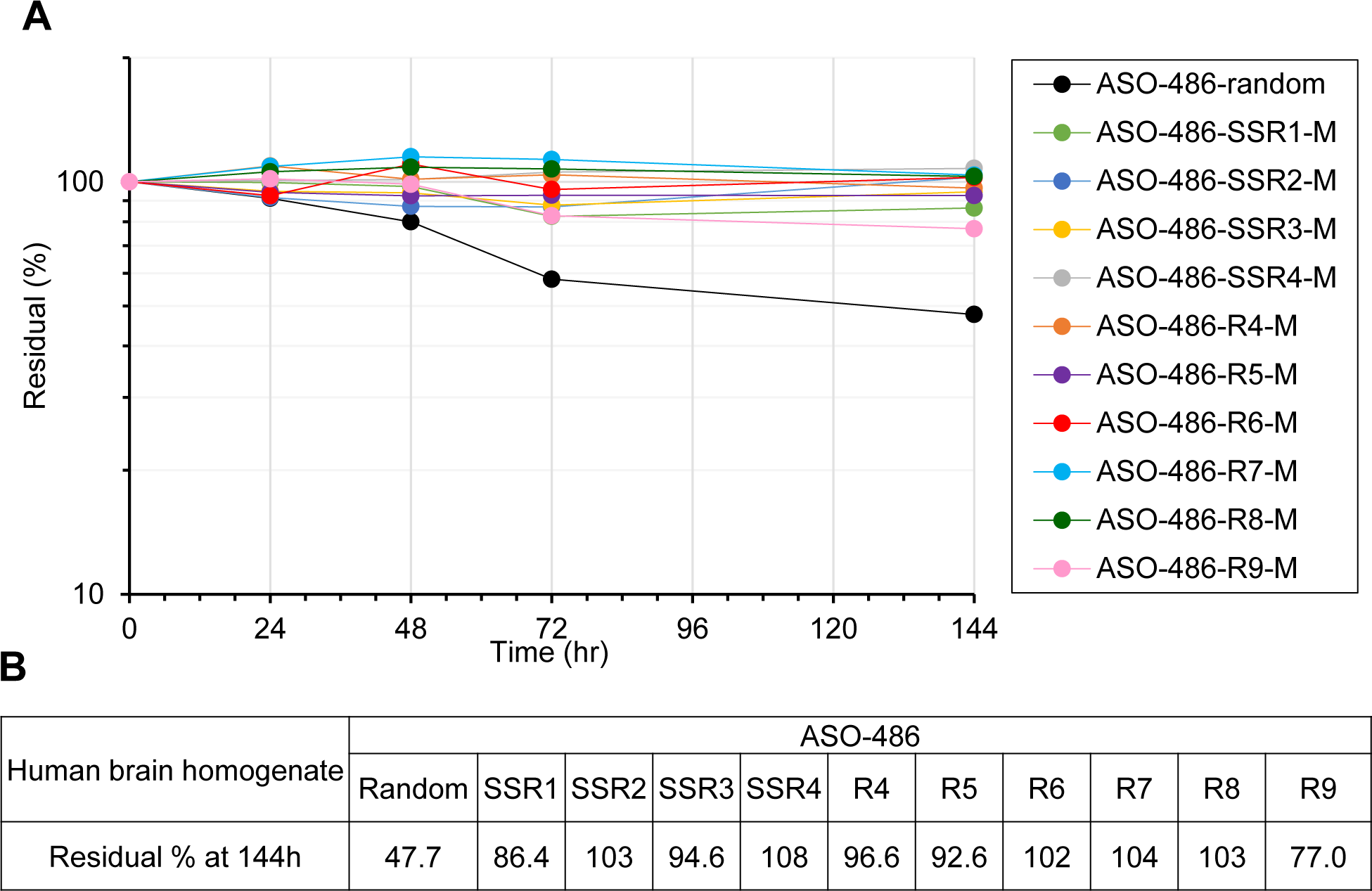
In vitro stability of stereo-controlled ASO-486 in human brain homogenates. Residual percentages over time (A) and at 144 h (B) for stereo-random and stereodefined ASO-486 in human brain homogenate. Each ASO at 1 µM was incubated in gender mixed human brain homogenate at 10 mg/mL for 144 h. The samples after the pretreatment were analyzed by ion-pair liquid chromatography coupled with mass spectrometry. The residual percentages were calculated by dividing the peak area ratios at 24, 48, 72, and 144 h by that at 0 h. The high residual percentages after 144 h incubation were shown in stereodefined ASO-486 groups as compared with stereorandom ASO-486.

For the evaluation of individual stereoisomers, the stereopure isomers of ASO-486-SSR2-M and ASO-486-R5-M (ASO-486-SSR2-S, ASO-486-SSR2-R, ASO-486-R5-S and ASO-486-R5-R) were synthesized. All the stereoisomers showed good knockdown activity in 3 day in vivo study with a tendency of junction *S*p isomers showing slightly better activity than the corresponding *R*p isomers (Figure 11). In 2-month in vivo study, the stereopure ASO-486-SSR2-S and ASO-486-SSR2-R exhibited around 10% body weight loss at 60 μg dose (Figure 12B). In addition, ASO-486-SSR2-S showed increased cytokine release (IL-6, IL-8, IL-13 and IFN-γ) at high doses in human PBMCs (Supplementary Figure S6). On the other hand, ASO-486-SSR2-R did not show any changes in cytokine release providing another evidence that a single phosphorus stereochemistry change has impact on safety profile. Gratifyingly, ASO-486-R5-S and ASO-486-R5-R did not show any body weight changes in 2-month in vivo study or increases in cytokine release in human PBMCs. Based on the results, ASO-486-R5-S and ASO-486-R5-R were selected as candidates for further development.

**Figure 11.**
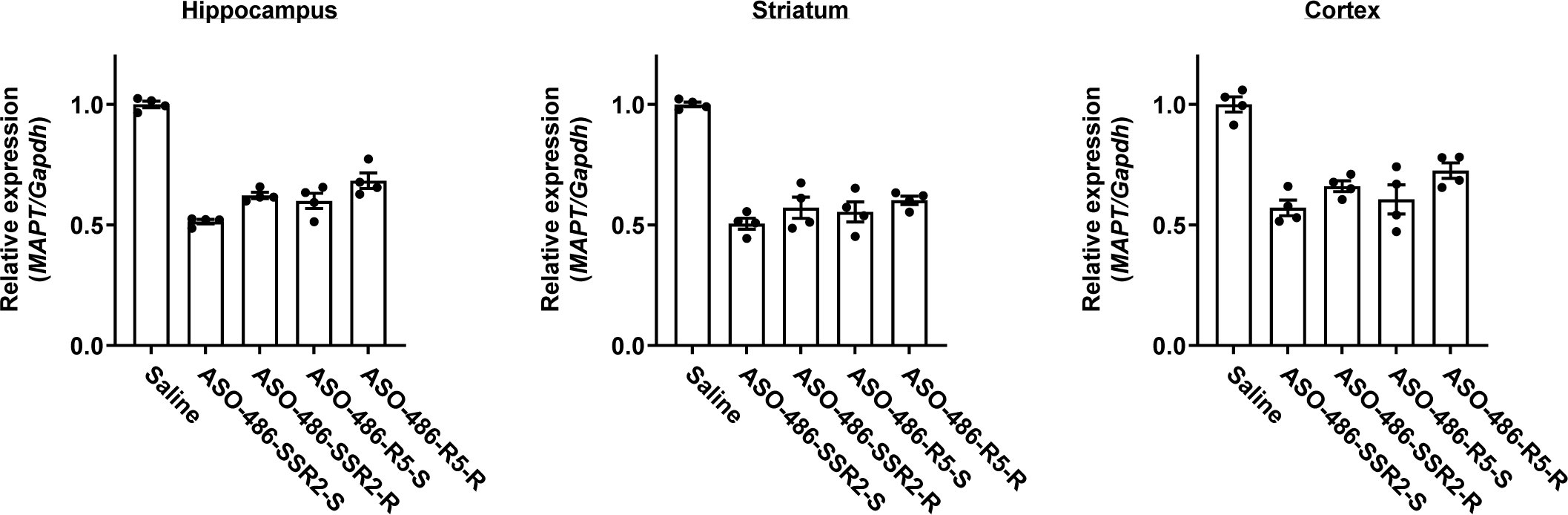
In vivo knockdown activity of completely stereodefined ASO-486-SSR2 and ASO-486-R5 with hTau KI mice. hTau KI mice were treated with 60 mg of ASO by ICV injection. *MAPT* mRNA in hippocampus, striatum, and cortex were measured after 3 days. Error bars represent standard error of the mean.

**Figure 12.**
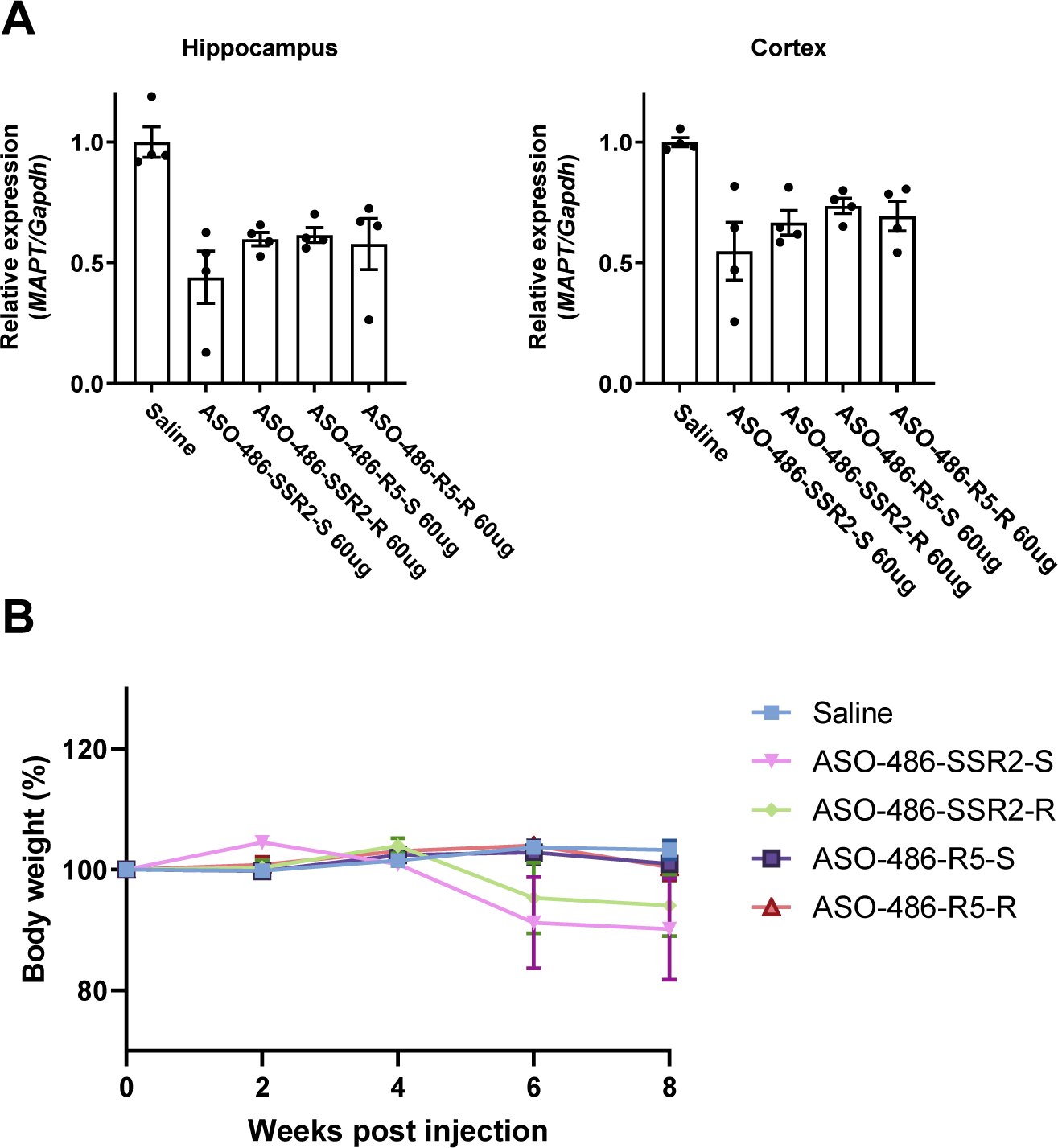
Long-term (8 weeks) *in vivo* study of completely stereodefined ASO-486-SSR2 and ASO-486-R5 with hTau KI mice. (A) Knockdown of *MAPT* mRNA 2 months after treatment with completely stereodefined ASO-486-SSR2 and ASO-486-R5 in hippocampus and cortex. Error bars represent standard error of the mean. (B) Body weight changes after treatment with ASO.

## DISCUSSION

ASOs are an important therapeutic modality for the selective reduction of target genes. The clinical application of ASOs have successfully expanded to central nervous system (CNS) diseases to treat spinal muscular atrophy (SMA, nusinersen) and amyotrophic lateral sclerosis (ALS, tofersen) by intrathecal injection. To further expand the application of ASOs in CNS field, improved chemical modifications would be desirable to increase the activity, safety profile and stability to lengthen the dosing interval after intrathecal injection for patients’ convenience. Since the stabilization strategies for ASOs have replaced the oxygen(s) at the phosphorous internucleotide linkages with other atoms, the issue of stereocontrol has emerged. Thus far, stereorandom ASO modalities consisting of mixtures of thousands to billions of individual components have been evaluated pre-clinically and clinically, with several commercial products as well. Recent publications have highlighted several serious issues with stereorandom ASOs: 1) analytical techniques for traditional small molecules are incapable of detecting, let alone quantifying single components in a complex stereorandom ASO mixture, 2) batch-to-batch consistency is not possible for producing complex mixtures of stereoisomers, 3) biological assessments of different batches of the “same” stereorandom ASO reveal significant differences in behavior at the pre-clinical and clinical stage (28, 29, 30). In order to investigate the full potential of the ASO modality, it was thus desirable to apply small molecule standards (i.e., single well-characterized molecule with well-defined and consistent biological properties) to this therapeutic platform. Whereas several technologies have recently been developed to assemble phosphorothioate linked oligonucleotides in a stereocontrolled manner, thus far no method has been reported to assemble phosphorodiamidate linked PMOs with any control of stereochemistry at the internucleotide linkage. Recently, stereopure PMO monomers were reported from our labs that enable precise control of stereochemistry at each stereogenic phosphorous atom. With a view to create single-molecule tau-targeting ASOs with well-defined quality, safety, and efficacy, a goal was set to combine the stereodefined PMO and phosphorothioate methodologies to assemble a chimeric ASO (i.e., PMO-gapmers).

For the evaluation of therapeutic application of PMO-gapmers, a solid phase synthesis of stereorandom PMO-gapmers for sequence screening and a convergent solution-phase synthetic method for stereodefined PMO-gapmers were developed. The solution phase synthesis for stereodefined PMO-gapmers utilizes a block coupling of 3’-fragment with PSI-activated 5’-fragment. The synthetic protocol consistently gives the desired product in good yield, and could be the basis for an efficient method for manufacturing clinical candidates (41). Considering the low throughput of the solution phase method, development of a high throughput solid-phase synthesis is desirable for the screening of stereodefined PMO-gapmers, and the progress will be reported in due course.

With optimal sequences for *MAPT* knockdown, searching for optimal phosphorus stereochemistry was conducted with the stereoisomers selected based on hypotheses in ASO-409 or ASO-486 (2^17^= 131,072 possible stereoisomers). For the initial screening of stereochemistry, a mixture of two stereoisomers without control of the phosphorus stereochemistry at the 5’-junciton of PMO and DNA was used for synthetic convenience. The stereochemistry of the phosphorodiamidate linkages in PMO wings was fixed as *S*p to have better binding to the target sequences. Considering that the stereorandom PMO-gapmer has lower Tm than the corresponding MOE-gapmer, *S*p linkages at the wing region were expected to partially compensate for the differences in Tm and achieve optimal binding affinity. For the selection of the stereochemistry for the DNA gap region, it was desirable to have more *S*p linkages than *R*p to have better stability against endonucleases (40), which would translate to longer duration in in vivo study. Thus, single *R*p walk and *SSR* walk in the DNA gap region were conducted to find the stereoisomers showing the best activity. Similar strategies have been reported for the identification of the optimal stereochemistry (42, 43). After evaluating the activity of all the synthesized stereoisomers, it was found that *R*p at specific positions (e.g., at G7-A8 for ASO-409 (Figure 5) and at G8-A9 for ASO-486 (Figure 6)) is crucial to have good knockdown activity. In addition, the presence of a second *R*p linkage did not give significant activity gain (e.g., ASO-409-SSR2-M vs. ASO-409-R3-M, and ASO-486-SSR2-M vs. ASO-486-R5-M). It is reported that the position of *R*p linkage in the gap region guides the RNase H cleavage site and selection of the position of *R*p linkage reinforcing the cleavage at the native cleavage sites gives a cleaner cleavage profile and higher knockdown activity (42, 43). The RNase H cleavage site analysis showed that ASO-486-SSR2-R and ASO-486-R5-R result in an increased cleavage at the same cleavage site presumably due to the *R*p linkage at G8-A9 (Supplementary Figure S7). ASO-486-SSR2-R showed additional cleavage at the secondary cleavage site due to the additional *R*p linkage at C11-C12. However, the cleavage at the secondary site was very weak and did not change overall knockdown activity.

Stereochemical structure activity relationship (SSAR) studies revealed remarkably different safety profiles for two stereodefined PMO-gapmers differing at a *single* phosphorus stereogenic center. In 2-month in vivo study to evaluate the duration of activity and safety profile, ASO-409-SSR2-M showed mortality after 3 weeks (no acute adverse event was observed after ICV injection). The quality check of study samples confirmed that the endotoxin level and osmolarity of the samples were in the acceptable range (44, 45). The stereopure components of ASO-409-SSR2-M and ASO-409-R3-M were synthesized and subjected to the 2-month study at 60 μg dose to unambiguously evaluate the safety profile of each component. ASO-409-SSR2-S and ASO-409-R3-S showed similar toxicity with the corresponding mixture and more toxic results than ASO-409-SSR2-R and ASO-409-R3-R, respectively. Surprisingly, ASO-409-R3-R did not show any toxicity or weight changes after 2 months although ASO-409-R3-S showed total mortality. These results strengthen the hypothesis that rigorous investigation of single stereodefined ASOs with control of phosphorus stereochemistry will be essential to select safe ASOs within a given sequence.

The mice treated with ASO-409-SSR2-M showed mortality, forelimb dysfunction or abnormal gait, and body weight loss in 3-week and 2-month in vivo study. In histopathologic examination, degeneration and/or necrosis of Purkinje cells with vacuolation in the pons and spinal ventibular nucleus was observed in all animals treated with ASO-409-SSR2-M at 30 μg and above. However, ASO-409-SSR2-S and ASO-409-R3-S showed similar target knockdown activity at cerebellum and cortex (Supplementary Figure S8). Taken together, the results ruled out the possibility that disproportionate ASO distribution in different brain region would have caused the toxicity. The ASOs having the same sequence and chemistry but different stereochemistry did not show any toxicity (e.g., ASO-409-R8-M in Figure 7). Thus, it was proposed that the observed toxicity is not due to the ASO modality but potentially due to knockdown of off-target genes. In fact, *Atp2b2* with 1-base mismatch was identified as one of potential causal genes using RNA-sequence method and *in silico* mouse off-target search (data not shown). Considering that the target sequence in mouse *Atp2b2* gene does not overlap with human gene sequence, the observed adverse event might be limited to mice.

Towards identification of development candidates, ASO-486-R5 showed good activity and drug properties. ASO-486-R5-M showed good knockdown activity lasting over 2 months. Even though ASO-486-R5-M showed slightly weaker activity than ASO-486-SSR2-M in 3 weeks, it showed similar activity after 8 weeks without any body weight loss. The corresponding fully stereocontrolled ASOs, ASO-486-R5-S and ASO-486-R5-R, showed almost the same binding affinity to the target sequence measured by Tm (60.7 °C and 60.5 °C, respectively), similar knockdown activity in 3 day in vivo study (Figure 11) and no body weight change in 2-month in vivo study at 60 μg dose. Since the stereo-patterns of ASO-486 were selected to have better stability, all the stereoisomers including ASO-486-R5-M showed better stability than the corresponding stereorandom mixtures in human brain homogenates. In addition, ASO-486-R5-S and ASO-486-R5-R did not show any increase in cytokine release in human PBMCs. Further studies in mice and non-human primates will be conducted for the evaluation of both compounds towards clinical development.

In conclusion, a novel PMO-gapmer was discovered and evaluated as a potentially safer and more stable gapmer construct. Methods were also described that enable convergent synthesis of stereodefined PMO-gapmers and the stereochemical structure activity studies (SSAR) employing the PMO-gapmers. SSAR studies revealed remarkable differences in the safety and toxicity of stereodefined PMO-gapmers differing at a *single* phosphorus atom stereogenic center. After series of *in silico*, in vitro and in vivo screening, ASO-486-R5-S and ASO-486-R5-R were identified as potential development candidates targeting *MAPT* for the treatment of tauopathies. Further studies in mice and NHP are warranted to evaluate the safety, activity and duration of ASO-486-R5-S and ASO-486-R5-R.

## SUPPLEMENTARY DATA

Supplementary Data are available at NAR online.

## ACKNOWLEDGEMENT

The authors would like to thank prof. Phil Baran for valuable discussion and suggestions, and Yu Chen, Ph.D. and Alexander Amatuni, Ph.D. for technical review and proofreading of the manuscript.

## REFERENCES

1. Crooke, S.T., Liang, X., Baker, B.F. and Crooke, R.M. (2021) Antisense Technology: A Review. Journal of Biological Chemistry, 296, 100416.

2. Tran, H., Moazami, M.P., Yang, H., McKenna-Yasek, D., Douthwright, C.L., Pinto, C., Metterville, J., Shin, M., Sanil, N., Dooley, C., et al. (2021) Suppression of mutant C9orf72 expression by a potent mixed backbone antisense oligonucleotide. Nat Med, 28, 117–124.

3. Migawa, M.T., Shen, W., Wan, W.B., Vasquez, G., Oestergaard, M.E., Low, A., De Hoyos, C.L., Gupta, R., Murray, S., Tanowitz, M., et al. (2019) Site-specific replacement of phosphorothioate with alkyl phosphonate linkages enhances the therapeutic profile of gapmer ASOs by modulating interactions with cellular proteins. Nucleic Acids Res, 47, 5465–5479.

4. Shen, W., Hoyos, C.L.D., Migawa, M.T., Vickers, T.A., Sun, H., Low, A., Bell, T.A., Rahdar, M., Mukhopadhyay, S., Hart, C.E., et al. (2019) Chemical modification of PS-ASO therapeutics reduces cellular protein-binding and improves the therapeutic index. Nature Biotechnology, 37, 640–650.

5. Yoshida, T., Morihiro, K., Naito, Y., Mikami, A., Kasahara, Y., Inoue, T. and Obika, S. (2022) Identification of nucleobase chemical modifications that reduce the hepatotoxicity of gapmer antisense oligonucleotides. Nucleic Acids Res, 50, 7224–7234.

6. Zhang, L., Liang, X., Hoyos, C.L.D., Migawa, M., Nichols, J.G., Freestone, G., Tian, J., Seth, P.P. and Crooke, S.T. (2022) The Combination of Mesyl-Phosphoramidate Inter-Nucleotide Linkages and 2′-O-Methyl in Selected Positions in the Antisense Oligonucleotide Enhances the Performance of RNaseH1 Active PS-ASOs. Nucleic Acid Ther, 32, 401–411.

7. Vasquez, G., Migawa, M.T., Wan, W.B., Low, A., Tanowitz, M., Swayze, E.E. and Seth, P.P. (2022) Evaluation of Phosphorus and Non-Phosphorus Neutral Oligonucleotide Backbones for Enhancing Therapeutic Index of Gapmer Antisense Oligonucleotides. Nucleic Acid Ther, 32, 40–50.

8. Kandasamy, P., Liu, Y., Aduda, V., Akare, S., Alam, R., Andreucci, A., Boulay, D., Bowman, K., Byrne, M., Cannon, M., et al. (2022) Impact of guanidine-containing backbone linkages on stereopure antisense oligonucleotides in the CNS. Nucleic Acids Res, 50, 5401–5423.

9. Vasquez, G., Migawa, M.T., Wan, W.B., Low, A., Tanowitz, M., Swayze, E.E. and Seth, P.P. (2022) Evaluation of Phosphorus and Non-Phosphorus Neutral Oligonucleotide Backbones for Enhancing Therapeutic Index of Gapmer Antisense Oligonucleotides. Nucleic Acid Ther, 32, 40–50.

10. Sheng, L., Rigo, F., Bennett, C.F., Krainer, A.R. and Hua, Y. (2020) Comparison of the efficacy of MOE and PMO modifications of systemic antisense oligonucleotides in a severe SMA mouse model. Nucleic Acids Research, 48, 2853–2865.

11. Wan, W.B., Migawa, M.T., Vasquez, G., Murray, H.M., Nichols, J.G., Gaus, H., Berdeja, A., Lee, S., Hart, C.E., Lima, W.F., et al. (2014) Synthesis, biophysical properties and biological activity of second generation antisense oligonucleotides containing chiral phosphorothioate linkages. Nucleic Acids Res, 42, 13456–13468.

12. Funder, E.D., Albæk, N., Moisan, A., Sewing, S. and Koch, T. (2020) Refining LNA safety profile by controlling phosphorothioate stereochemistry. Plos One, 15, e0232603.

13. Østergaard, M.E., De Hoyos, C.L., Wan, W.B., Shen, W., Low, A., Berdeja, A., Vasquez, G., Murray, S., Migawa, M.T., Liang, X., et al. (2020) Understanding the effect of controlling phosphorothioate chirality in the DNA gap on the potency and safety of gapmer antisense oligonucleotides. Nucleic Acids Res, 48, 1691–1700.

14. Iwamoto, N., Butler, D.C.D., Svrzikapa, N., Mohapatra, S., Zlatev, I., Sah, D.W.Y., Meena, Standley, S.M., Lu, G., Apponi, L.H., et al. (2017) Control of phosphorothioate stereochemistry substantially increases the efficacy of antisense oligonucleotides. Nature Biotechnology, 35, 845–851.

15. Byrne, M., Vathipadiekal, V., Apponi, L., Iwamoto, N., Kandasamy, P., Longo, K., Liu, F., Looby, R., Norwood, L., Shah, A., et al. (2021) Stereochemistry Enhances Potency, Efficacy, and Durability of Malat1 Antisense Oligonucleotides In Vitro and In Vivo in Multiple Species. Translational Vision Science & Technology, 10, 23.

16. Hagedorn, P.H., Persson, R., Funder, E.D., Albæk, N., Diemer, S.L., Hansen, D.J., Møller, M.R., Papargyri, N., Christiansen, H., Hansen, B.R., et al. (2018) Locked nucleic acid: modality, diversity, and drug discovery. Drug Discovery Today, 23, 101–114.

17. Funder, E.D., Albæk, N., Moisan, A., Sewing, S. and Koch, T. (2020) Refining LNA safety profile by controlling phosphorothioate stereochemistry. Plos One, 15, e0232603.

18. Sierzchała, A., Okruszek, A. and Stec, W.J. (1996) Oxathiaphospholane Method of Stereocontrolled Synthesis of Diribonucleoside 3‘, 5‘-Phosphorothioates. J Org Chem, 61, 6713–6716.

19. Stec, W.J., Karwowski, B., Boczkowska, M., Guga, P., Koziołkiewicz, M., Sochacki, M., Wieczorek, M.W. and Błaszczyk, J. (1998) Deoxyribonucleoside 3‘-O-(2-Thio- and 2-Oxo-“spiro”-4, 4-pentamethylene-1, 3, 2-oxathiaphospholane)s: Monomers for Stereocontrolled Synthesis of Oligo(deoxyribonucleoside phosphorothioate)s and Chimeric PS/PO Oligonucleotides §. J. Am. Chem. Soc., 120, 7156–7167.

20. Karwowski, B., Okruszek, A., Wengel, J. and Stec, W.J. (2001) Stereocontrolled synthesis of LNA Dinucleoside phosphorothioate by the oxathiaphospholane approach. Bioorganic Medicinal Chem Lett, 11, 1001–1003.

21. Wilk, A., Grajkowski, A., Phillips, L.R. and Beaucage, S.L. (2000) Deoxyribonucleoside Cyclic N-Acylphosphoramidites as a New Class of Monomers for the Stereocontrolled Synthesis of Oligothymidylyl- and Oligodeoxycytidylyl-Phosphorothioates. J Am Chem Soc, 122, 2149–2156.

22. Iwamoto, N., Oka, N., Sato, T. and Wada, T. (2009) Stereocontrolled Solid-Phase Synthesis of Oligonucleoside H-Phosphonates by an Oxazaphospholidine Approach. Angewandte Chemie Int Ed, 48, 496–499.

23. Oka, N., Kondo, T., Fujiwara, S., Maizuru, Y. and Wada, T. (2009) Stereocontrolled Synthesis of Oligoribonucleoside Phosphorothioates by an Oxazaphospholidine Approach. Organic Letters, 11, 967–970.

24. Oka, N. and Wada, T. (2011) Stereocontrolled synthesis of oligonucleotide analogs containing chiral internucleotidic phosphorus atoms. Chem Soc Rev, 40, 5829–5843.

25. Knouse, K.W., deGruyter, J.N., Schmidt, M.A., Zheng, B., Vantourout, J.C., Kingston, C., Mercer, S.E., Mcdonald, I.M., Olson, R.E., Zhu, Y., et al. (2018) Unlocking P(V): Reagents for chiral phosphorothioate synthesis. Science, 361, eaau3369.

26. Knouse, K.W., Flood, D.T., Vantourout, J.C., Schmidt, M.A., Mcdonald, I.M., Eastgate, M.D. and Baran, P.S. (2021) Nature Chose Phosphates and Chemists Should Too: How Emerging P(V) Methods Can Augment Existing Strategies. Acs Central Sci, 7, 1473–1485.

27. Huang, Y., Knouse, K.W., Qiu, S., Hao, W., Padial, N.M., Vantourout, J.C., Zheng, B., Mercer, S.E., Lopez-Ogalla, J., Narayan, R., et al. (2021) A P(V) platform for oligonucleotide synthesis. Science, 373, 1265–1270.

28. Marafini, I., Stolfi, C., Troncone, E., Lolli, E., Onali, S., Paoluzi, O.A., Fantini, M.C., Biancone, L., Calabrese, E., Grazia, A.D., et al. (2021) A Pharmacological Batch of Mongersen that Downregulates Smad7 is Effective as Induction Therapy in Active Crohn’s Disease: A Phase II, Open-Label Study. Biodrugs, 35, 325–336.

29. Arrico, L., Stolfi, C., Marafini, I., Monteleone, G., Demartis, S., Bellinvia, S., Viti, F., McNulty, M., Cabani, I., Falezza, A., et al. (2022) Inhomogeneous Diastereomeric Composition of Mongersen Antisense Phosphorothioate Oligonucleotide Preparations and Related Pharmacological Activity Impairment. Nucleic Acid Ther, 32, 312–320.

30. Monteleone, G., Stolfi, C., Marafini, I., Atreya, R. and Neurath, M.F. (2022) Smad7 Antisense Oligonucleotide-Based Therapy in Crohn’s Disease: Is it Time to Re-Evaluate? Mol. Diagn. Ther., 26, 477–481.

31. Nelson, P.T., Alafuzoff, I., Bigio, E.H., Bouras, C., Braak, H., Cairns, N.J., Castellani, R.J., Crain, B.J., Davies, P., Tredici, K.D., et al. (2012) Correlation of Alzheimer Disease Neuropathologic Changes With Cognitive Status: A Review of the Literature. J. Neuropathol. Exp. Neurol., 71, 362– 381.

32. Hutton, M., Lendon, C.L., Rizzu, P., Baker, M., Froelich, S., Houlden, H., Pickering-Brown, S., Chakraverty, S., Isaacs, A., Grover, A., et al. (1998) Association of missense and 5′-splice-site mutations in tau with the inherited dementia FTDP-17. Nature, 393, 702–705.

33. Poorkaj, P., Bird, T.D., Wijsman, E., Nemens, E., Garruto, R.M., Anderson, L., Andreadis, A., Wiederholt, W.C., Raskind, M. and Schellenberg, G.D. (1998) Tau is a candidate gene for chromosome 17 frontotemporal dementia. Ann. Neurol., 43, 815–825.

34. DeVos, S.L., Miller, R.L., Schoch, K.M., Holmes, B.B., Kebodeaux, C.S., Wegener, A.J., Chen, G., Shen, T., Tran, H., Nichols, B., et al. (2017) Tau reduction prevents neuronal loss and reverses pathological tau deposition and seeding in mice with tauopathy. Sci Transl Med, 9, eaag0481.

35. Mummery, C.J., Börjesson-Hanson, A., Blackburn, D.J., Vijverberg, E.G.B., Deyn, P.P.D., Ducharme, S., Jonsson, M., Schneider, A., Rinne, J.O., Ludolph, A.C., et al. (2023) Tau-targeting antisense oligonucleotide MAPTRx in mild Alzheimer’s disease: a phase 1b, randomized, placebo-controlled trial. Nat Med, 29, 1437–1447.

36. Saito, T., Mihira, N., Matsuba, Y., Sasaguri, H., Hashimoto, S., Narasimhan, S., Zhang, B., Murayama, S., Higuchi, M., Lee, V.M.Y., et al. (2019) Humanization of the entire murine Mapt gene provides a murine model of pathological human tau propagation. J. Biol. Chem., 294, 12754–12765.

37. Endo, A., Yu, R.T., Fang, F., Choi, H.W. and Shan, M. (2017) Patent WO2017/024264.

38. Eckstein, F. (1983) Phosphorothioate Analogues of Nucleotides—Tools for the Investigation of Biochemical Processes. Angew. Chem. Int. Ed. Engl., 22, 423–439.

39. Sobkowski, M., Jankowska, J., Stawinski, J. and Kraszewski, A. (2005) A CAUTIONARY NOTE ON THE USE OF THE 31P NMR SPECTROSCOPY IN STEREOCHEMICAL CORRELATION ANALYSIS. Nucleosides, Nucleotides Nucleic Acids, 24, 1033–1036.

40. Iwamoto, N., Butler, D.C.D., Svrzikapa, N., Mohapatra, S., Zlatev, I., Sah, D.W.Y., Meena, Standley, S.M., Lu, G., Apponi, L.H., et al. (2017) Control of phosphorothioate stereochemistry substantially increases the efficacy of antisense oligonucleotides. Nature Biotechnology, 35, 845–851.

41. Zhou, X., Kiesman, W.F., Yan, W., Jiang, H., Antia, F.D., Yang, J., Fillon, Y.A., Xiao, L. and Shi, X. (2022) Development of Kilogram-Scale Convergent Liquid-Phase Synthesis of Oligonucleotides. J Org Chem, 87, 2087-2110.

42. Kiełpiński, Ł.J., Funder, E.D., Schmidt, S. and Hagedorn, P.H. (2021) Characterization of Escherichia coli RNase H Discrimination of DNA Phosphorothioate Stereoisomers. Nucleic Acid Ther, 31, 383-391.

43. Duschmalé, J., Schäublin, A., Funder, E., Schmidt, S., Kiełpiński, Ł.J., Nymark, H., Jensen, K., Koch, T., Duschmalé, M., Koller, E., et al. (2022) Investigating discovery strategies and pharmacological properties of stereodefined phosphorodithioate LNA gapmers. Mol Ther - Nucleic Acids, 29, 176–188.

44. Malyala, P. and Singh, M. (2008) Endotoxin limits in formulations for preclinical research. J Pharm Sci, 97, 2041–2044.

45. Lim, M. and Dibble, A. (2016) Osmolality of antisense oligonucleotide parenteral formulations: Implications on counterion dissociation and recommended osmometry techniques. Int J Pharmaceut, 515, 788–799.

