## Supplementary figures for "Discovery and characterization of stereodefined PMO-gapmers targeting tau"

A

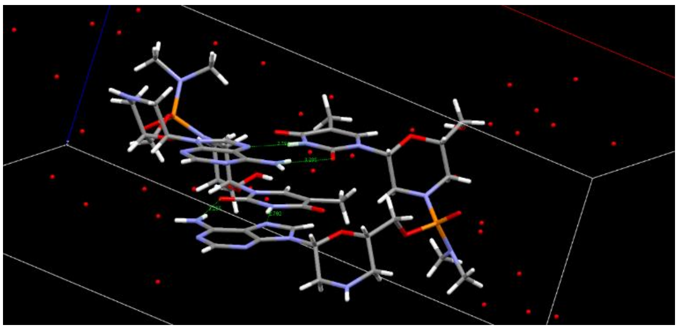

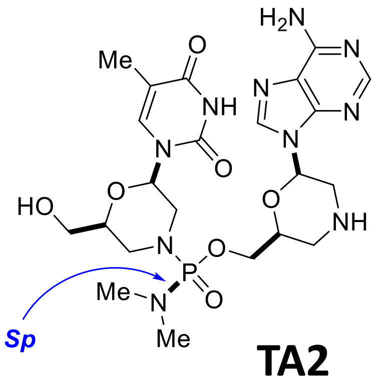


TA2

B

| Dimer | P NMR (ppm) | Assigned stereochmistry |
|---|---|---|
| CT1 | 15.41 | S |
| CT2 | 15.25 | R |
| AT1 | 15.76 | S |
| AT2 | 15.70 | R |
| UT1 | 15.73 | S |
| UT2 | 15.32 | R |
| GT1 | 15.89 | S |
| GT2 | 15.84 | R |
| AC1 | 15.73 | S |
| AC2 | 15.62 | R |
| CA1 | 15.24 | R |
| CA2 | 15.78 | S |
| AG1 | 15.34 | R |
| AG2 | 15.41 | S |

C

| Activated Monomer | P NMR (ppm) | Assigned stereochemistry* |
|---|---|---|
| A1 | 18.416 | S |
| A2 | 18.069 | R |
| T1 | 18.016 | R |
| T2 | 18.355 | S |
| G1 | 18.466 | S |
| G2 | 18.119 | R |
| C1 | 18.027 | R |
| C2 | 18.435 | S |

- A1 and A2 mean the early eluting isomer (A1) and late eluting isomer (A2) of the activated A monomer by a chiral HPLC (37).

**Table S1**. (A) Crystal structure of TA2 dimer unambiguously determined the phosphorus stereochemistry resulted from A2 monomer as Sp (37). (B) The absolute stereochemistry of phosphorus atoms in all the dimers was assigned by relative $^{31}$P NMR chemical shift in pyridine-*d*5 based on the stereochemistry of TA2 dimer. Relative $^{31}$P NMR chemical shifts correlate with phosphorus stereochemistry, which has been widely used for the determination of phosphorus stereochemistry for phosphorothioates (38, 39). CA2 dinucleotide having Sp linkage showed downfield shift (15.78 ppm) than the corresponding Rp isomer CA1 (15.24 ppm). Thus, all the dinucleotides showing downfield shift (CT1, AT1, UT1, GT1, AC1 and AG2) were assigned as *S*p and the others as *R*p (CT2, AT2, UT2, GT2, AC2 and AG1). The relative $^{31}$P chemical shifts were not affected by the nucleotide at the 5' end and CT, AT, UT and GT dinucleotides showed the same trend. (C) Absolute phosphorus stereochemistry of all the monomers were assigned by using relative $^{31}$P NMR chemical shift in $CDCl_3$ based on the stereochemistry of A monomer. The phosphorus stereochemistry of A2 monomer was assigned as *R*p based on inversion during dinucleotide formation.

| Sequence | Compound No. (SEQ ID NO) | Relative expression (*MAPT*/*GAPDH*) | | |
|---|---|---|---|---|
| | | 10 nM | 30 nM | 100 nM |
| **GGGGACTCGCTGACATGG** | **ASO-369** (SEQ ID NO: 1) | 0.771 | 0.675 | 0.581 |
| **TGGGTGTAGCGAGAATCC** | **ASO-373** (SEQ ID NO: 2) | 0.863 | 0.542 | 0.401 |
| **GGGTGCACTAGTTTATAG** | **ASO-380** (SEQ ID NO: 3) | 0.811 | 0.587 | 0.377 |
| **GGGGTCTTCTAATATCCT** | **ASO-388** (SEQ ID NO: 4) | 0.619 | 0.410 | 0.291 |
| **AGGTTCTCGCTATATCGC** | **ASO-389** (SEQ ID NO: 5) | 0.850 | 0.628 | 0.357 |
| **GAGTTAGAAGCTTTGACT** | **ASO-401** (SEQ ID NO: 6) | 0.801 | 0.480 | 0.378 |
| **GCAGATGACCCTTAGACA** | **ASO-409** (SEQ ID NO: 7) | 0.866 | 0.587 | 0.373 |
| **CAAACCTGTCACACCCGA** | **ASO-413** (SEQ ID NO: 8) | 0.898 | 0.785 | 0.547 |
| **TTAAACCCCATAGACATA** | **ASO-417** (SEQ ID NO: 9) | 0.959 | 0.865 | 1.070 |
| **GAGGCCCAAATGATCACA** | **ASO-418** (SEQ ID NO: 10) | 0.972 | 0.853 | 0.822 |
| **TGGATTTAGCAGTAGGGT** | **ASO-463** (SEQ ID NO: 11) | 0.896 | 0.710 | 0.441 |
| **AGCAGATGACCCTTAGAC** | **ASO-467** (SEQ ID NO: 12) | 0.806 | 0.618 | 0.496 |
| **AGCCGGCATACAGTATAT** | **ASO-468** (SEQ ID NO: 13) | 0.955 | 0.698 | 0.578 |
| **TGTGCTCTTTATGGATGG** | **ASO-469** (SEQ ID NO: 14) | 0.764 | 0.632 | 0.435 |
| **GGATTTAGCAGTAGGGTG** | **ASO-470** (SEQ ID NO: 15) | 1.245 | 0.844 | 0.480 |
| **CCCCATGACTACAGTGTG** | **ASO-473** (SEQ ID NO: 16) | 0.893 | 0.747 | 0.442 |
| **GCTTTTGTGACCAGGGAC** | **ASO-474** (SEQ ID NO: 17) | 0.793 | 0.381 | 0.173 |

**Table S2**. In vitro activity of 17 selected stereorandom 5-8-5 PMO-gapmers. SH-SY5Y cells were transfected with 10, 30 or 100 nM ASOs. After 2 days incubation, *MAPT* mRNA was measured.

| Sequence | Compound No. (SEQ ID NO) | Relative expression (*MAPT*/*GAPDH*) | | |
|---|---|---|---|---|
| | | 30 nM | 100 nM | 300 nM |
| **TGGATTTAGCAGTAGGGT** | **ASO-483** (SEQ ID NO: 11) | 0.613 | 0.492 | [not tested] |
| **GCTTTTGTGACCAGGGAC** | **ASO-484** (SEQ ID NO: 17) | 0.628 | 0.484 | 0.320 |
| **AGGTTCTCGCTATATCGC** | **ASO-485** (SEQ ID NO: 5) | 0.646 | 0.520 | 0.429 |
| **AGCAGATGACCCTTAGAC** | **ASO-486** (SEQ ID NO: 12) | 0.735 | 0.576 | 0.415 |
| **TGTGCTCTTTATGGATGG** | **ASO-487** (SEQ ID NO: 14) | 0.812 | 0.663 | 0.541 |
| **CCCCATGACTACAGTGTG** | **ASO-488** (SEQ ID NO: 16) | 0.705 | 0.565 | 0.439 |
| **TGGGTGTAGCGAGAATCC** | **ASO-489** (SEQ ID NO: 2) | 0.600 | 0.450 | 0.324 |
| **GGGTGCACTAGTTTATAG** | **ASO-490** (SEQ ID NO: 3) | 0.638 | 0.520 | 0.442 |
| **GGGGTCTTCTAATATCCT** | **ASO-491** (SEQ ID NO: 4) | 0.556 | 0.423 | 0.311 |
| **GCAGATGACCCTTAGACA** | **ASO-492** (SEQ ID NO: 7) | 0.709 | 0.545 | 0.436 |
| **GAGGCCCAAATGATCACA** | **ASO-493** (SEQ ID NO: 10) | 0.747 | 0.528 | 0.486 |
| **TTAAACCCCATAGACATA** | **ASO-494** (SEQ ID NO: 9) | 0.989 | 0.914 | 0.801 |

**Table S3**. In vitro activity of 12 selected stereorandom 4-10-4 PMO-gapmers. SH-SY5Y cells were transfected with 30, 100 or 300 nM ASOs. After 2 days incubation, *MAPT* mRNA was measured.

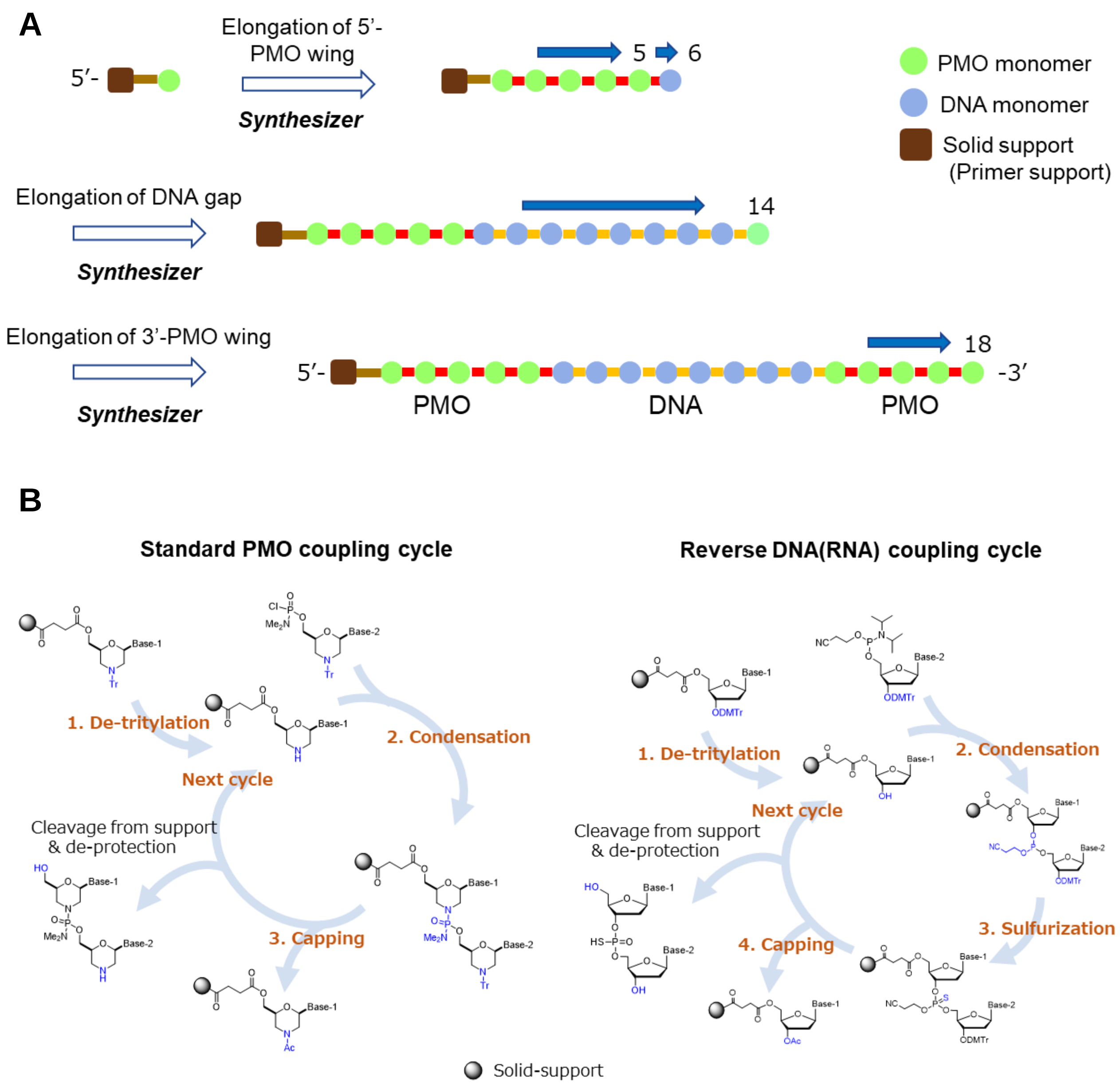


**Figure S1**. (A) Schematic drawing and (B) Synthetic cycle of solid phase synthesis of stereorandom PMO-gapmers.

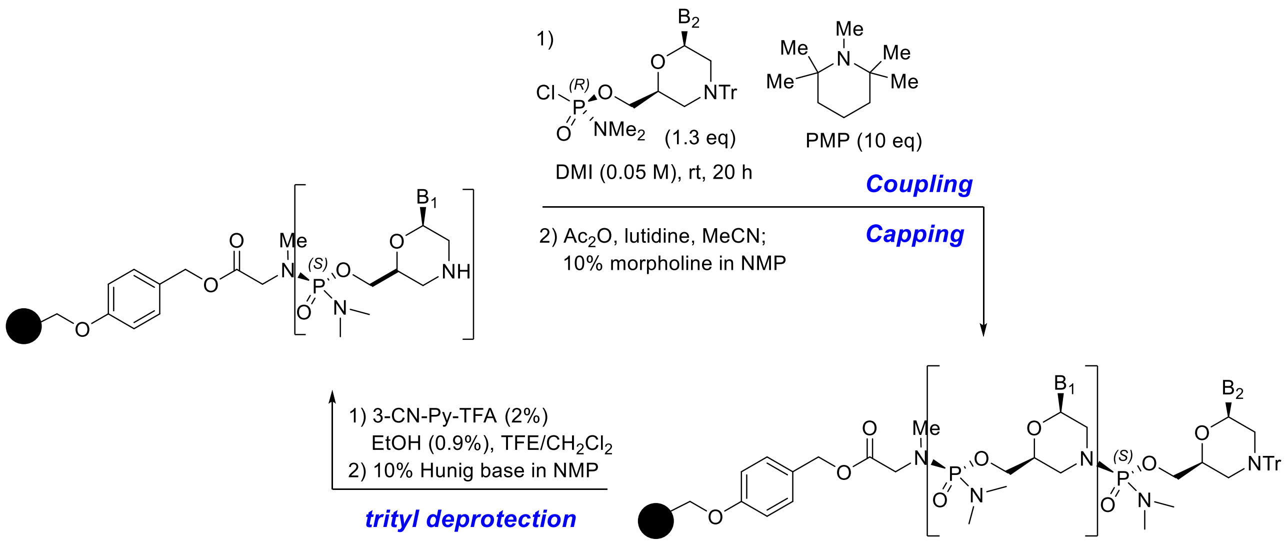


**Figure S2**. General scheme for the solid phase synthesis of stereopure PMOs.

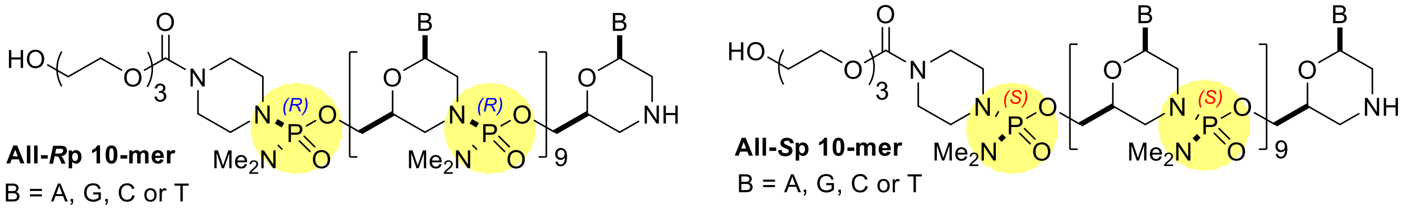


**Tm (°C) of PMO/RNA Duplex**

| P-chirality | A | G | T | C |
|---|---|---|---|---|
| *S*p | 68.5 | 88.8 | 31.4 | 77.3 |
| *R*p | 37.0 | 73.1 | 19.3 | 63.5 |
| **Δ Tm (*Sp*-*Rp*)** | **+31.5** | **+15.7** | **+12.1** | **+13.8** |

**Figure S3.** Tm analysis of homogeneous PMO 10-mers of A, C, T and G. The PMO 10-mers assigned as *S*p based on $^{31}$P NMR chemical shifts consistently showed higher Tm than the corresponding *R*p 10-mers.

NM_001123066.3

| ASO No<br>5-8-5 / 4-10-4 | Start | End | Sequence | GC content | PredictedTm |
|---|---|---|---|---|---|
| ASO-373 / 489 | 93904 | 93921 | TGGGTGTAGCGAGAATCC | 56 | 62 |
| ASO-409 / 492 | 102324 | 102341 | GCAGATGACCCTTAGACA | 50 | 59 |
| ASO-467 /486 | 102325 | 102342 | AGCAGATGACCCTTAGAC | 50 | 62 |
| ASO-469 / 487 | 124521 | 124538 | TGTGCTCTTTATGGATGG | 44 | 50 |

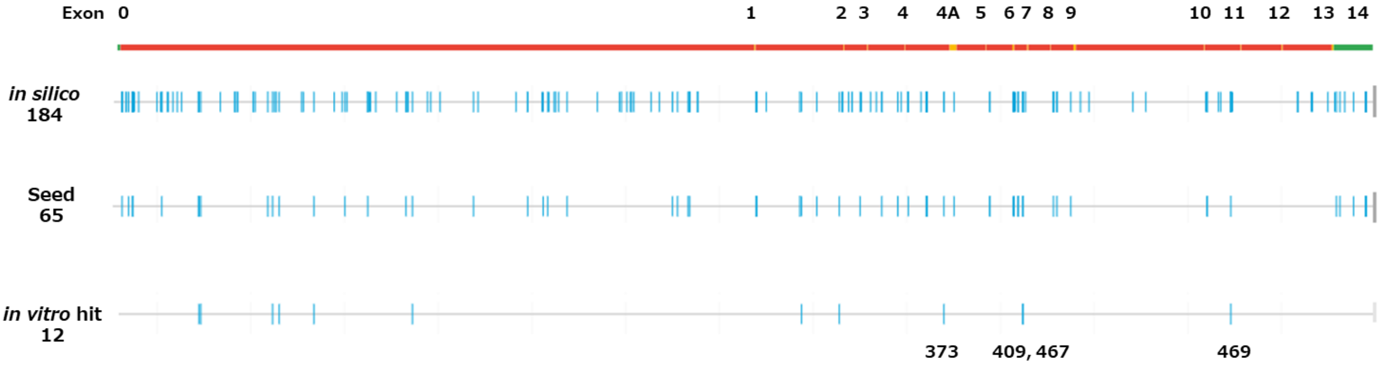


red: intron, yellow: exon

**Figure S4.** Best 4 ASO sequences and their location in *MAPT* gene.

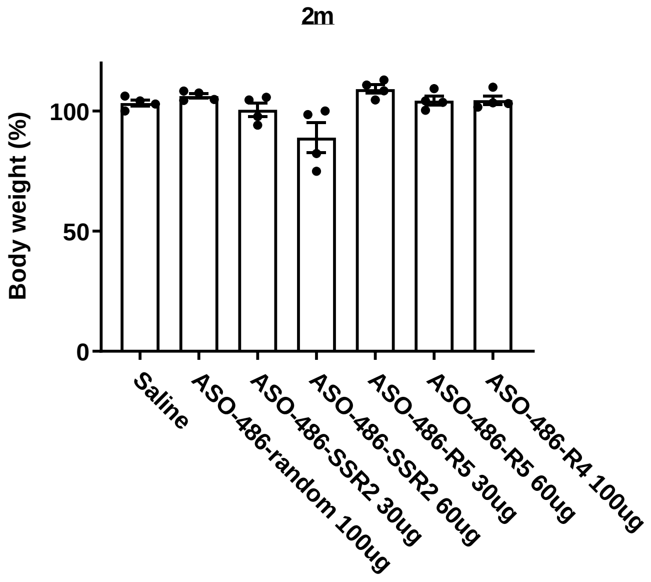


**Figure S5**. Body weight changes 2 months after treatment with stereodefined ASO-486. hTau KI mice (4 mice per group) were treated with 30-100 μg of ASO by ICV injection.

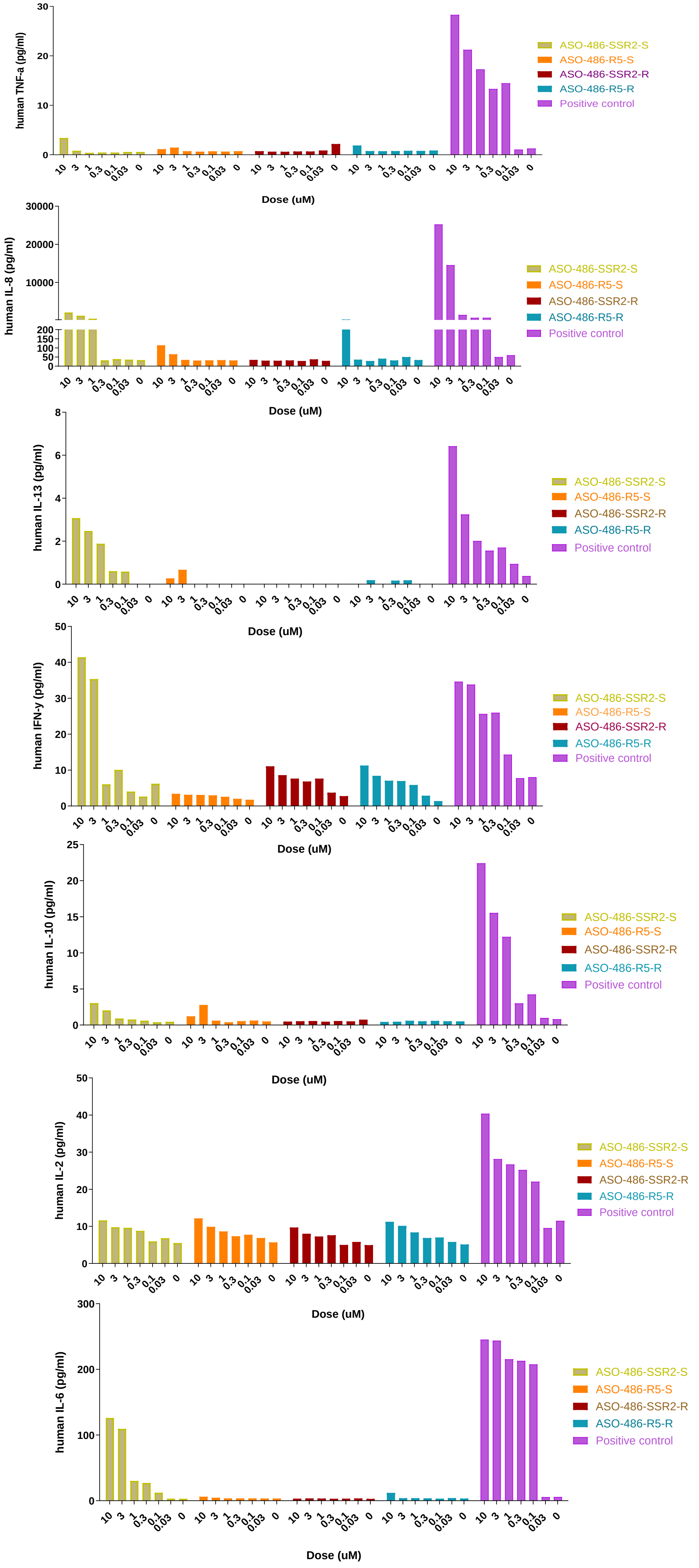


**Figure S6**. Human PBMC cytokine release assay (ODN2006 was used as the positive control).

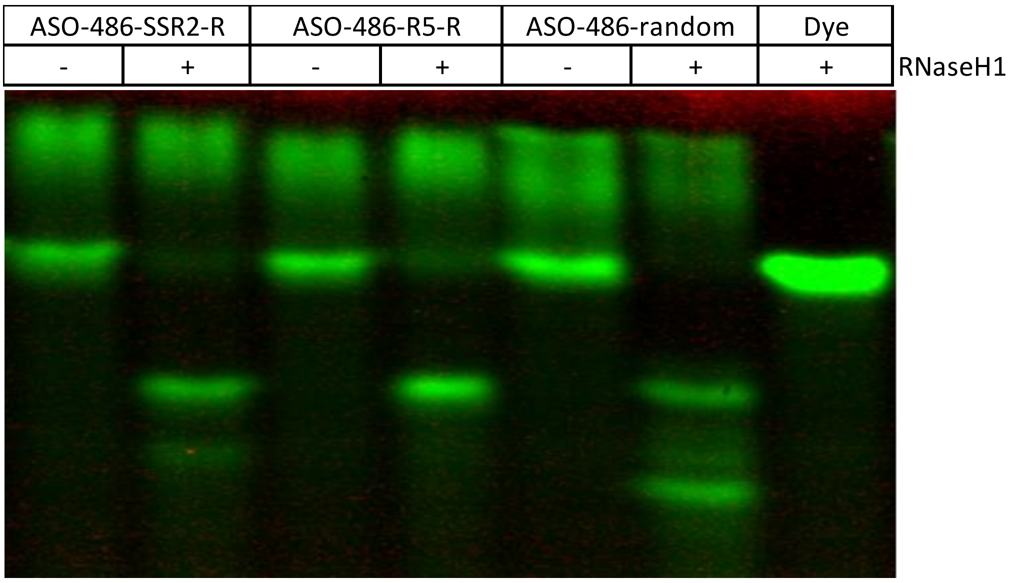


**Figure S7**. Human RNase H1 cleavage of ASO-486-SSR2-R, ASO-486-R5-R and ASO-486-random with 5'-FAM RNA probe.

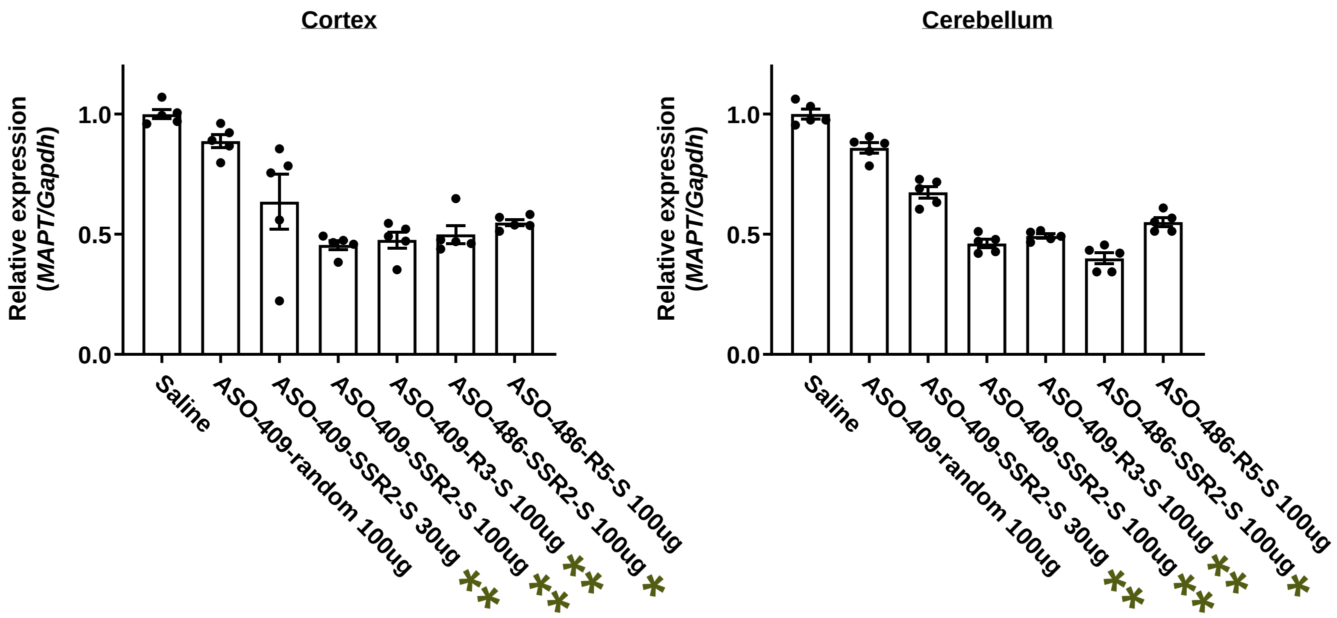


***** : Abnormal gait and motor deficit were observed up to 2m
******: Mortality was observed up to 2m

**Figure S8**. In vivo KD activity of completely stereodefined SCOT-409-SSR2, ASO-409-R3, ASO-486-SSR2, and ASO-486-R5 in cortex and cerebellum. hTau KI mice (5 mice per group) were treated with 30 - 100 μg of ASO by ICV injection. *MAPT* mRNA was measured after 3 days.
