## Supplementary information for "Discovery and characterization of stereodefined PMO-gapmers targeting tau"

The chemical names for the compounds in the following examples were created based on the chemical structures using “E-Notebook 2014” version 13 or E-Notebook version 18.1.1.0073 (PerkinElmer Co., Ltd.).

In Examples, flash chromatography separations were performed using SNAP cartridges (Biotage®) or Hi-FlashTM Column Silicagel or Amino (YAMAZENE CORPORATION).

Proton nuclear magnetic resonance (NMR) spectra were recorded on a JEOL JNM-ECZ 400S/L1 or JEOL JNM-ECZ 500R/S1 or Varian Inova 500 MHz or Varian Inova 400 MHz, or Bruker 400 MHz spectrometer. Chemical shifts are reported in the unit of a (ppm) and coupling constants are reported in the unit of Hertz (Hz). Abbreviations for splitting patterns are as follows: s: singlet; d: doublet; t: triplet; m: multiplet; and brs: broad singlet. $^{31}$P nuclear magnetic resonance (NMR) spectra were recorded on Varian Inova 400 MHz or Bruker 400 MHz spectrometer. Chemical shifts are reported in the unit of a (ppm). Abbreviation for splitting patterns is as follows: s: singlet.

Mass spectrometry was carried out using an Acquity UPLC and SQD2 (Waters), or a Acquity UPLC and Synapt G2 (Waters), or a Nexera X3 UHPLC (Shimadzu) and a Q Exactive Plus (ThermoFisherScientific).

#### 1. Synthesis of monomers and loading of morpholino monomers on solid support

Synthesis of ((2R,3S,5R)-3-(bis(4-methoxyphenyl)(phenyl)methoxy)-5-(5-methyl-2,4-dioxo-3,4-dihydropyrimidin-1(2H)-yl)tetrahydrofuran-2-yl)methyl dimethylphosphoramidochloridate (**S2**)

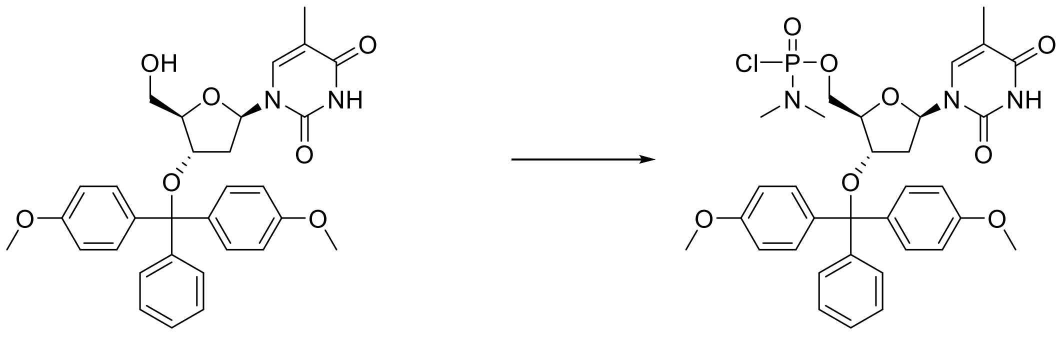

Method-1

To a solution of 3'-O-[Bis(4-methoxyphenyl)(phenyl)methyl]thymidine (**S1**, CAS 76054-81-4) (3.00 g, 5.51 mmol) in $CH_2Cl_2$ (20 mL) was added 1-methylimidazole (0.524 mL, 6.61 mmol), 2,6-lutidine (1.60 mL, 13.8 mmol), followed by (dimethylamino)phosphonoyl dichloride (1.63 mL, 13.8 mmol) in one portion with ice-cooling. The resulting solution was stirred for 6 h at room temperature. To 5 % citric acid aqueous solution (60 mL) was added the reaction mixture with ice-cooling. The mixture was separated and the aqueous layer was extracted with $CH_2Cl_2$. The organic layer was washed with brine, dried over $Na_2SO_4$, filtered and concentrated in vacuo to give the crude. Silica gel column chromatography of the residue using 50 % to 80 % EtOAc/Heptane afforded**S2** (2.71 g).

Method-2

To a solution of 3'-O-[Bis(4-methoxyphenyl)(phenyl)methyl]thymidine (**S1**, 3.00 g, 5.51 mmol) in $CH_3CN$ (55 mL) and $CH_2Cl_2$ (55 mL) was added lithium bromide (1.58 g, 18.2 mmol) and DBU (2.74 mL, 18.2 mmol), followed by (dimethylamino)phosphonoyl dichloride (0.853 mL, 7.16 mmol) in one portion at 0 °C and stirred at the same temperature for 15 min. The resulting solution was stirred at room temperature for 1 h. To the reaction mixture was added a solution of citric acid monohydrate (5.0 g, 23.8 mmol) in water (95 mL) at 0 °C. To the mixture was added $CH_2Cl_2$ (50 mL) and the mixture was separated by ISOLUTE™ phase separater (Biotage) and the organic layer was concentrated in vacuo to give the crude. Silica gel column chromatography of the residue using 50% to 100% EtOAc/Heptane afforded **S2** (1.18 g). $^1$H NMR (396 MHz, CHLOROFORM-d) δ 7.28-7.36 (m, 7 H), 7.94 (br s, 1 H), 7.42-7.46 (m, 2 H), 6.80-6.88 (m, 4 H), 6.34-6.45 (m, 1 H), 4.26-4.35 (m, 1 H), 3.86-4.03 (m, 2 H), 3.79 (s, 6 H), 3.45-3.57 (m, 1 H), 2.59-2.67 (m, 7 H), 2.04-2.20 (m, 1 H), 1.84-1.91 (m, 3 H), 1.61-1.73 (m, 1 H).

Synthesis of ((2R,3S,5R)-5-(4-benzamido-2-oxopyrimidin-1(2H)-yl)-3-(bis(4-methoxyphenyl)(phenyl)methoxy)tetrahydrofuran-2-yl)methyl dimethylphosphoramidochloridate (**S4**)

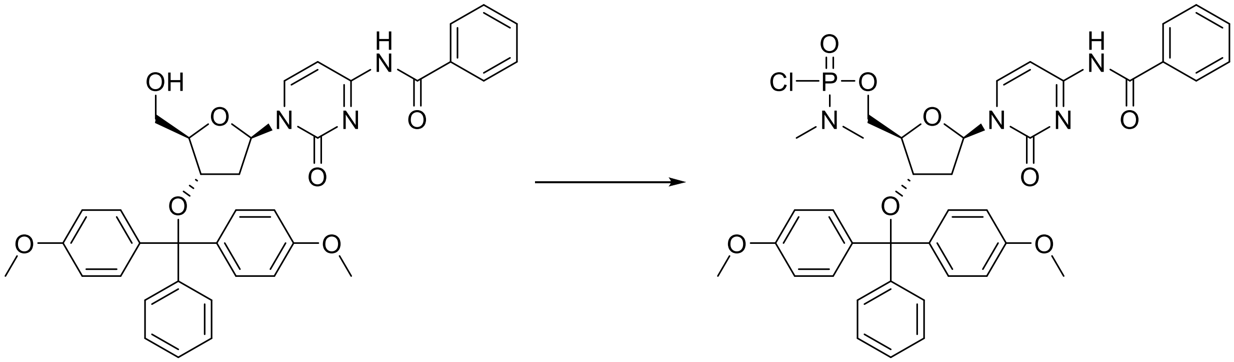

S3 S4

To a solution of N-Benzoyl-3'-O-[bis(4-methoxyphenyl)(phenyl)methyl]-2'-deoxycytidine (**S3**, CAS 140712-80-7) (2.00 g, 3.16 mmol) in $CH_3CN$ (20 mL) and $CH_2Cl_2$ (28 mL) was added lithium bromide (0.850 g, 9.78 mmol) and DBU (1.46 mL, 9.78 mmol), followed by (dimethylamino)phosphonoyl dichloride (0.560 mL, 4.73 mmol) in one portion at -10 °C. The resulting solution was stirred for 4 h at -10 °C. To the reaction mixture was added 5 % citric acid aqueous solution (220 mL). The mixture was stirred at -10 °C for 5 min. To the mixture was added $CH_2Cl_2$ and then the phases were separated. The aqueous layer was extracted with $CH_2Cl_2$, and the combined organic layer was washed with water, then washed with brine, dried over $Na_2SO_4$, filtered and concentrated in vacuo to give the crude. Silica gel column chromatography of the residue using 60 % to 80 % EtOAc/Heptane afforded **S4** (1.49 g).

$^1$H NMR (CHLOROFORM-d, 396 MHz) δ 8.02-8.05 (m, 1H), 7.87 (br d, 2H, *J*=7.7 Hz), 7.60 (t, 1H, *J*=7.7 Hz), 7.44-7.52 (m, 5H), 7.28-7.36 (m, 6H), 7.21-7.26 (m, 1H), 6.83-6.85 (m, 4H), 6.38-6.42 (m, 1H), 4.29-4.32 (m, 1H), 3.99-4.04 (m, 0.5H), 3.92-3.93 (m, 0.5H), 3.83-3.87 (m, 1H), 3.79 (s, 6H), 3.44-3.52 (m, 1H), 2.63 (s, 1.5H), 2.63 (s, 1.5H), 2.60 (s, 1.5H), 2.59 (s, 1.5H), 1.63-1.73 (m, 2H).

MS (ESI) m/z: $[M+H]^+$ calcd for $C_{39}H_{41}ClN_4O_8P$: 759.235; Found:759.372.

<u>Synthesis of ((2R,3S,5R)-5-(6-benzamido-9H-purin-9-yl)-3-(bis(4-methoxyphenyl)(phenyl)methoxy)tetrahydrofuran-2-yl)methyl dimethylphosphoramidochloridate (**S6**)</u>

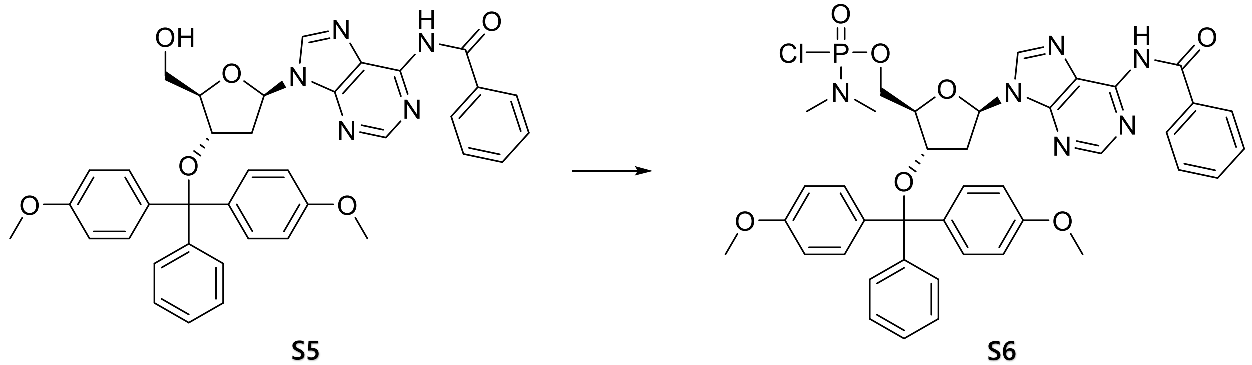

S5 S6

To a solution of N-Benzoyl-3'-O-[bis(4-methoxyphenyl)(phenyl)methyl]-2'-deoxyadenosine (**S5**, CAS 140712-79-4) (3.00 g, 4.56 mmol), 1-methylimidazole (0.434 mL, 5.47 mmol), and 2,6-lutidine (1.32 mL, 11.4 mmol) in $CH_2Cl_2$ (22.6 mL, 351.2 mmol) at 0 °C was added (dimethylamino)phosphonoyl dichloride (1.35 mL, 11.4 mmol). The mixture was gradually warmed to room temperature and stirred at room temperature for 5 h. The reaction mixture was poured into the ice-cold 5% citric acid aqueous solution, then extracted with EtOAc (2 times). The combined organic layers were washed with brine, dried over $Na_2SO_4$, filtered, and concentrated in vacuo. Silica gel column chromatography of the residue using 20 % to 80% EtOAc/Heptane afforded **S6** (2.10 g).

$^1$H NMR (396 MHz, CHLOROFORM-d) δ ppm 8.84-8.95 (m, 1H), 8.78 (s, 1H), 8.13 (m, 1H), 8.00 (m, 2H), 7.58-7.64 (m, 1H), 7.47-7.53 (m, 4H), 7.28-7.42 (m, 6H), 6.79-6.92 (m, 4H), 6.54 (m, 1H), 4.48-4.57 (m, 1H), 4.06-4.17 (m, 2H), 3.94-4.05 (m, 1H), 3.80 (m, 1H), 3.79 (s, 6H), 2.59-2.60 (m, 3H), 2.55-2.56 (m, 3H), 2.33-2.46 (m, 1H), 2.11-2.30 (m, 1H).

MS (ESI) m/z: [M+H]+ Calcd for $C_{40}H_{41}ClN_6O_7P$: 783.246; Found:783.368.

<u>Synthesis of ((2R,3S,5R)-3-(bis(4-methoxyphenyl)(phenyl)methoxy)-5-(2-isobutyramido-6-oxo-1,6-dihydro-9H-purin-9-yl)tetrahydrofuran-2-yl)methyl dimethylphosphoramidochloridate</u>

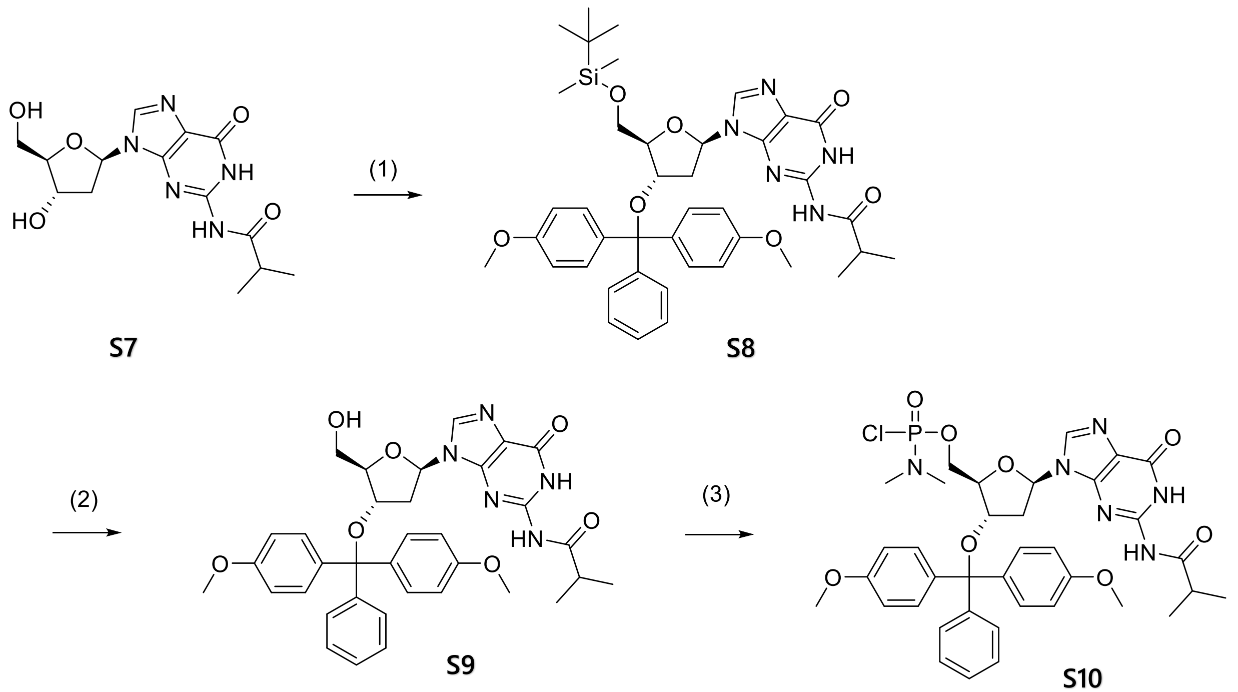

(1) N-(9-((2R,4S,5R)-4-(bis(4-methoxyphenyl)(phenyl)methoxy)-5-(((tert-butyldimethylsilyl)oxy)methyl)tetrahydrofuran-2-yl)-6-oxo-6,9-dihydro-1H-purin-2-yl)isobutyramide

To a solution of N-(9-((2R,4S,5R)-4-hydroxy-5-(hydroxymethyl)tetrahydrofuran-2-yl)-6-oxo-6,9-dihydro-1H-purin-2-yl)isobutyramide (**S7**, CAS 68892-42-2) (5.00 g, 14.8 mmol) in pyridine (33.5 mL, 0.414 mol) was added tert-butylchlorodimethylsilane (3.35 g, 22.2 mmol) with ice-cooling. The resulting solution was stirred for 190 min at room temperature. To the solution was added 4,4'-(chloro(phenyl)methylene)bis(methoxybenzene) (8.54 g, 25.2 mmol). The resulting solution was stirred for 2 h at 50 °C. To the reaction mixture was added sat. $NaHCO_3$ aqueous solution (150 mL) and then the phases were separated. The aqueous layer was extracted with $CH_2Cl_2$ twice. The combined organic layer was washed with water and brine, then dried over $Na_2SO_4$, filtered and concentrated in vacuo to give the crude. Silica gel column chromatography of the residue using 33 % to 66 % EtOAc/Heptane afforded **S8** (8.78 g).

$^1$H NMR (CHLOROFORM-d, 396 MHz) δ 11.87 (s, 1H), 7.98 (s, 1H), 7.80 (s, 1H), 7.45-7.47 (m, 2H), 7.28-7.36 (m, 6H), 7.21-7.24 (m, 1H), 6.82-6.84 (m, 4H), 6.20-6.24 (m, 1H), 4.36-4.38 (m, 1H), 4.05-4.07 (m, 1H), 3.78 (s, 6H), 3.58-3.62 (m, 1H), 3.31-3.35 (m, 1H), 2.54-2.61 (m, 1H), 1.94-2.01 (m, 1H), 1.83-1.88 (m, 1H), 1.27-1.29 (m, 6H), 0.77 (s, 9H), -0.07 (s, 3H), -0.09 (s, 3H).

MS (ESI) m/z: $[M+H]^+$ Calcd for $C_{41}H_{52}N_5O_7Si$: 754.363; Found: 754.387.

(2) <u>N-(9-((2R,4S,5R)-4-(bis(4-methoxyphenyl)(phenyl)methoxy)-5-(hydroxymethyl)tetrahydrofuran-2-yl)-6-oxo-6,9-dihydro-1H-purin-2-yl)isobutyramide (**S8**)</u>

To a solution of N-(9-((2R,4S,5R)-4-(bis(4-methoxyphenyl)(phenyl)methoxy)-5-(((tert-butyldimethylsilyl)oxy)methyl)tetrahydrofuran-2-yl)-6-oxo-6,9-dihydro-1H-purin-2-yl)isobutyramide (**S8**, 4.50 g, 5.97 mmol) in THF (41 mL) was added tetra-n-butylammonium fluoride (1 M THF solution, 6.57 mL, 6.57 mmol). The resulting solution was stirred for 18 h at room temperature. The reaction mixture was diluted with EtOAc (400 mL) and washed with sat. $NH_4Cl$ aqueous solution (200 mL), sat. $NaHCO_3$ aqueous solution (200 mL) and brine (200 mL). The organic layer was dried over $Na_2SO_4$, filtered and concentrated in vacuo to give the crude. Silica gel column chromatography of the residue using 0 % to 20 % MeOH/$CH_2Cl_2$ afforded the mixture containing target material. Further silica gel column chromatography of the mixture using 1 % to 5 % MeOH/$CH_2Cl_2$ afforded **S9** (3.03 g).

$^1$H NMR (CHLOROFORM-d, 396 MHz) δ 12.00 (br s, 1H), 8.24 (br s, 1H), 7.65 (s, 1H), 7.43-7.46 (m, 2H), 7.28-7.35 (m, 6H), 7.21-7.23 (m, 1H), 6.81-6.85 (m, 4H), 6.15 (dd, 1H, *J*=5.3, 9.7 Hz), 5.14 (br d, 1H, *J*=11.0 Hz), 4.50 (d, 1H, *J*=5.7 Hz), 4.05 (s, 1H), 3.78 (s, 3H), 3.78 (s, 3H), 3.68-3.71 (m, 1H), 3.27 (t, 1H, *J*=11.0 Hz), 2.55-2.62 (m, 1H), 2.41 (ddd, 1H, *J*=5.7, 9.7, 13.6 Hz), 1.70 (dd, 1H, *J*=5.3, 13.6 Hz), 1.22-1.23 (m, 6H).

MS (ESI) m/z: $[M+H]^+$ Calcd for $C_{35}H_{38}N_5O_7$: 640.277; Found: 640.615.

(3) <u>((2R,3S,5R)-3-(bis(4-methoxyphenyl)(phenyl)methoxy)-5-(2-isobutyramido-6-oxo-1,6-dihydro-9H-purin-9-yl)tetrahydrofuran-2-yl)methyl dimethylphosphoramidochloridate (**S10**)</u>

To a solution of N-(9-((2R,4S,5R)-4-(bis(4-methoxyphenyl)(phenyl)methoxy)-5-(hydroxymethyl)tetrahydrofuran-2-yl)-6-oxo-6,9-dihydro-1H-purin-2-yl)isobutyramide (**S9**, 2.38 g, 3.73 mmol) in $CH_3CN$ (32 mL) and $CH_2Cl_2$ (32 mL) were added lithium bromide (1.29 g, 14.9 mmol) and DBU (2.25 mL, 14.9 mmol), followed by (dimethylamino)phosphonoyl dichloride (0.887 mL, 7.45 mmol) in one portion with ice-cooling. The resulting solution was stirred for 45 min with ice-cooling. To the reaction mixture was added 5 % citric acid aqueous solution (300 mL). The mixture was stirred with ice-cooling for 5 min. To the mixture was added $CH_2Cl_2$ (270 mL) and then the phases were separated. The aqueous layer was extracted with $CH_2Cl_2$ twice, and the combined organic layer was washed with water. The water layer was extracted with $CH_2Cl_2$ twice.The combined organic layer was dried over $Na_2SO_4$, filtered and concentrated in vacuo to give the crude. Silica gel column chromatography of the residue using 0 % to 16 % THF/ $CH_2Cl_2$ afforded **S10** (2.08 g).

$^1$H NMR (CHLOROFORM-d, 396 MHz) δ 12.15 (s, 0.5H), 12.11 (s, 0.5H), 10.01 (s, 0.5H), 9.93 (s, 0.5H), 7.64 (s, 0.5H), 7.61 (s, 0.5H), 7.44-7.47 (m, 2H), 7.30-7.36 (m, 6H), 7.21-7.24 (m, 1H), 6.83-6.86 (m, 4H), 6.27-6.31 (m, 0.5H), 6.16-6.20 (m, 0.5H), 4.70-4.76 (m, 0.5H), 4.48-4.49 (m, 0.5H), 4.32-4.38 (m, 1H), 4.20-4.25 (m, 1H), 3.99-4.03 (m, 0.5H), 3.85-3.88 (m, 0.5H), 3.78 (s, 3H), 3.78 (s, 3H), 2.65-2.76 (m, 2H), 2.62 (s, 1.5H), 2.61 (s, 1.5H), 2.59 (s, 1.5H), 2.58 (s, 1.5H), 1.94-1.99 (m, 0.5H), 1.67-1.72 (m, 0.5H), 1.16-1.21 (m, 6H).

MS (ESI) m/z: $[M+H]^+$ Calcd for $C_{37}H_{43}ClN_6O_8P$: 765.256; Found:765.383.

Synthesis of ((2S,6R)-6-(2-isobutyramido-6-oxo-1,6-dihydro-9H-purin-9-yl)-4-tritylmorpholin-2-yl)methyl dimethylphosphoramidochloridate (**S12**)

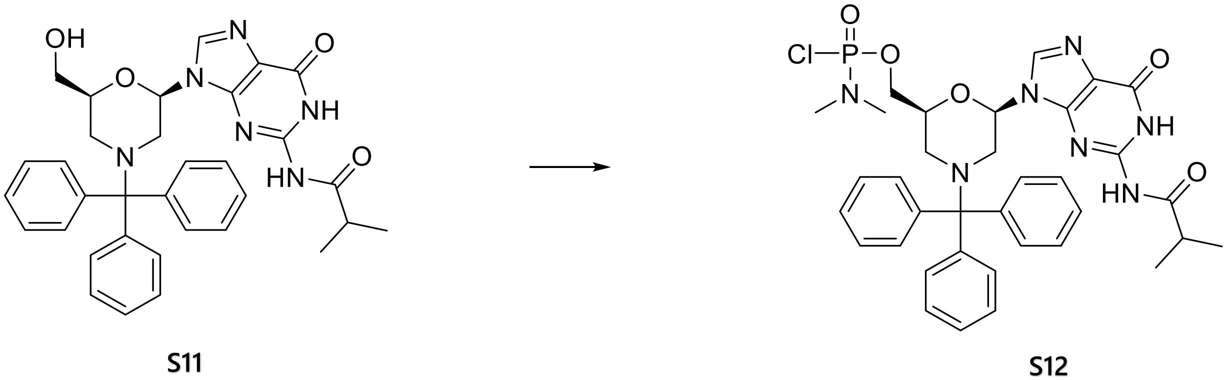

To a solution of N-(9-((2R,6S)-6-(hydroxymethyl)-4-tritylmorpholin-2-yl)-6-oxo-6,9-dihydro-1H-purin-2-yl)isobutyramide (**S11**, 4.00 g, 6.91 mmol) in $CH_3CN$ (59 mL) and $CH_2Cl_2$ (59 mL) were added lithium bromide (2.40 g, 27.6 mmol) and DBU (4.17 mL, 27.6 mmol), followed by (dimethylamino)phosphonoyl dichloride (1.65 mL, 13.8 mmol) in one portion with ice-cooling. The resulting solution was stirred for 35 min with ice bath. To the reaction mixture was added 5 % citric acid aqueous solution (220 mL). The mixture was stirred with ice bath for 5 min. To the mixture was added $CH_2Cl_2$ (180 mL) and then the layers were separated. The aqueous layer was extracted with $CH_2Cl_2$ twice.The combined organic layer was dried over

Na2SO4, filtered and concentrated in vacuo to give the crude. Silica gel column chromatography of the residue using 0 % to 16 % THF/ $CH_2Cl_2$ afforded **S12** (2.70 g).

1H NMR (CHLOROFORM-d, 396 MHz) δ 11.98 (br s, 0.5H), 11.97 (br s, 0.5H), 8.62 (s, 0.5H), 8.46 (s, 0.5H), 7.58 (s, 0.5H), 7.57 (s, 0.5H), 7.44 (br s, 6H), 7.28-7.31 (m, 6H), 7.17-7.21 (m, 3H), 5.96-6.01 (m, 1H), 4.42-4.47 (m, 1H), 4.02-4.18 (m, 2H), 3.41-3.44 (m, 1H), 3.19-3.23 (m, 1H), 2.66-2.71 (m, 1H), 2.64 (s, 1.5H), 2.63 (s, 1.5H), 2.61 (s, 1.5H), 2.59 (s, 1.5H), 1.69-1.75 (m, 1H), 1.50-1.57 (m, 1H), 1.26-1.31 (m, 6H). MS (ESI) m/z: $[M+H]^+$ Calcd for $C_{35}H_{40}ClN_7O_5P$: 704.251; Found: 704.380.

<u>Synthesis of ((2S,6R)-6-(6-(2-cyanoethoxy)-2-isobutyramido-9H-purin-9-yl)-4-tritylmorpholin-2-yl)methyl (2-cyanoethyl) diisopropylphosphoramidite (**S14**)</u>

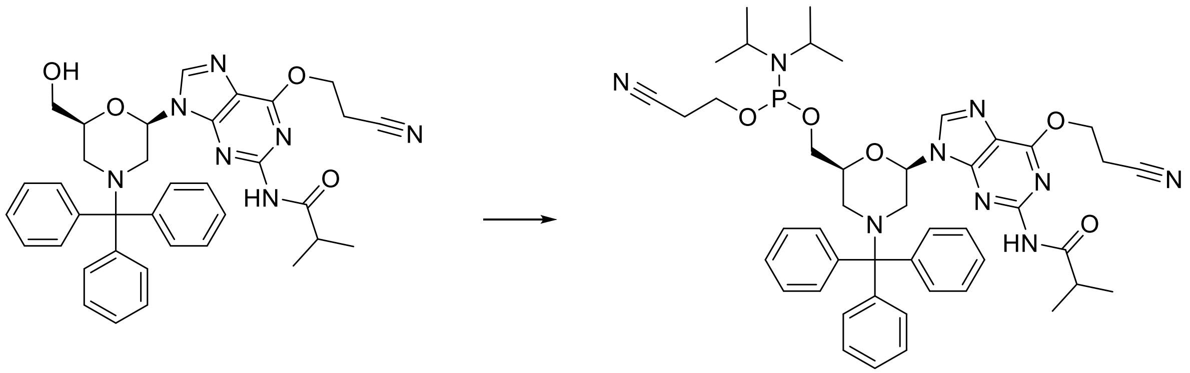

S13 S14

To a solution of N-(6-(2-cyanoethoxy)-9-((2R,6S)-6-(hydroxymethyl)-4-tritylmorpholin-2-yl)-9H-purin-2-yl)isobutyramide (**S13**, 3.00 g, 4.75 mmol) in $CH_2Cl_2$ (30 mL) was added DIPEA (1.82 mL, 10.5 mmol), followed by 2-CYANOETHYL N,N-DIISOPROPYLCHLOROPHOSPHORAMIDITE (1.17 mL, 5.22 mmol) at 0 °C.The reaction mixture was stirred for 1 h at room temperature. To the mixture was added sat. $NaHCO_3$ aqueous solution at 0 °C. The organic layer was separated by ISOLUTE™ phase separater (Biotage) and concentrated in vacuo to give the crude. Silica gel column chromatography of the residue using 50% to 100% EtOAc/Heptane afforded **S14** (1.50 g).

$^1$H NMR (400 MHz, CHLOROFORM-d) δ 7.76-7.82 (m, 2 H), 7.43-7.53 (m, 5 H), 7.26-7.32 (m, 6 H), 7.15-7.22 (m, 3 H), 6.18-6.25 (m, 1 H), 4.69-4.83 (m, 2 H), 4.32-4.41 (m, 1 H), 3.43-3.76

(m, 8 H), 3.21-3.33 (m, 1 H), 2.93-3.09 (m, 3 H), 2.45-2.57 (m, 2 H), 1.68-1.81 (m, 1 H), 1.32-1.36 (m, 6 H), 1.10-1.14 (m, 6 H), 0.99-1.06 (m, 6 H).

Synthesis of 4-(((2S,6R)-6-(5-methyl-2,4-dioxo-3,4-dihydropyrimidin-1(2H)-yl)-4-tritylmorpholin-2-yl)methoxy)-4-oxobutanoic acid loaded onto aminomethylpolystyrene resin (**S16**)

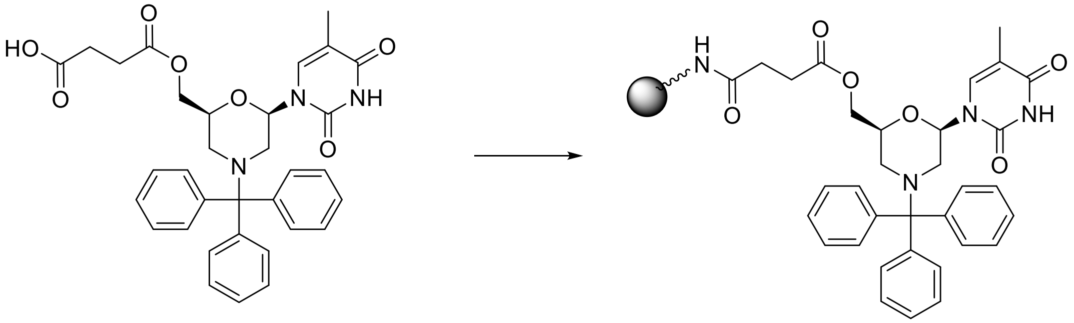

**S15** **S16**

4-(((2S,6R)-6-(5-Methyl-2,4-dioxo-3,4-dihydropyrimidin-1(2H)-yl)-4-tritylmorpholin-2-yl)methoxy)-4-oxobutanoic acid (**S15**, CAS 1362664-41-2) (360 mg, 0.617 mmol) was dissolved in DMF (15.4 mL). HATU (793 mg, 2.09 mmol) and DIPEA (0.539 mL, 3.08 mmol) were added and then Aminomethyl Polystyrene Resin (Primer Support$^{TM}$ 5G Amino, 29-0999-92, manufactured by GE Healthcare) (2.00 g, amine content: 400 μmol/g) was added to the reaction mixture and gently shaken at room temperature on Bio-shaker (110 rpm) for 12 h. The resin was filtered, washed with $CH_2Cl_2$, 50% MeOH in $CHCl_3$, $CH_2Cl_2$ and ether in this order. The resin was dried under vacuum for 1 h. The unreacted amines on the resin were capped by reacting with Cap B Solution-1 (THF/1-Me-imidazole/Pyridine (8:1:1)) (97 mL) and Cap A Solution-1 (10vol% Ac2O/THF) (65 mL) on Bio-shaker (110 rpm) for 1 h at room temperature. The resin was filtered, washed with $CH_2Cl_2$, 20% MeOH in $CH_2Cl_2$, $CH_2Cl_2$ and ether in this order. The resin was dried under high vacuum afforded **S16** (1.80 g, loading: 229 μmol/g).

Synthesis of 4-(((2S,6R)-6-(4-benzamido-2-oxopyrimidin-1(2H)-yl)-4-tritylmorpholin-2-yl)methoxy)-4-oxobutanoic acid loaded onto aminomethylpolystyrene resin (**S18**)

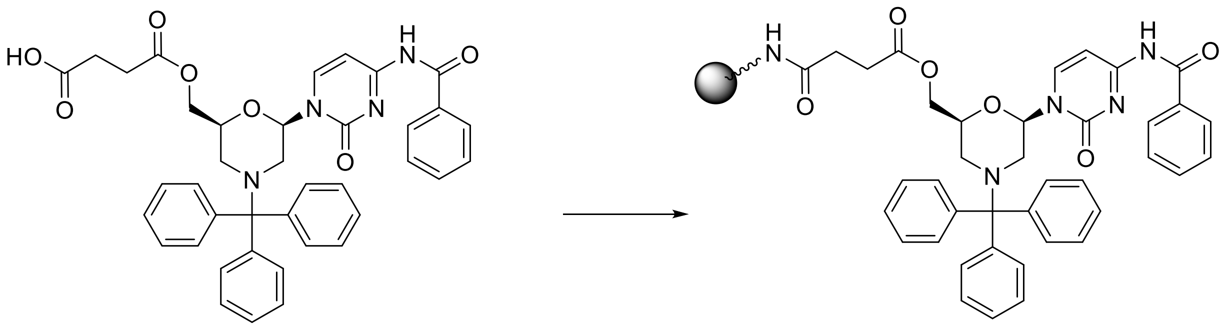

**S17** **S18**

4-(((2S,6R)-6-(4-Benzamido-2-oxopyrimidin-1(2H)-yl)-4-tritylmorpholin-2-yl)methoxy)-4-oxobutanoic acid (**S17**, CAS 1362664-31-0) (540 mg, 0.803 mmol) was dissolved in DMF (22 mL). HATU (1.03 g, 2.71 mmol) and DIPEA (0.701 mL, 4.01 mmol) were added. Aminomethyl Polystyrene Resin (Primer Support™ 5G Amino, 29-0999-92, manufactured by GE Healthcare) (2.32 g, amine content: 450 μmol/g) was added to the reaction mixture and gently shaken at room temperature on Bio-shaker (110 rpm) for 12 h. The resin was filtered, washed with $CH_2Cl_2$, 50% MeOH in $CHCl_3$, $CH_2Cl_2$ and ether in this order. The resin was dried under vacuum for 1 h. The unreacted amines on the resin were capped by reacting with Cap B Solution-1 (THF/1-Me-imidazole/Pyridine (8:1:1)) (127 mL) and Cap A Solution-1 (10vol% Ac2O/THF) (84 mL) on Bio-shaker (110 rpm) for 2 h at room temperature. The resin was filtered, washed with $CH_2Cl_2$, 20% MeOH in $CH_2Cl_2$, $CH_2Cl_2$ and ether in this order. The resin was dried under high vacuum to afford **S18** (2 g, loading: 194 μmol/g).

Synthesis of 4-(((2S,6R)-6-(6-benzamido-9H-purin-9-yl)-4-tritylmorpholin-2-yl)methoxy)-4-oxobutanoic acid loaded onto aminomethylpolystyrene resin (**S19**)

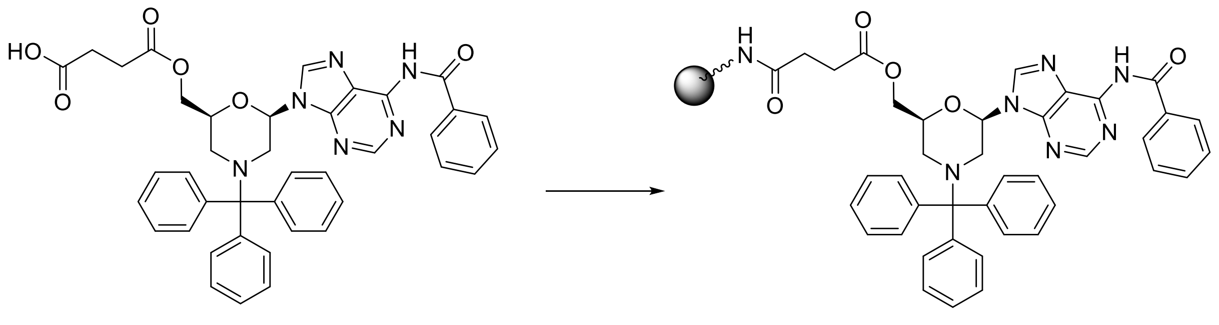

**S19** **S20**

4-(((2S,6R)-6-(6-Benzamido-9H-purin-9-yl)-4-tritylmorpholin-2-yl)methoxy)-4-oxobutanoic acid (**S19**, CAS 446206-67-2) (174 mg, 0.250 mmol) was dissolved in DMF (6.3 mL). HATU (321 mg, 0.845 mmol),DIPEA (0.218 mL, 1.25 mmol) and then Aminomethyl Polystyrene Resin (Primer Support™ 5G Amino, 29-0999-92, manufactured by GE Healthcare) (813 mg, amine content: 400 μmol/g) were added. The reaction mixture was gently shaken at room temperature on Bio-shaker (110 rpm) for 12 h. The resin was filtered, washed with $CH_2Cl_2$, 50% MeOH in $CHCl_3$, $CH_2Cl_2$ and ether in this order. The resin was dried under vacuum for 1 h. The unreacted amines on the resin were capped by reacting with Cap B Solution-1 (THF/1-Me-imidazole/Pyridine (8:1:1)) (39.4 mL) and Cap A Solution-1 (10vol% Ac2O/THF) (26.2 mL) on Bio-shaker (110 rpm) for 1 h at room temperature. The resin was filtered, washed with $CH_2Cl_2$, 20% MeOH in $CH_2Cl_2$, $CH_2Cl_2$ and ether in this order. The resin was dried under high vacuum to afford **S20** (827 mg, loading: 196 μmol/g).

<u>Synthesis of 4-(((2S,6R)-6-(6-(2-cyanoethoxy)-2-isobutyramido-9H-purin-9-yl)-4-tritylmorpholin-2-yl)methoxy)-4-oxobutanoic acid loaded onto aminomethylpolystyrene resin **(S22)**</u>

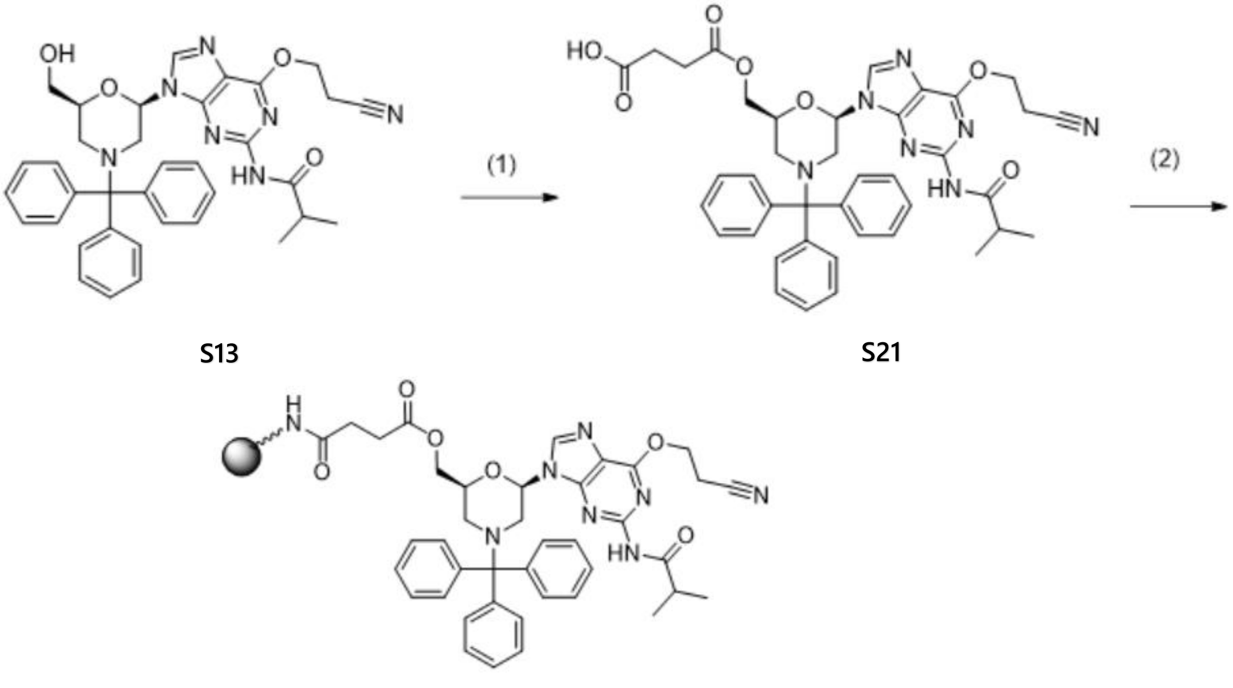

(1) 4-(((2S,6R)-6-(6-(2-cyanoethoxy)-2-isobutyramido-9H-purin-9-yl)-4-tritylmorpholin-2-yl)methoxy)-4-oxobutanoic acid (**S21**)

To a solution of N-(6-(2-cyanoethoxy)-9-((2R,6S)-6-(hydroxymethyl)-4-tritylmorpholin-2-yl)-9H-purin-2-yl)isobutyramide (**S13**, 1.50 g, 2.37 mmol) and DMAP (0.87 g, 7.12 mmol) in 1,2-Dichloroethane (15 mL) was added succinic anhydride (0.475 g, 4.75 mmol) at room temperature.. The reaction mixture was stirred for 1.5 h at 45 °C and cooled to room temperature. MeOH (5 mL) was added and the resulting mixure was concentrated in vacuo. EtOAc and 0.5M $KH_2PO_4$ aq (pH~7) were added to the residue, and the organic layer was separated. The aqueous layer was extracted with EtOAc. The combined organic layers were washed with 0.5M $KH_2PO_4$ aq (acidic), water and brine, dried over $MgSO_4$, filtered,and concentrated in vacuo to give the 4-(((2S,6R)-6-(6-(2-cyanoethoxy)-2-isobutyramido-9H-purin-9-yl)-4-tritylmorpholin-2-yl)methoxy)-4-oxobutanoic acid (**S21**, 1.51 g).

$^1$H NMR (396 MHz, CHLOROFORM-d) δ 9.22-9.36 (m, 1 H), 7.73-7.79 (m, 1 H), 7.43-7.54 (m, 5 H), 7.28-7.35 (m, 6 H), 7.15-7.23 (m, 4 H), 5.95-6.05 (m, 1 H), 4.71-4.88 (m, 2 H), 4.45-4.56 (m, 1 H), 4.30-4.39 (m, 1 H), 3.77-3.89 (m, 1 H), 3.38-3.46 (m, 1 H), 3.13-3.21 (m, 1 H), 2.97-3.09 (m, 2 H), 2.80-2.92 (m, 2 H), 2.47-2.67 (m, 4 H), 2.05-2.11 (m, 1 H), 1.23-1.30 (m, 6 H).
MS (ESI) m/z: $[M+H]^+$ Calcd for $C_{40}H_{42}N_7O_7$: 732.314; Found: 732.493.

(2) 4-(((2S,6R)-6-(6-(2-cyanoethoxy)-2-isobutyramido-9H-purin-9-yl)-4-tritylmorpholin-2-yl)methoxy)-4-oxobutanoic acid loaded onto aminomethylpolystyrene resin (**S22**)

4-(((2S,6R)-6-(6-(2-Cyanoethoxy)-2-isobutyramido-9H-purin-9-yl)-4-tritylmorpholin-2-yl)methoxy)-4-oxobutanoic acid (**S21**, 183 mg, 0.25 mmol) was dissolved in DMF (7.5 mL). HATU (321 mg, 0.845 mmol) ,DIPEA (0.218 mL, 1.25 mmol), and thenAminomethyl Polystyrene Resin (Primer Support$^{TM}$ 5G Amino, 29-0999-92, manufactured by GE Healthcare) (813 mg, amine content: 400 μmol/g) were added to the reaction mixture. The resulting mixture wasgently shaken at room temperature on Bio-shaker (110 rpm) for 18 h. The resin was filtered, washed with $CH_2Cl_2$, 50% MeOH in $CHCl_3$, $CH_2Cl_2$ and ether in this order. The resin was dried under vacuum for 1 h. The unreacted amines on the resin were capped by reacting with Cap B Solution-1 (THF/1-Me-imidazole/Pyridine (8:1:1)) (39.4 mL) and Cap A Solution-1 (10vol% $Ac_2O$/THF) (26.2 mL) on Bio-shaker (110 rpm) for 1 h at room temperature. The resin was filtered, washed with $CH_2Cl_2$, 20% MeOH in $CH_2Cl_2$, $CH_2Cl_2$ and ether in this order. The resin was dried under high vacuum to afford **S22** (750 mg, loading: 208 mol/g).

#### 2. Solid-Phase Synthesis of Stereorandom PMO-Gapmers

Oligonucleotides were synthesized on a NTS DNA/RNA synthesizer (NIHON TECHNO SERVICE) and a nS-8II synthesizer (GeneDesign). All syntheses were performed using an empty synthesis column of 1.0 µmol scale (Empty Synthesis Columns-TWIST, Glen Research) packed with a N-Tr-morpholino monomers loaded PrimerSupport (Primer Support$^{TM}$ 5G Amino, GE Healthcare, succinate linker).

Coupling of N-Tr-morpholino (PMO)-dimethylphosphoramidochloridate or 3'-DMT-DNA-5'-dimethylphosphoramidochloridate was performed by NTS DNA/RNA synthesizer. Dimethylphosphoramidochloridate reagents were prepared as 0.20 M solutions in 1,3-dimethyl-2-imidazolidinone (DMI), and 0.3 M solution of 1,2,2,6,6-Pentamethylpiperidine (PMP) in DMI was used as coupling activator. Detritylations were performed using 3% trichloroacetic acid (TCA) in $CH_2Cl_2$ and capping was done with Cap Mix A (THF/2,6-Lutidine/$Ac_2O$, Glen Research) and Cap Mix B (16% 1-Me-imidazole/THF, Glen Research). Neutrizations were performed using DIPEA in DMI and $CH_2Cl_2$. Remaining $Ac_2O$ in the solid support was removed by 0.4 M solution of morpholine in DMI. A stepwise description of the synthesis cycle is described in **Table S1**.

**Table S1**: Synthesis cycle for the coupling of PMO- or DNA-dimethylphosphoramidochloridate.

| Step | Reaction | Reagent | Time |
|---|---|---|---|
| 1 | $Ac_2O$ removal | Morpholine in DMI (0.4 M) | 540 sec |
| 2 | Wash | $CH_2Cl_2$ | |
| 3 | Detritylation | 3wt/v% TCA in $CH_2Cl_2$ | 40 sec |
| 4 | Wash | $CH_2Cl_2$ | |
| 5 | Neutrization | DIPEA in DMI and $CH_2Cl_2$ (10:45:45) | 120 sec |
| 6 | Wash | $CH_2Cl_2$ | |

| | | | |
|---|---|---|---|
| 7 | Coupling | Dimethylphosphoramidochloridate in DMI (0.2 M)<br>PMP in DMI (0.3 M)<br>(final concentration of dimethylphosphoramido-chloridate was 0.1 M) | 8 h |
| 8 | Wash | $CH_2Cl_2$ | |
| 9 | Capping | Cap Mix A (THF/2,6-Lutidine/$Ac_2O$)<br>Cap Mix B (16% 1-Me-imidazole/THF) | 60 sec |
| 10 | Wash | $CH_2Cl_2$ | |

Performed by NTS DNA/RNA synthesizer (Nihon-techno service)

Coupling of 3'-DMT-DNA-5'-cyanoethyl phosphoramidites and N-Tr-morpholino-5'-cyanoethyl phosphoramidites was performed by nS-8II synthesizer. The phosphoramidites were prepared as 0.20 M or 0.30 M solutions in $CH_3CN$ as shown in Table 2. A 0.40 M solution of 5-(Ethylthio)-1H-tetrazole (ETT) in $CH_3CN$ was used as coupling activator. Detritylations were performed using 3% trichloroacetic acid in $CH_2Cl_2$ and capping was done with Cap A Solution-1 (10vol% Ac2O/THF, WAKO) and Cap B Solution-1 (THF/1-Me-imidazole/Pyridine, (8:1:1, WAKO). Sulfurizations were carried out with 0.05 M solution of ((dimethylamino-methylidene)amino)-3H-1,2,4-dithiazoline-3-thione (DDTT) in pyridine and $CH_3CN$ (3:2). A stepwise description of the synthesis cycle is described in **Table S2**.

**Table S2**: Synthesis cycle for the coupling of DNA- or PMO-phosphoramidites.

| Step | Reaction | Reagent | Time |
|---|---|---|---|
| 1 | Detritylation | 3wt/v% TCA in $CH_2Cl_2$ | 20 sec |
| 2 | Coupling | DNA-amidites in $CH_3CN$ (0.20 M)<br>PMO-amidites in $CH_3CN$ (0.20 M for A,T,C and 0.3 M for G)<br>ETT in $CH_3CN$ (0.4 M)<br>(final concentration of amidites was 0.1M except for 0.15 M of PMO-G) | 5 min |
| 3 | Sulfurization | DDTT in pyridine and $CH_3CN$(3:2) (0.05 M) | 10 min |

| | | | |
|---|---|---|---|
| 4 | Capping | Cap A (10vol% acetic anhydride in THF)<br>Cap B (1-Me-imidazole/THF/Pyridine) | 30 sec |

Performed by nS-8II synthesizer (GeneDesign)

Cleavage and de-protection of oligonucleotides: after completion of the automated synthesis, the solid support was treated with 20 vol% diethylamine in $CH_3CN$ and then allowed to stand still for 1 h. The support was washed with anhydrous $CH_3CN$ and dried with argon. The support was transferred into empty screwcap tube and treated with a solution of 28% $NH_4OH$ and EtOH (3:1, 1 mL) at 60 °C for overnight. The support was filtered with Disc SyringeFilter (Hydrophilic PTEE, 0.45 µm, Shimadzu). The filtrate was dried with $N_2$ flow. The resultant residue was dissolved in water (Further filtration was performed when there was a suspension in the solution). The crude material was analyzed by reverse-phase high-performance liquid chromatography (RP-HPLC) and liquid chromatography mass spectrometry (LCMS).

Purification of N-Tr: the crude material was purified by RP-HPLC with purification condition-1 (small scale) or condition-2 (medium scale). The obtained fractions were collected and dried with $N_2$ flow.

Purification Condition-1:

Column: XBridge BEH C18 OBD prep (10 x 150 mm, Particle size 5 µm, Waters)

Detection: 260 nm

Column temperature: 55 °C

Eluent A: 100 mM HFIP, 8.6 mM TEA / water

Eluent B: 100% MeOH

Gradient B: 25% to 56% in 25 min

Flow rate: 3.5 mL/min

Purification Condition-2:

Column: XBridge BEH Prep C18 OBD (19 x 150 mm, Particle size 5 µm, Waters)

Detection: 260 nm

Column temperature: 55 °C

Eluent A: 100 mM HFIP, 8.6 mM TEA / water

Eluent B: 100% MeOH

Gradient B: 10% to 70% in 20 min

Flow rate: 20 mL/min

Detritylation and purification (for in vitro/in vivo): The solution for detritylation was prepared by mixing TFA (0.17 mL), $Et_3N$ (0.16 mL), EtOH (0.25 mL), 2,2,2-trifluoroethanol (2.5 mL) and $CH_2Cl_2$ (22.25 mL). To the residue of purified N-Tr was added the above solution (excess amount) at 0 °C. After several hours at 0 °C, 5% DIPEA in $CH_2Cl_2$ was added to the mixture for neutrization. Then the mixture was dried by $N_2$ flow. The residue was dissolved by water and purified by RP-HPLC with purification condition-1 using gradient B: 25% to 35% in 25 min. The obtained fractions were collected and dried with $N_2$ flow.

Desalting of oligonucleotides (for in vitro study): The purified oligonucleotides after detritylation was diluted with water to 2.5 mL of total volume and then desalted by Illustra™ NAP™-25 Columns (GE Healthcare) using water as an equilibration buffer according to the manufacturer's protocol. The obtained solution were dried with $N_2$ flow.

Ion-exchange of oligonucleotides (for in vivo study): the purified oligonucleotides after detritylation were diluted with start buffer (0.02 M Na phosphate buffer (pH 8.0), 20% CH3CN) until the total volume became 1 mL. Anion-exchange was carried out by HiTrapQ HP (1 mL, GE Healthcare) following the manufacturer's protocol using the start buffer and elution buffer (start buffer with 1.5 M NaCl). The obtained fractions were collected and dried with N2 flow. The residue was diluted with water to 2.5 mL of total volume and then desalted by Illustra™ NAP™-25 Columns (GE Healthcare) using water as an equilibration buffer according to the manufacturer's protocol. The obtained solution were dried with N2 flow.

Ion-exchange of oligonucleotides (for in vivo-2): anion-exchange was carried out by using centrifugal spin filters (Vivaspin 20, 3,000 molecular weight cut-off, GE Healthcare). The purified oligonucleotides after detritylation were dissolved with NaOAc (0.1 M) up to 14 mL of total volumn and then the solution was applied to the spin filter. The sample was concentrated to less than 5 mL with centrifuge. The concenrated solution was diluted with water up to 14 mL of total volume and concentrated to less than 5 mL. This dilution and concentration process was repeated twice. The residue was transferred to empty tube and concentrated with the vacuum concentrator.

Analysis: the obtained residue was dissolved with water and the concentration was determined by the absorbance at 260 nm (measured with Nanodrop) and the factor value (ng・cm/μL).

##### 3. MALDI-MASS Analysis of stereorandom PMO-gapmers

MALDI-MASS analysis was conducted for seventeen 5-8-5 PMO-gapmers and twelve 4-10-4 gapmers, with the results shown in **Table S3** and **Table S4**, respectively. MASS spectra were obtained by negative mode on Autoflex MALDI-TOF-MS spectrometer calibrated by standard oligonucleotide (Bruker). 3-Hydroxypicolinic acid with the addition of Diammonium hydrogen citrate was used as matrix.

**Table S3** – MALDI-MASS for 5-8-5 PMO-Gapmers

| Compound No. (SEQ ID NO:) | Sequence (5'-3') | | |
|---|---|---|---|
| | | Theoretical | Found |
| **ASO-369** (SEQ ID NO: 1) | GGGGACTCGCTGACATGG | 5942.9 | 5944.0 |
| **ASO-373** (SEQ ID NO: 2) | TGGGTGTAGCGAGAATCC | 5942.0 | 5943.8 |
| **ASO-380** (SEQ ID NO: 3) | GGGTGCACTAGTTTATAG | 5931.9 | 5933.7 |
| **ASO-388** (SEQ ID NO: 4) | GGGGTCTTCTAATATCCT | 5842.9 | 5844.9 |

| | | | |
|---|---|---|---|
| **ASO-389** (SEQ ID NO: 5) | AGGTTCTCGCTATATCGC | 5827.9 | 5828.6 |
| **ASO-401** (SEQ ID NO: 6) | GAGTTAGAAGCTTTGACT | 5915.9 | 5916.0 |
| **ASO-409** (SEQ ID NO: 7) | GCAGATGACCCTTAGACA | 5854.9 | 5857.3 |
| **ASO-413** (SEQ ID NO: 8) | CAAACCTGTCACACCCGA | 5759.9 | 5759.7 |
| **ASO-417** (SEQ ID NO: 9) | TTAAACCCCATAGACATA | 5797.9 | 5800.8 |
| **ASO-418** (SEQ ID NO: 10) | GAGGCCCAAATGATCACA | 5863.9 | 5865.9 |
| **ASO-463** (SEQ ID NO: 11) | TGGATTTAGCAGTAGGGT | 5972.0 | 5972.8 |
| **ASO-467** (SEQ ID NO: 12) | AGCAGATGACCCTTAGAC | 5854.9 | 5853.3 |
| **ASO-468** (SEQ ID NO: 13) | AGCCGGCATACAGTATAT | 5869.9 | 5869.8 |
| **ASO-469** (SEQ ID NO: 14) | TGTGCTCTTTATGGATGG | 5913.9 | 5914.7 |
| **ASO-470** (SEQ ID NO: 15) | GGATTTAGCAGTAGGGTG | 5997.0 | 5996.2 |
| **ASO-473** (SEQ ID NO: 16) | CCCCATGACTACAGTGTG | 5821.9 | 5822.1 |
| **ASO-474** (SEQ ID NO: 17) | GCTTTTGTGACCAGGGAC | 5892.9 | 5892.7 |

**Table S4** – MALDI-MASS for 4-10-4 PMO-Gapmers

| Compound No. (SEQ ID NO:) | Sequence (5'-3') | | |
|---|---|---|---|
| | | Theoretical | Found |
| **ASO-483** (SEQ ID NO: 11) | TGGATTTAGCAGTAGGGT | 5951.9 | 5952.9 |
| **ASO-484** (SEQ ID NO: 17) | GCTTTTGTGACCAGGGAC | 5872.9 | 5873.9 |
| **ASO-485** (SEQ ID NO: 5) | AGGTTCTCGCTATATCGC | 5807.8 | 5809.0 |
| **ASO-486** (SEQ ID NO: 12) | AGCAGATGACCCTTAGAC | 5834.9 | 5837.1 |
| **ASO-487** (SEQ ID NO: 14) | TGTGCTCTTTATGGATGG | 5893.9 | 5897.6 |
| **ASO-488** (SEQ ID NO: 16) | CCCCATGACTACAGTGTG | 5801.8 | 5803.9 |
| **ASO-489** (SEQ ID NO: 2) | TGGGTGTAGCGAGAATCC | 5921.9 | 5921.7 |
| **ASO-490** (SEQ ID NO: 3) | GGGTGCACTAGTTTATAG | 5911.9 | 5911.4 |
| **ASO-491** (SEQ ID NO: 4) | GGGGTCTTCTAATATCCT | 5822.8 | 5822.2 |
| **ASO-492** (SEQ ID NO: 7) | GCAGATGACCCTTAGACA | 5834.9 | 5835.6 |
| **ASO-493** (SEQ ID NO: 10) | GAGGCCCAAATGATCACA | 5843.9 | 5844.1 |
| **ASO-494** (SEQ ID NO: 9) | TTAAACCCCATAGACATA | 5777.8 | 5778.7 |

#### 4. Solution-Phase Synthesis of Stereodefined PMO-Gapmers

The stereochemistry of the phosphorus atoms in the phosphorothioate linkages between the deoxyribonucleosides of the PMO-gamers were controlled by using similar methods as those disclosed by Knouse and deGruyter *et al*. (see Knouse, K. and deGrutyer, J. *et al*, "Unlocking P(V): Reagents for chiral phosphorothioate synthesis", *Science*, 2018, 361(6408): 1234-1238) and Stec *et al* (see Stec *et al*, "Deoxyribonucleoside 3¢-*O*-(2-Thio-and 2-Oxo-"spiro"-4,4-pentamethylene-1,3,2-oxathiaphospholane)s: Monomers for Stereocontrolled Synthesis of Oligo(deoxyribonucleoside phosphorothioate)s and Chimeric PS/PO Oligonucleotides", *J. Am. Chem. Soc.* 1998**,** *120,* 7156-7167; Karwowski and Stec *et al*, "Stereocontrolled synthesis of LNA Dinucleoside phosphorothioate by the oxathiaphospholane approach", *Bioorg. Med. Chem. Lett.*, 11 (2001) 1001–1003; and Karwowski and Stec *et al*, "Nucleoside 3′-*O*-(2-Oxo-"*Spiro*"-4.4-Pentamethylene-1.3.2-Oxathiaphospholane)S: Monomers For Stereocontrolled Synthesis Of Oligo(Nucleoside Phosphorothioate/Phosphate)S", *Nucleosides & Nucleotides*, 17(9-1l), 1747-1759 (1998)), which are herein fully incorporated by reference.

The solution phase synthesis of stereodefined PMO-gapmers presented within this example differs from previous solution phase syntheses of antisense oligonucleotides in that the present synthesis utilizes a 5 + 13 coupling step. Prior solution phase sytheses typically couple one nucleotide at a time until the final product is formed; however, these coupling methods lead to an increased chance that the final product will be contaminated with other species of oligonucleotides of varying lengths. This increased chance of contamination is due to the occurrence of not all of the oligonucleotides having enough time to interact with the next nuceleotide added into the solution. Therefore, not only does the final product have an increased chance of containing nucleotides of varying lengths, but also varying nucleotide sequences.

An advantage of performing a 5 + 13 coupling is that it lowers the number of steps where one nucleotide is added at a time before formation of the final product, hence potentially leading to final products with increased purity and yields.

The following example reports the preparation of a stereodefined 4-10-4 gapmer (**ASO-486):**

| Compound No. (SEQ ID NO:) | Sequence (5'-3') |
|---|---|
| **ASO-486** (SEQ ID NO: 12) | AGCAGATGA$^{m}$C $^{m}$C $^{m}$C TTAGAC<br>$^{m}$C: 5-MethylCytosine |

The synthesized gapmer has a chirality represented herein as:

SSSSSSSRSSSSSSSSS (**ASO-486-R5-S**), SSSRSSSRSSSSSSSSS (**ASO-486-R5-R**) or SSSMSSSRSSSSSSSSS (**ASO-486-R5-M**)

"M" means a mixture of R configuration and S configuration.

With the benefit of this specification, including the other examples presented herein, a person of skill in the art would recognize that gapmers with the same sequence but different chirality could be prepared with reference to the chirality of the added reagents in the coupling steps.

**Preparation of Compound ASO-486-R5-R**

**Preparation of 5'-PMO wing**

<u>2-mer of 5'-PMO: coupling</u>

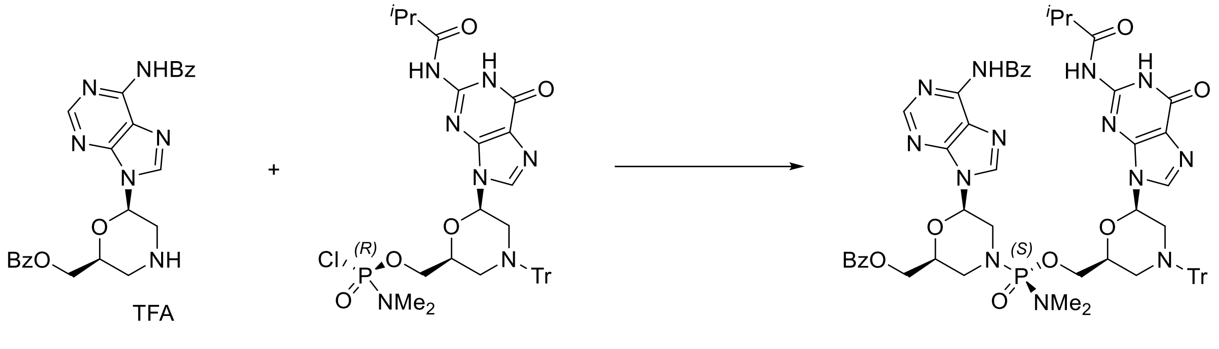

**S23** **G'2** **S24**

To a solution of starting material **S23** (1.00 g, 1.42 mmol) in 1,3-Dimethyl-2-imidazolidinone (10 mL) were added reactant **G'2** (0.854 g, 1.491 mmol) and 1,2,2,6,6-pentamethylpiperidine (1.03 mL, 5.68 mmol) at -room temperature. The reaction solution was stirred overnight and treated with THF (10 mL) followed by MTBE (100 mL) and n-heptane (100 mL). The supernatant was decanted/filtered and the sticky stuff was rinsed with a mixture of THF/MTBE/n-heptane (20 mL/100 mL/100 mL). The leftover material was dissolved in $CH_2Cl_2$ and purified on silica gel column chromatography with a gradient of 0% to 20% MeOH in EtOAc to afford target compound **S24** (1.33 g).

MS (ESI) m/z: $[M+H]^+$ Calcd for $C_{59}H_{61}N_{13}O_9P$ 1126.44; Found 1126.29.

<u>2-mer of 5'-PMO: deprotection</u>

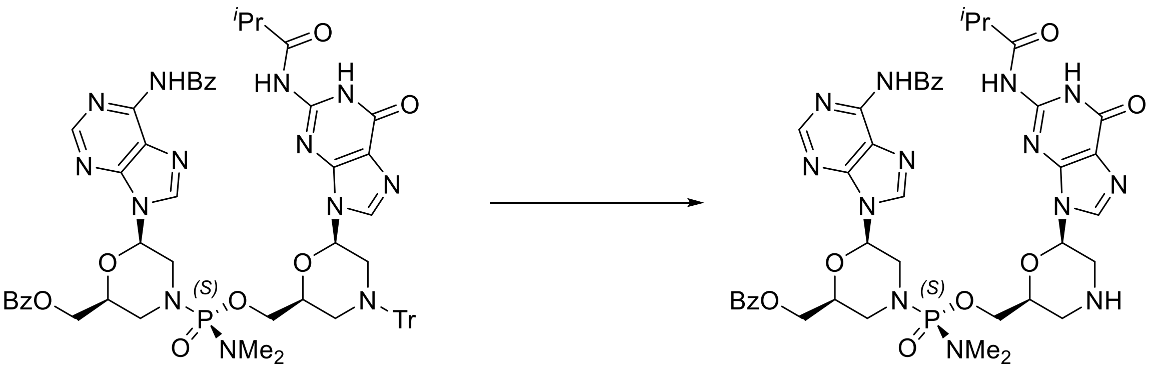

**S24** **S25**

To a flask charged with starting material **S24** (1.33 g, 1.18 mmol) was added ethanol (0.690 mL, 11.8 mmol) followed by a solution of TFA (0.364 mL, 4.72 mmol) in $CH_2Cl_2$ (20 mL) at room temperature. The reaction solution was stirred for 25 min and treated with EtOAc (7.5 mL) followed by *n*-heptane (40 mL). The slurry was filtered and the cake was rinsed with a mixture of $CH_2Cl_2$ (15 mL), EtOAc (7.5 mL) and *n*-heptane (40 mL). The TFA salt was then redissolved in $CH_2Cl_2$ (20 mL) at room temperature, and 1,2,2,6,6-pentamethylpiperidine (2.14 mL, 11.8 mmol) was added. The reaction mixture was stirred for 5-10 min before *n*-heptane (100 mL) was added. The slurry was sonicated to break down any aggregated pieces, and then filtered. The cake was rinsed with a mixture of $CH_2Cl_2$ (20 mL) and *n*-heptane (100 mL) to afford target compound **S25** (0.93 g).

MS (ESI) m/z: $[M+H]^+$ Calcd for $C_{40}H_{47}N_{13}O_9P$ 884.34 ; Found 884.26.

3-mer of 5'-PMO: coupling

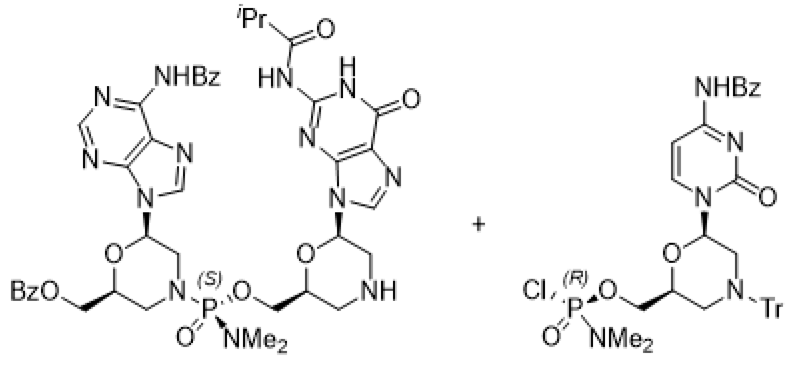

S25 C1

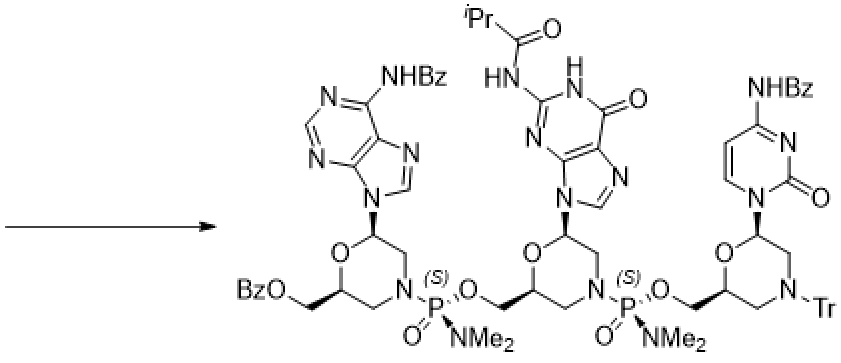

S26

To a solution of starting material **S25** (0.930 g, 1.05 mmol) in 1,3-dimethyl-2-imidazolidinone (9.24 mL) was added 1,2,2,6,6-pentamethylpiperidine (0.571 mL, 3.16 mmol) followed by reactant **C1** (0.918 g, 1.32 mmol) at room temperature. The reaction solution was stirred overnight and treated with EtOAc (10 mL) followed by MTBE (150 mL) and *n*-heptane (50 mL). The slurry was filtered and the cake was rinsed with a mixture of EtOAc (10 mL), MTBE (75 mL) and *n*-heptane (25 mL) to afford target compound **S26** (1.70 g).

MS (ESI) m/z: $[M+H]^+$ Calcd for $C_{77}H_{83}N_{18}O_{14}P_2$ 1545.58 ; Found 1545.58.

3-mer of 5’-PMO: deprotection

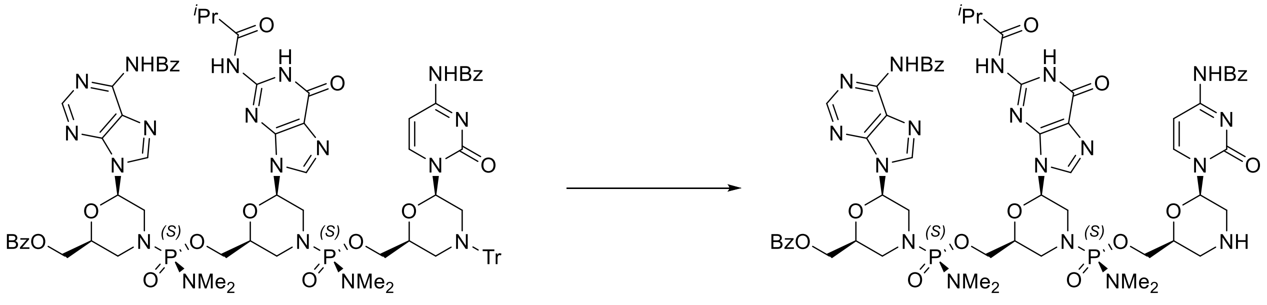

**S26** **S27**

To a flask charged with starting material **S26** (1.70 g, 1.10 mmol) was added ethanol (0.642 mL, 11.0 mmol) followed by a solution of TFA (0.339 mL, 4.40 mmol) in $CH_2Cl_2$ (25.5 mL) at room temperature. The reaction solution was stirred for 1 h and treated with EtOAc (12.5 mL) followed by *n*-heptane (45 mL). The slurry was filtered and the cake was rinsed with a mixture of $CH_2Cl_2$ (25 mL), EtOAc (12.5 mL) and *n*-heptane (40 mL). The TFA salt was then dissolved in $CH_2Cl_2$ (25.5 mL) at room temperature, and 1,2,2,6,6-pentamethylpiperidine (1.99 mL, 11.0 mmol) was added. The reaction solution was stirred for ca. 10 min and treated with EtOAc (12.5 mL) followed by MTBE (70 mL). The slurry was then filtered and the cake was rinsed with a mixture of $CH_2Cl_2$ (25.5 mL), EtOAc (12.5 mL) and MTBE (70 mL) to afford target compound **S27** (1.19 g).

MS (ESI) m/z: $[M+H]^+$ Calcd for $C_{58}H_{69}N_{18}O_{14}P_2$ 1303.47; Found 1303.45.

4-mer of 5’-PMO: coupling

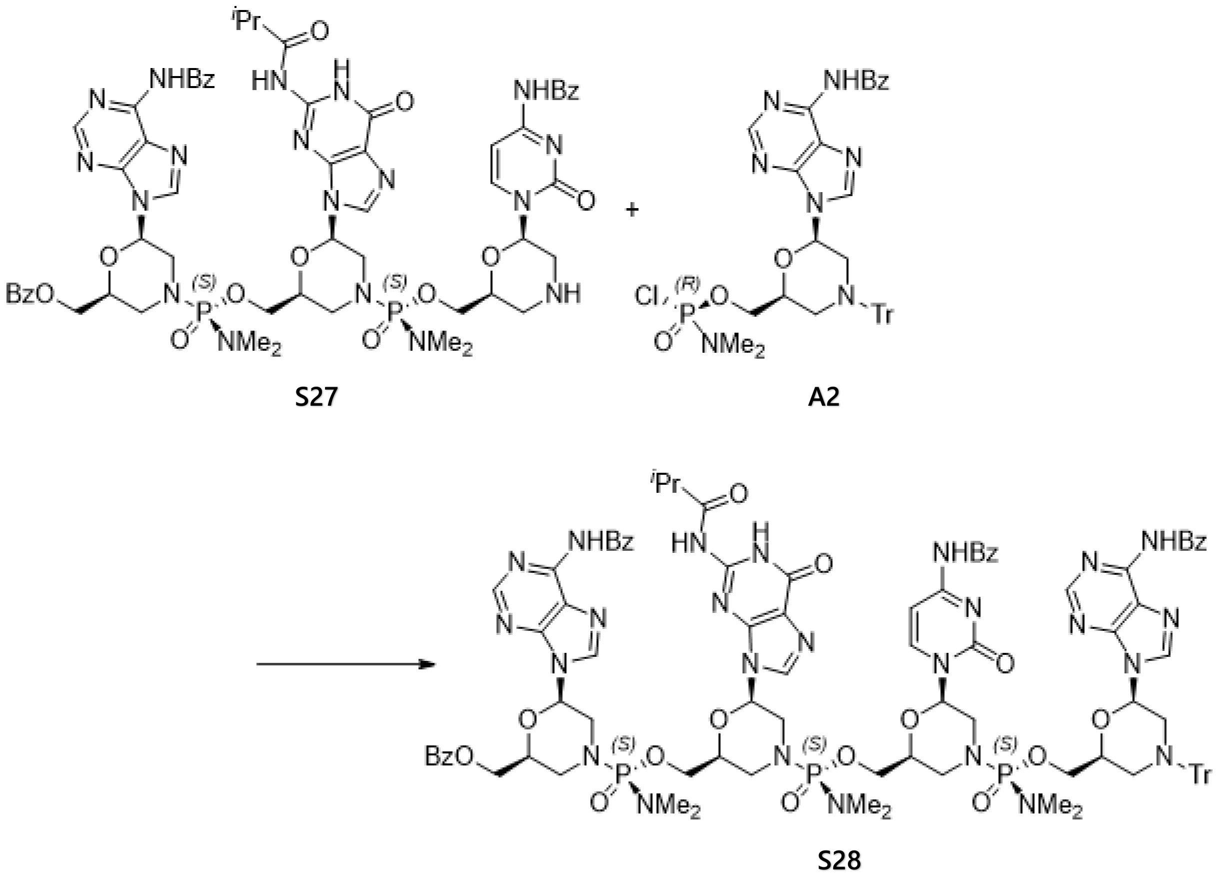

To a solution of staring material **S27** (1.19 g, 0.913 mmol) in 1,3-dimethyl-2-imidazolidinone (8.0 mL) was added 1,2,2,6,6-pentamethylpiperidine (0.496 mL, 2.74 mmol) followed by reactant **A2** (0.824 g, 1.14 mmol) at room temperature. The reaction solution was stirred overnight and treated with EtOAc (8 mL) followed by MTBE (100 mL). The slurry was filtered and rinsed with a mixture of EtOAc (16 mL) and MTBE (100 mL) to afford target compound **S28** (2.04 g).

MS (ESI) m/z: $[M+H]^+$ Calcd for $C_{96}H_{105}N_{25}O_{18}P_3$ 1988.73; Found 1988.67.

4-mer of 5’-PMO: deprotection

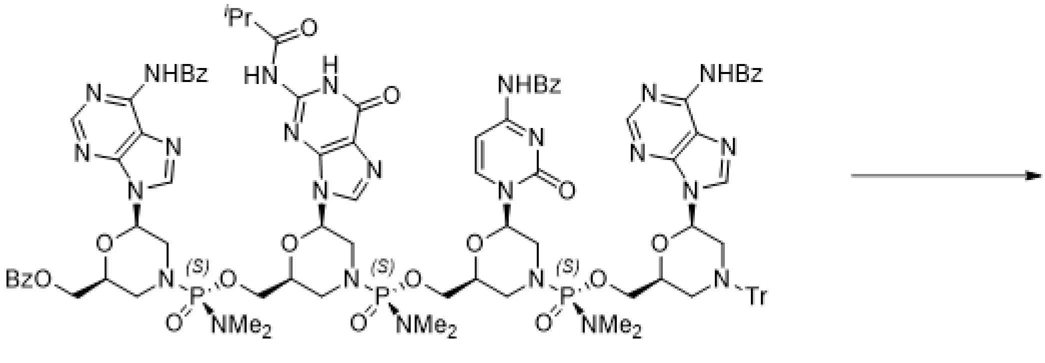

**S28**

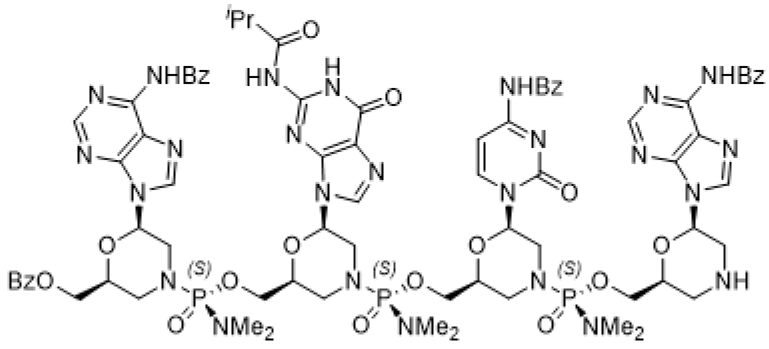

**S29**

To a flask charged with starting material **S28** (2.04 g, 1.03 mmol) was added ethanol (0.599 mL, 10.3 mmol) followed by a solution of TFA (0.474 mL, 6.15 mmol) in $CH_2Cl_2$ (24 mL) at room temperature. The reaction solution was stirred for 1.5 h and treated with EtOAc (12 mL) followed by *n*-heptane (40 mL). The slurry was filtered and the cake was rinsed with a mixture of $CH_2Cl_2$ (24 mL), EtOAc (12 mL) and *n*-heptane (40 mL). The TFA salt was then dissolved in $CH_2Cl_2$ (23.8 mL), and treated with 1,2,2,6,6-pentamethylpiperidine (1.856 mL, 10.26 mmol) for ca. 10 min before EtOAc (48 mL) was added followed by addition of MTBE (48 mL). The slurry was filtered and rinsed with a mixture of $CH_2Cl_2$ (24 mL), EtOAc (48 mL) and MTBE (48 mL) to afford target compound **S29** (1.50 g).

MS (ESI) m/z: $[M+H]^+$ Calcd for $C_{77}H_{91}N_{25}O_{18}P_3$ 1746.62; Found 1746.51.

5-mer of 5' PMO: coupling

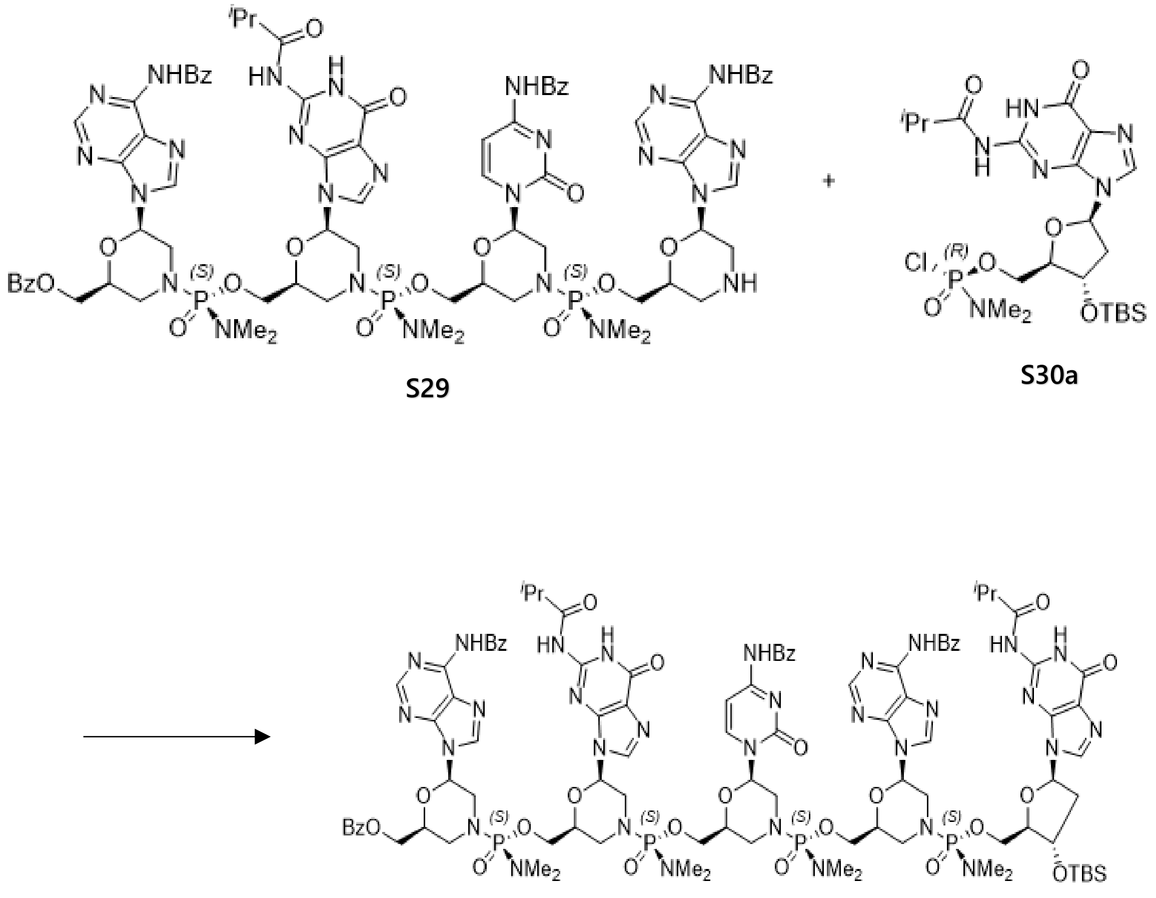

To a solution of starting material **S29** (500 mg, 0.286 mmol) in 1,3-dimethyl-2-imidazolidinone (7.5 mL) was added 1,2,2,6,6-pentamethylpiperidine (0.16 mL, 0.86 mmol) followed by reactant **S30a** (206 mg, 0.358 mmol) (synthesized according to the process undermentioned) at room temperature. The reaction solution was stirred overnight and treated with EtOAc (7.5 mL) followed by MTBE (100 mL). The slurry was filtered and rinsed with a mixture of EtOAc (15 mL) and MTBE (100 mL) to give target compound **S31** (710 mg).

$^{31}$P NMR (162 MHz, METHANOL-d4) δ ppm 17.42 (s, 1 P), 17.07 (s, 1 P), 17.02 (s, 1 P), 16.82 (s, 1 P).

MS (ESI) m/z: $[M+2H]^{2+}$ Calcd for $C_{99}H_{129}N_{31}O_{24}P_4Si$ 1143.93; Found 1144.03.

5-mer of 5’ PMO: deprotection

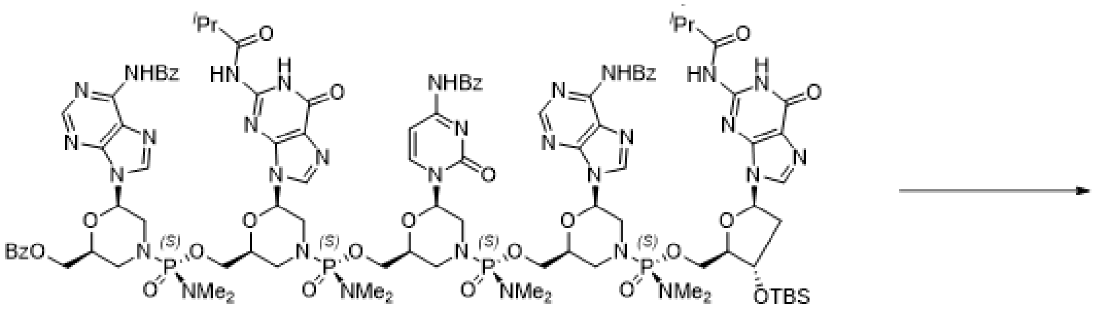

**S31**

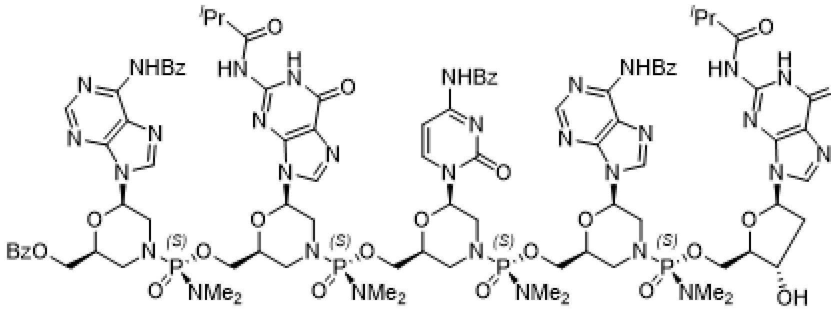

**S32**

To a flask charged with starting material **S31** (710 mg, 0.31 mmol) at room temperature was added pyridine (5.90 mL, 73.0 mmol), triethylamine (5.93 mL, 42.5 mmol) and $CH_2Cl_2$ (5.9 mL). The solution was then treated with triethylamine trihydrofluoride (759 µL, 4.66 mmol). The reaction solution was stirred overnight, cooled in an ice bath, and then treated with methoxytrimethylsilane (2.95 ml, 21.4 mmol). The mixture was stirred with ice bath for 1 h and treated with 1,3-dimethyl-2-imidazolidinone (5.9 mL) followed by EtOAc (100 mL) and MTBE (50 mL). The slurry was filtered and rinsed with a mixture of $CH_2Cl_2$ (5.9 mL), EtOAc (118 mL) and MTBE (50 mL) to afford target compound **S32** (627 mg).

$^{31}$P NMR (162 MHz, CHLOROFORM-d) δ ppm 17.37 (s, 1P), 17.08 (s, 1P), 17.03 (s, 1P), 16.82 (s, 1P).

MS (ESI) m/z: $[M+2H]^{2+}$ Calcd for $C_{93}H_{115}N_{31}O_{24}P_4$ 1087.39; Found 1087.17.

5-mer of 5' PMO: activation with (-)-PSI

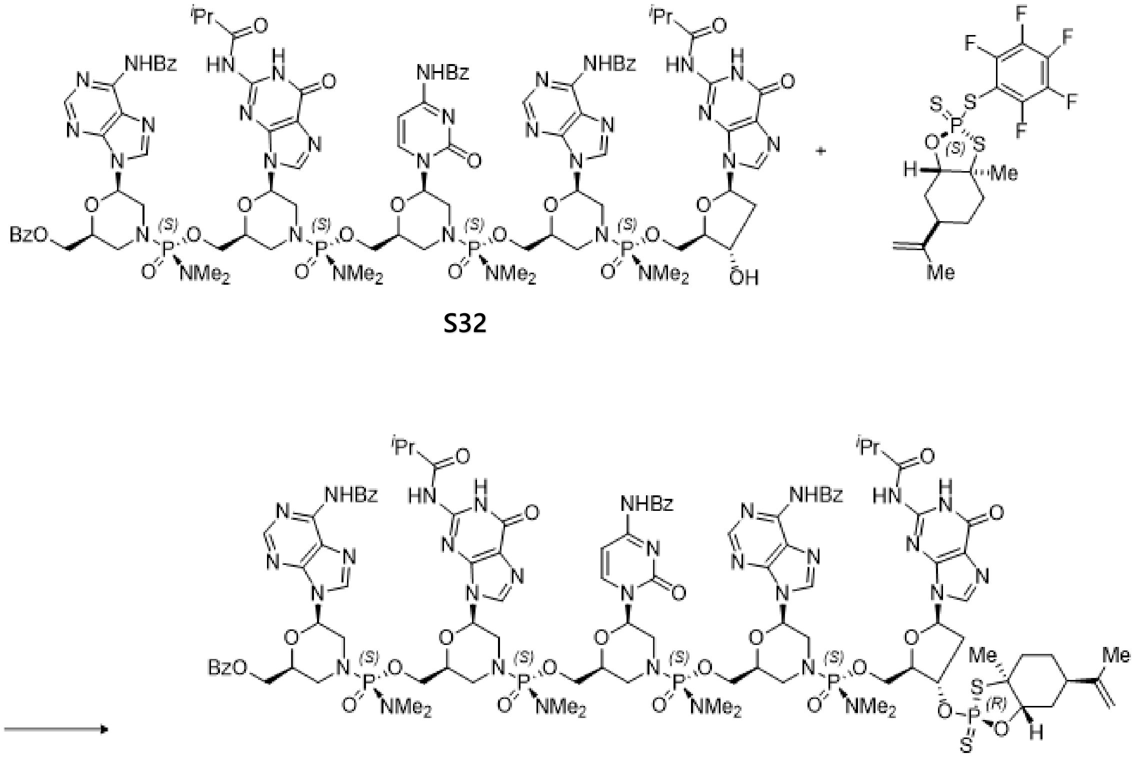

S32 → S33

To a solution of starting material **S32** (510 mg, 0.235 mmol) in a mixture of $CH_2Cl_2$ (21.9 mL), THF (7.1 mL) and 1,3-dimethyl-2-imidazolidinone (1.7 mL) was added (-)-PSI (Aldrich, CAS: 2245335-70-8, 194 mg, 0.434 mmol) at room temperature followed by activated 4Å molecular sieves (2.5 g). The mixture was stirred for 50 min and treated dropwise with a solution of DBU (49.5 µL, 0.329 mmol) in $CH_2Cl_2$ (0.872 mL). The reaction mixture was then stirred for 30 min. The precipitate was filtered and the cake was rinsed with a mixture of $CH_2Cl_2$ (43.6 mL), THF (14.2 mL) and 1,3-dimethyl-2-imidazolidinone (3.5 mL). The filtrate was treated with MTBE (218 mL), the resulting precipitate was filtered, and the cake was rinsed with a mixture of $CH_2Cl_2$ (31.8 mL), THF (10.6 mL) and MTBE (100 mL) to afford target product **S33** (548 mg).

$^{31}$P NMR (162 MHz, $CD_2Cl_2$) δ ppm 101.46 (s, 1P), 16.74 (s, 1P), 16.46 (s, 1P), 16.32 (s, 1P), 16.13 (s, 1P).

MS (ESI) m/z: $[M+2H]^{2+}$ Calcd for $C_{103}H_{130}N_{31}O_{25}P_5S_2$ 1210.40; Found 1210.09.

Synthesis of Compounds **S30a** and **S30b**

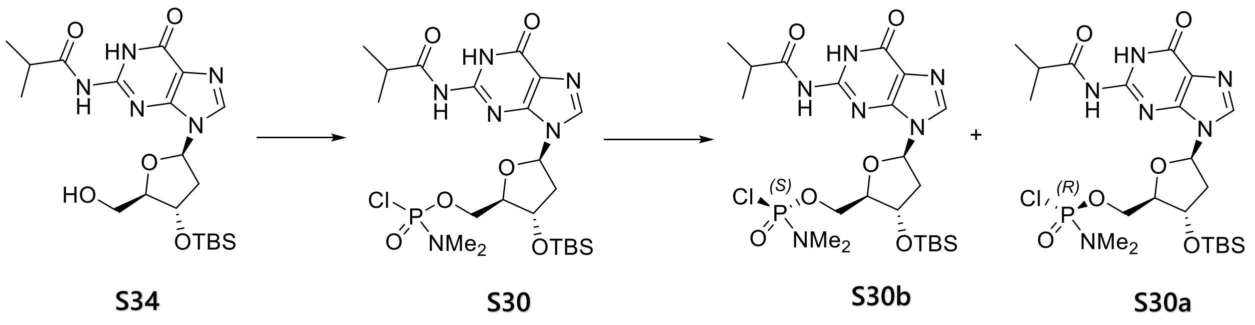

To a solution of N-(9-((2R,4S,5R)-4-((tert-butyldimethylsilyl)oxy)-5-(hydroxymethyl)tetrahydrofuran-2-yl)-6-oxo-6,9-dihydro-1H-purin-2-yl)isobutyramide **S34** (2.76 g, 6.11 mmol) in acetonitrile (40 mL) and $CH_2Cl_2$ (40 mL) were added DBU (3.04 mL, 20.2 mmol) and LiBr (1.75 g, 20.2 mmol) followed by dimethylphosphoramidic dichloride (1.16 mL, 9.78 mmol) at 0 ºC. The reaction solution was stirred at 0 ºC for 1 h and then quenched with 10% aqueous citric acid (77 mL). The mixture was extracted two times with $CH_2Cl_2$ (200 mL each time). The combined organic layers were subsequently washed twice with water and 15 wt% NaCl aqueous solution, dried over $Na_2SO_4$, and concentrated in vacuo. Biotage purification with a gradient of 90% to 100% EtOAc in *n*-heptane afforded target product **S30** (1.91 g) as a mixture of two diastereomers **S30a** and **S30b**. The mixture of two diastereomers was subjected to prep. HPLC separation to afford **S30b** (444 mg) and **S30a** (304 mg).

*HPLC Conditions for separation*

Column: Chiralpak IA, 21 x 250mm, 5 µ

Flowrate: 20 mL/min

Mobile Phase: 100% EtOAc

Gradient: Isocratic

Runtime 20 mins

Injection Volume: 500uL 150mg/ml concentration

Detection: 254nm

Peak1 (Rt 9.3 min)

((2R,3S,5R)-3-((tert-butyldimethylsilyl)oxy)-5-(2-isobutyramido-6-oxo-1,6-dihydro-9H-purin-9-yl)tetrahydrofuran-2-yl)methyl (S)-dimethylphosphoramidochloridate (**S30b**):

$^1$H NMR (400 MHz, CHLOROFORM-d) δ = 12.19 (br s, 1H), 9.93 (br s, 1H), 7.76 (br s, 1H), 6.25 (br t, *J* = 7.3 Hz, 1H), 4.98 - 4.90 (m, 1H), 4.67 (br d, *J* = 4.3 Hz, 1H), 4.39 - 4.26 (m, 2H), 3.08 - 2.99 (m, 1H), 2.82 - 2.73 (m, 1H), 2.73 (s, 3H), 2.69 (s, 3H), 2.28 (br dd, *J* = 5.9, 13.5 Hz, 1H), 1.26 (d, *J* = 6.9 Hz, 3H), 1.22 (d, *J* = 6.8 Hz, 3H), 0.93 (s, 9H), 0.14 (s, 3H), 0.14 (s, 3H).

$^{31}$P NMR (162 MHz, CHLOROFORM-d) δ ppm 20.39 (s, 1P).

MS (ESI) m/z: $[M+H]^+$ Calcd for $C_{22}H_{39}ClN_6O_6PSi$ 577.21; Found 577.07.

Peak2 (Rt 15.3 min)rt

((2R,3S,5R)-3-((tert-butyldimethylsilyl)oxy)-5-(2-isobutyramido-6-oxo-1,6-dihydro-9H-purin-9-yl)tetrahydrofuran-2-yl)methyl (R)-dimethylphosphoramidochloridate (**S30a**).

$^1$H NMR (400 MHz, CHLOROFORM-d) δ = 12.24 (br s, 1H), 10.34 (br s, 1H), 7.88 (br s, 1H), 6.27 (br t, *J* = 6.8 Hz, 1H), 5.27 - 5.13 (m, 1H), 4.91 - 4.85 (m, 1H), 4.37 - 4.26 (m, 1H), 4.15 - 4.07 (m, 1H), 3.24 - 3.16 (m, 1H), 2.80 (s, 3H), 2.76 (s, 3H), 2.75 - 2.71 (m, 1H), 2.37 (br dd, *J* = 6.9, 12.1 Hz, 1H), 1.25 (d, *J* = 6.8 Hz, 3H), 1.24 (d, *J* = 6.8 Hz, 3H), 0.92 (s, 9H), 0.12 (s, 3H), 0.12 (s, 3H)

$^{31}$P NMR (162 MHz, CHLOROFORM-d) δ ppm 19.67 (s, 1P).

MS (ESI) m/z: $[M+H]^+$ Calcd for $C_{22}H_{39}ClN_6O_6PSi$ 577.21; Found 577.07.

**Preparation of 3'-PMO wing**

<u>2-mer of 3'-PMO: coupling</u>

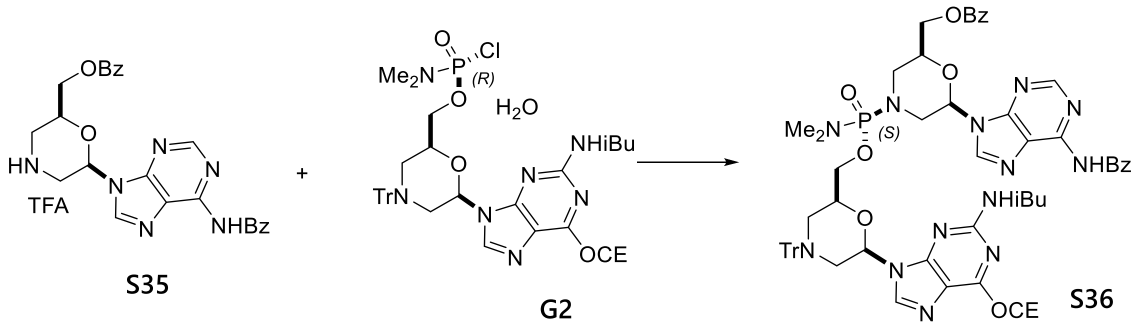

To a solution of starting material **S35** (1.33 g, 2.32 mmol) in THF (16 mL) was added 1,2,2,6,6-pentamethylpiperidine (1.15 mL, 6.34 mmol). The resulting solution was cooled to 0 ºC and treated with reactant **G2** (1.60 g, 2.11 mmol). The reaction mixture was warmed to room temperature and stirred overnight. A saturated $NaHCO_3$ solution (25 mL) and water (10 mL) were added, and the resulting mixture was extracted with $CH_2Cl_2$ (40 mL each) three times. The combined organic layers were washed with 30 wt% NaCl aqueous solution (20 mL), dried over $MgSO_4$, filtered, and concentrated in vacuo. The residue was purified by silica gel column chromatography. Elution with 3-15% MeOH in EtOAc afforded 2.316 g of target product **S36**.
MS (ESI) m/z: $[M+H]^+$ Calcd for $C_{62}H_{64}N_{14}O_9P$ 1179.47; Found 1179.41.

<u>2-mer of 3'-PMO: deprotection</u>

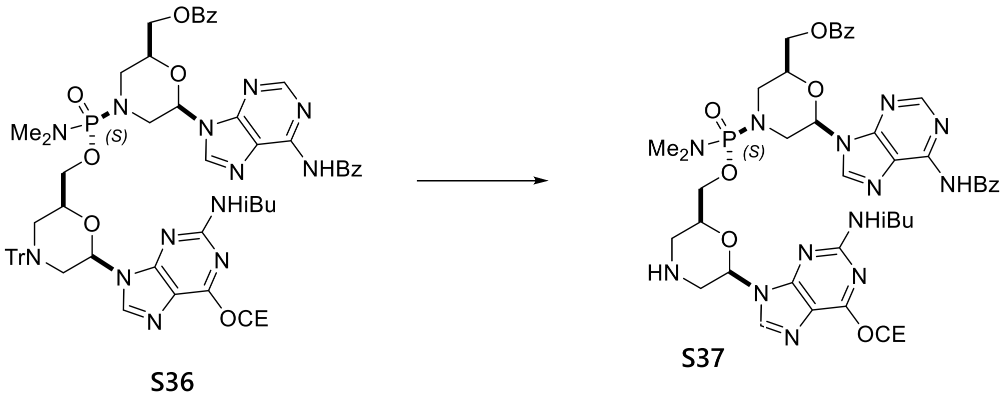

To a solution of starting material **S36** (2.316 g, 1.964 mmol) in $CH_2Cl_2$ (35 mL) at room temperature was added ethanol (1.2 mL, 20 mmol) followed by TFA (0.91 mL, 12 mmol). The reaction mixture was stirred at room temperature for 1 h, and then treated with 1,2,2,6,6-pentamethylpiperidine (2.7 mL, 15 mmol). The resulting mixture was concentrated in vacuo. The residue was treated with EtOAc (25 mL) followed by MTBE (50 mL). The resulting slurry was filtered through a glass filter and rinsed with a mixture of MTBE and EtOAc (15 mL/5 mL). The filter cake was dried in vacuo for 2 h to provide 1.75 g of target product **S37**.

MS (ESI) m/z: $[M+H]^+$ Calcd for $C_{43}H_{50}N_{14}O_9P$ 937.36; Found 937.10.

3-mer of 3'-PMO: coupling

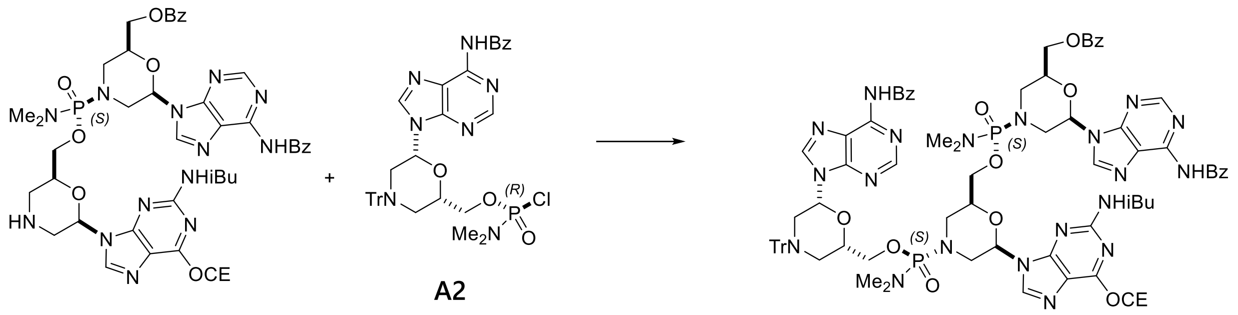

S37 S38

To a solution of starting material **S37** (1.75 g, 1.87 mmol) in 1,3-dimethyl-2-imidazolidinone (20 mL) at 0 °C was added 1,2,2,6,6-pentamethylpiperidine (0.68 mL, 3.7 mmol) followed by reactant **A2** (1.42 g, 1.96 mmol). The reaction mixture was warmed to room temperature and stirred overnight. To the reaction mixture was added EtOAc (20 mL) followed by MTBE (60 mL) and *n*-heptane (80 mL). The precipitate was collected by decantation. The isolated product (**S38**) was directly used for the next step without further purification.

MS (ESI) m/z: $[M+H]^+$ Calcd for $C_{81}H_{86}N_{21}O_{13}P_2$ 1622.62; Found 1622.59.

3-mer of 3'-PMO: deprotection

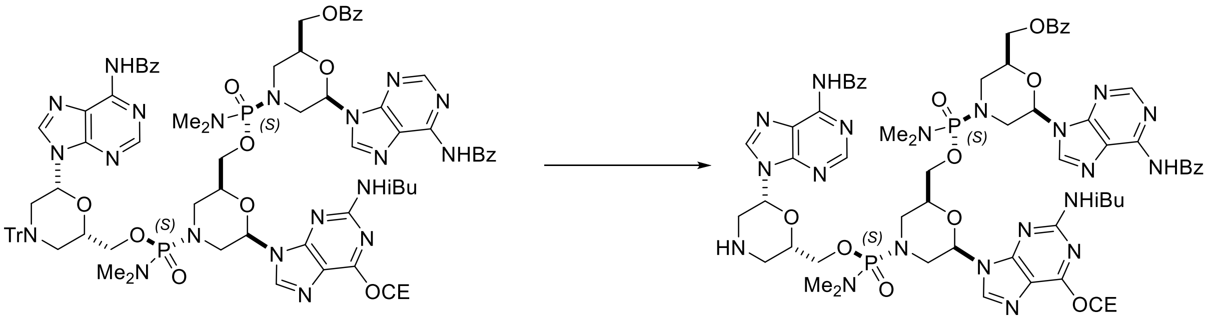

S38 S39

To a solution of starting material **S38** (3.03 g, 1.87 mmol in theory) in $CH_2Cl_2$ (24 mL) at room temperature were added ethanol (1.1 mL, 19 mmol) and TFA (0.86 mL, 11.2 mmol). The reaction mixture was stirred for 30 min before additional TFA (0.43 mL, 5.6 mmol) was added. After being stirred for 2 h, the reaction mixture was treated with EtOAc (75 mL) followed by MTBE (50 mL). The precipitate was collected by filtration and rinsed with EtOAc/MTBE (10 mL/10 mL). The resulting solid was dissolved in $CH_2Cl_2$ (25 mL) and treated with 1,2,2,6,6-pentamethylpiperidine (1.02 mL, 5.60 mmol) at room temperature. The mixture was stirred for 10 min before EtOAc (75 mL) and MTBE (50 mL) were added. The resulting precipitate was collected by filtration and rinsed with EtOAc/MTBE (15 mL/15 mL). Drying the filter cake in vacuo provided 2.25 g of target product **S39**.

MS (ESI) m/z: $[M+H]^+$ Calcd for $C_{62}H_{72}N_{21}O_{13}P_2$ 1380.51; Found 1380.31.

4-mer of 3’-PMO: coupling

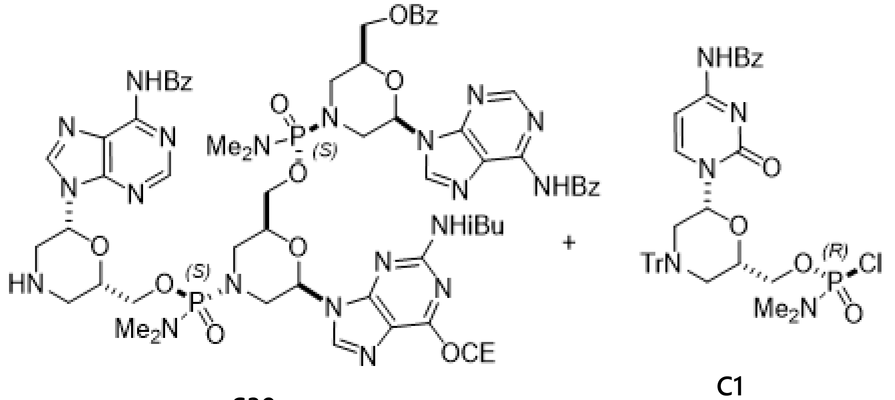

S39 C1

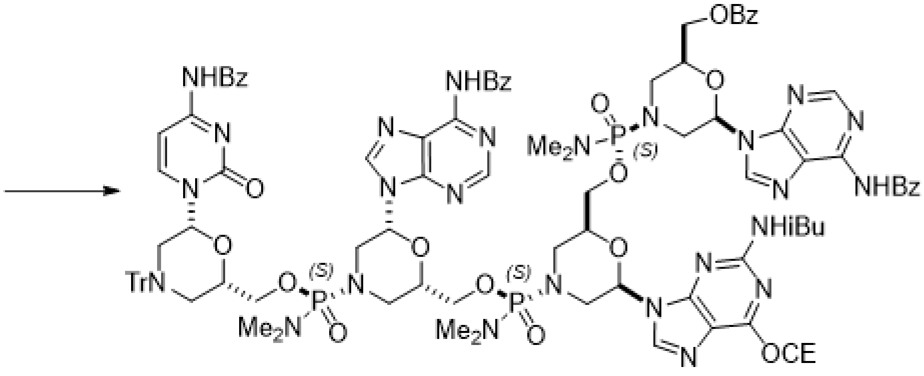

S40

To a solution of starting material **S39** (2.20 g, 1.59 mmol) in 1,3-dimethyl-2-imidazolidinone (20 mL) at room temperature was added 1,2,2,6,6-pentamethylpiperidine (0.73 mL, 4.0 mmol) followed by reactant **C1** (1.22 g, 1.75 mmol). The reaction mixture was stirred overnight before additional **C1** (0.20 g, 0.29 mmol) was added. After being stirred for additional 4 h, the reaction mixture was treated with morpholine (42 μL, 0.48 mmol). After 20 min, EtOAc (20 mL) and MTBE (150 mL) were added. The resulting precipitate was collected by filtration, rinsed with a mixture of EtOAc/MTBE (10 mL/20 mL) and dried in vacuo overnight. The resulting solid (3.74 g) was dissolved in $CH_2Cl_2$ (25 mL). To the solution was added EtOAc (25 mL) followed by MTBE (100 mL). The resulting precipitate was collected by filtration, rinsed with a mixture of EtOAc/MTBE (10 mL/30 mL), and dried in vacuo overnight. 3.20 g of target product **S40** was obtained.

MS (ESI) m/z: $[M\text{-}Tr+2H]^+$ Calcd for $C_{80}H_{94}N_{26}O_{18}P_3$ 1800.65; Found 1800.05.

4-mer of 3'-PMO: deprotection

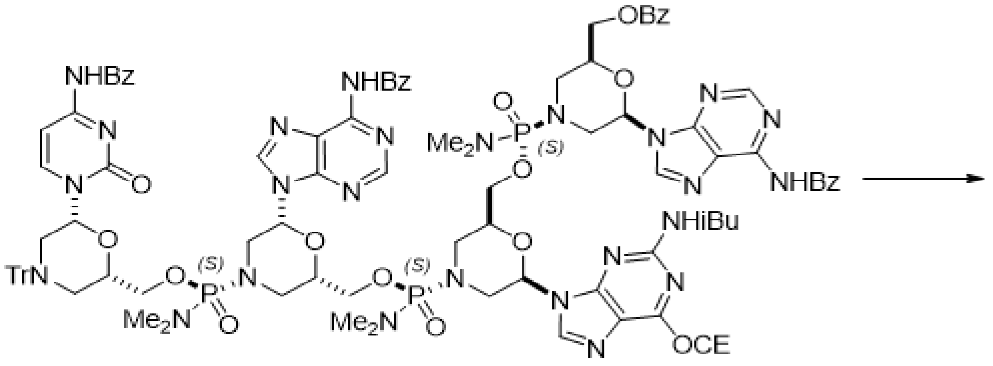

**S40**

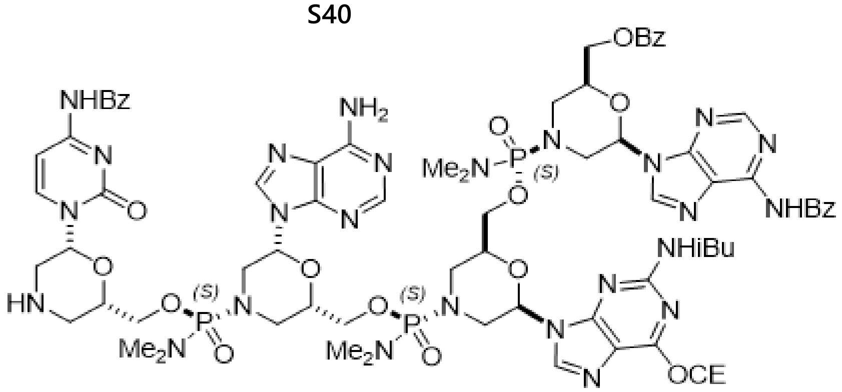

**S41**

To a solution of starting material **S40** (194 mg, 1.57 mmol) in $CH_2Cl_2$ (42 mL) at room temperature were added EtOH (0.92 mL) and TFA (0.96 mL, 12 mmol). The reaction mixture was stirred for 2 h and treated with EtOAc (4 mL) followed by MTBE (80 mL). The resulting precipitate was collected by filtration and rinsed with a mixture of EtOAc/MTBE (10 mL/20 mL). The resulting solid was dissolved in $CH_2Cl_2$ (42 mL) and treated with 1,2,2,6,6-pentamethylpiperidine (0.85 mL, 4.7 mmol). The resulting solution was stirred at room temperature for 10 min before EtOAc (40 ml) and MTBE (100 mL) were added. The precipitate was collected by filtration, rinsed with a mixture of EtOAc/MTBE (20 mL/40 mL), and dried in vacuo for 2 h. The solid was dissolved in $CH_2Cl_2$ (40 mL). To the solution was added EtOAc (40 mL) followed by MTBE (60 mL). The resulting precipitate was collected by filtration and rinsed with a mixture of EtOAc/MTBE (20 mL/20 mL). The solid was dissolved in $CH_2Cl_2$ (40 mL) and treated with EtOAc (80 mL). The resulting precipitate was collected by filtration and rinsed with EtOAc (~30 mL). Drying the filter cake in vacuo provided 2.05 g of target product **S41**.

MS (ESI) m/z: $[M+H]^+$ Calcd for $C_{80}H_{94}N_{26}O_{18}P_3$ 1800.65; Found 1800.68.

4-mer of 3’-PMO: global deprotection

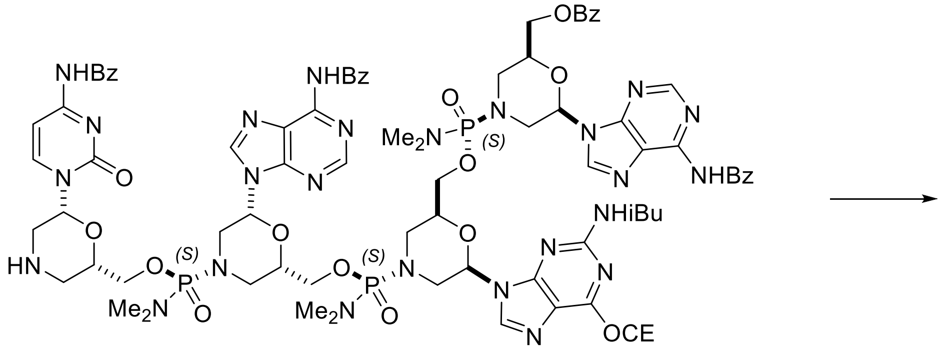

**S41**

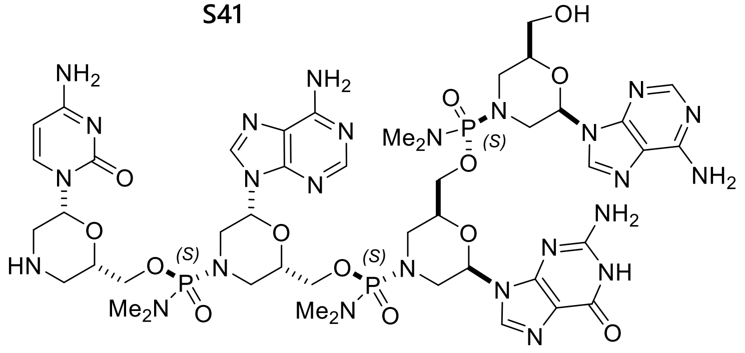

**S42**

Starting material **S41** (1.25 g, 0.695 mmol) was dissolved in a mixture of methanol (20 mL) and 28% ammonium hydroxide (20 mL) at room temperature. To the solution was added morpholine (0.73 mL, 8.3 mmol). The resulting mixture was heated at 50-52 °C for 15 h and cooled to room temperature. After concentration in vacuo, the residue was dissolved in $CH_2Cl_2$/MeOH (12.5 mL/5 mL) and treated with EtOAc (60 mL). The resulting precipitate was collected by filtration and rinsed with a mixture of EtOAc/$CH_2Cl_2$/MeOH (20 mL/2.5 mL/1 mL). Drying the filter cake in vacuo overnight afforded 928 mg of target product **S42**.

MS (ESI) m/z: $[M+H]^+$ Calcd for $C_{45}H_{69}N_{25}O_{13}P_3$ 1260.47; Found 1260.98.

4-mer of 3'-PMO: morpholine protection

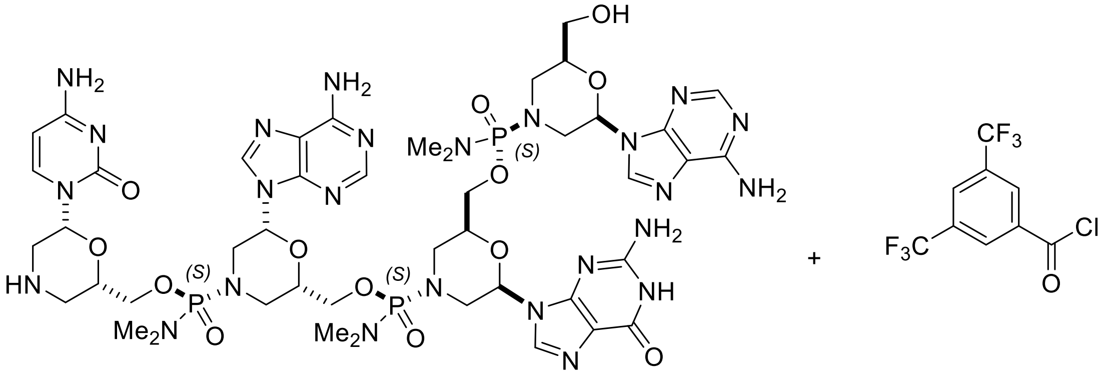

S42

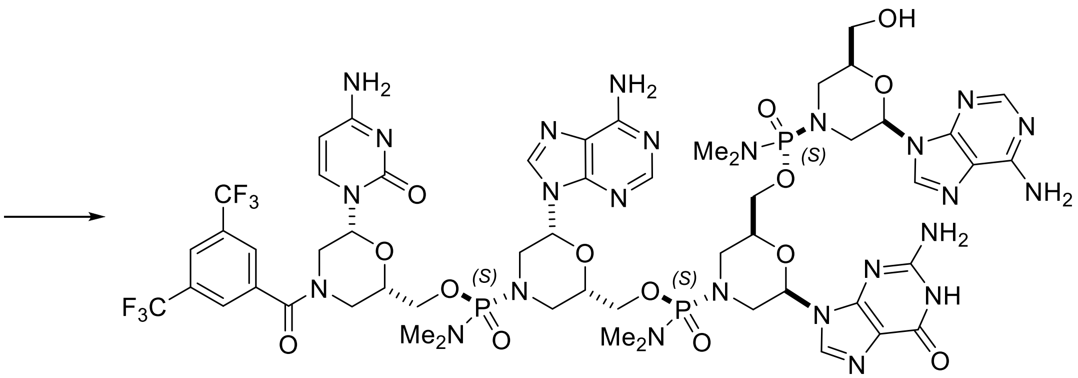

S43

To a solution of starting material **S42** (928 mg, 0.405 mmol in theory) in a mixture of THF/Water/MeOH (15 mL/2.5 mL/4.5 mL) were added 1,2,2,6,6-pentamethylpiperidine (0.367 mL, 2.02 mmol) and 3,5-bis(trifluoromethyl)benzoyl chloride (0.11 mL, 0.61 mmol). After 3 h, additional 0.025 mL of bis(trifluoromethyl)benzoyl chloride was added. After being stirred overnight, the reaction mixture was treated with EtOAc (60 mL). The resulting gummy solid was isolated by decantation and dissolved in a mixture of MeOH/ $CH_2Cl_2$ (2 mL/8 mL). To the solution was added EtOAc (50 mL). The resulting precipitate was isolated by filtration, rinsed with EtOAc, and dried in vacuo for 20 min. The resulting solid was treated with a mixture of MeCN/EtOAc (7.5 mL/7.5 mL). The slurry was filtered through a glass filter and rinsed with a mixture of MeCN/EtOAc (2.5 mL/2.5 mL). Drying the filter cake in vacuo for 1 h afforded 550 mg of target product **S43**.

$^{31}$P NMR (162 MHz, METHANOL-d4) δ = 17.16 (s, 1P), 17.11 (s, 1P), 16.97 (s, 1P)

MS (ESI) m/z: $[M+H]^+$ Calcd for $C_{54}H_{71}F_6N_{25}O_{14}P_3$ 1500.47; Found.1500.22.

**Elongation of DNA gap**

5-mer: coupling

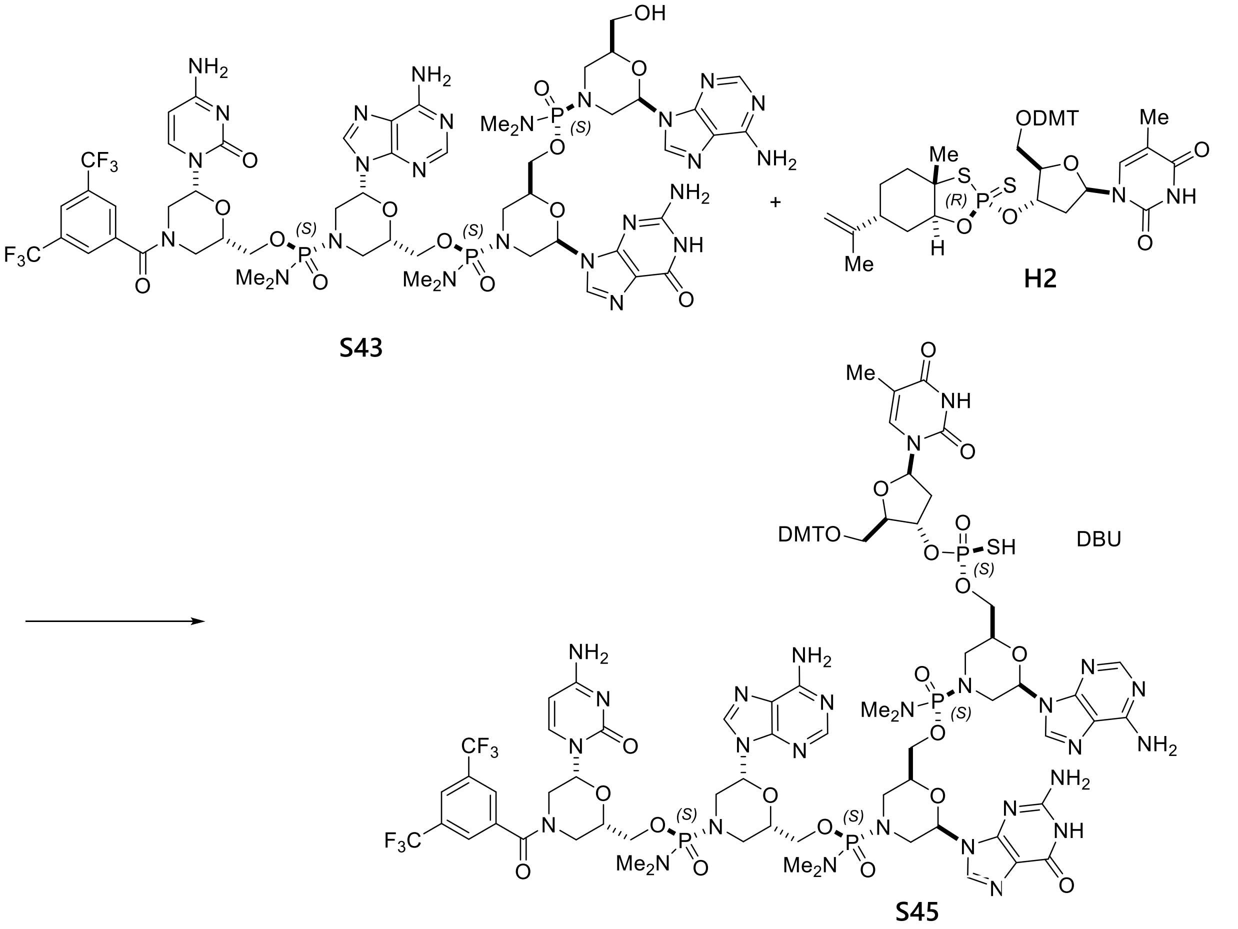

Starting material **S43** (550 mg, 0.367 mmol) and reactant **H2** (783 mg, 0.99 mmol) were dissolved in 1,3-dimethyl-2-imidazolidinone (19 mL). To the resulting solution was added 4Å molecular sieves (1.7 g). The reaction flask was applied to vacuum and filled with nitrogen. The process was repeated two more times. After being stirred for 30 min, the resulting mixture was treated with DBU (0.22 mL, 1.47 mmol). The reaction mixture was stirred for 1 hr at room temperature and then filtered through a syringe filter. The filtrate was added into EtOAc (30 mL), rinsing with 1,3-dimethyl-2-imidazolidinone (6 mL). To the resulting slurry was added additional EtOAc (50 mL). The resulting precipitate was collected by filtration and rinsed with a mixture of EtOAc/MeCN (10 mL/10 mL). The filter cake was treated with MeCN (20 mL) followed by EtOAc (20 mL). After 10 min, the resulting slurry was filtered through a glass filter and rinsed with EtOAc/MeCN (5 mL/5 mL). Drying the filter cake in vacuo for 3 days afforded 790 mg of target product **S45**.

$^{31}$P NMR (162 MHz, METHANOL-d4) δ = 57.76 (s, 1P), 17.10 (s, 1P), 17.02 (s, 1P), 16.90 (s, 1P).

MS (ESI) m/z: [M-DMT+2H]$^+$ Calcd for $C_{64}H_{84}F_6N_{27}O_{20}P_4S$ 1820.50; Found 1820.18.

5-mer: deprotection

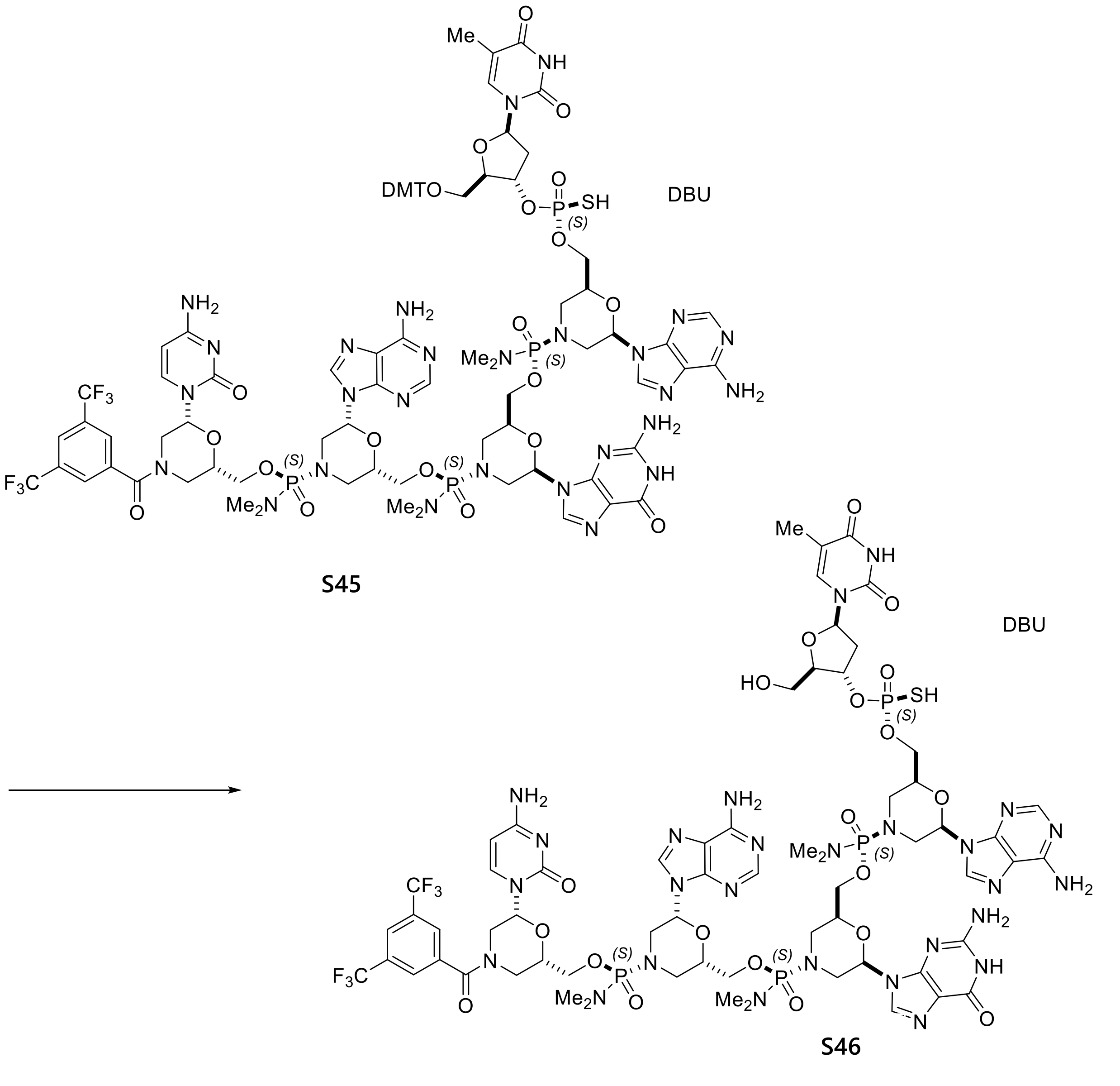

Starting material **S45** (0.790 g, 0.347 mmol) was dissolved in a mixture of 1,1,1,3,3,3-hexafluoro-2-propanol (8 mL), 2,2,2-trifluoroethanol (2 mL), $CH_2Cl_2$ (10 mL) and triethylsilane

(6 mL). The reaction mixture was stirred for 3 h at room temperature, and an additional mixture of 1,1,1,3,3,3-hexafluoro-2-propanol (2 mL), 2,2,2-trifluoroethanol (0.5 mL), $CH_2Cl_2$ (2.5 mL) and triethylsilane (1.5 mL) was added. After additional 1 h stirring, the reaction mixture was treated with EtOAc (150 mL) followed by MTBE (75 mL). The resulting precipitate was collected by centrifuge (3500 rpm, 35 min) and rinsed with a mixture of EtOAc/MeCN (10 mL/10 mL). The pellet was treated with MeCN (25 mL) to make a slurry. After 5 min stirring, EtOAc (25 mL) was added. The resulting slurry was filtered through a glass filter and rinsed with MeCN/EtOAc (10 mL/10 mL). Drying the filter cake in vacuo overnight provided 646 mg of target product **S46**.
MS (ESI) m/z: $[M-H]^-$ Calcd for $C_{64}H_{82}F_6N_{27}O_{20}P_4S$ 1818.48; Found 1818.37.

6-mer: coupling

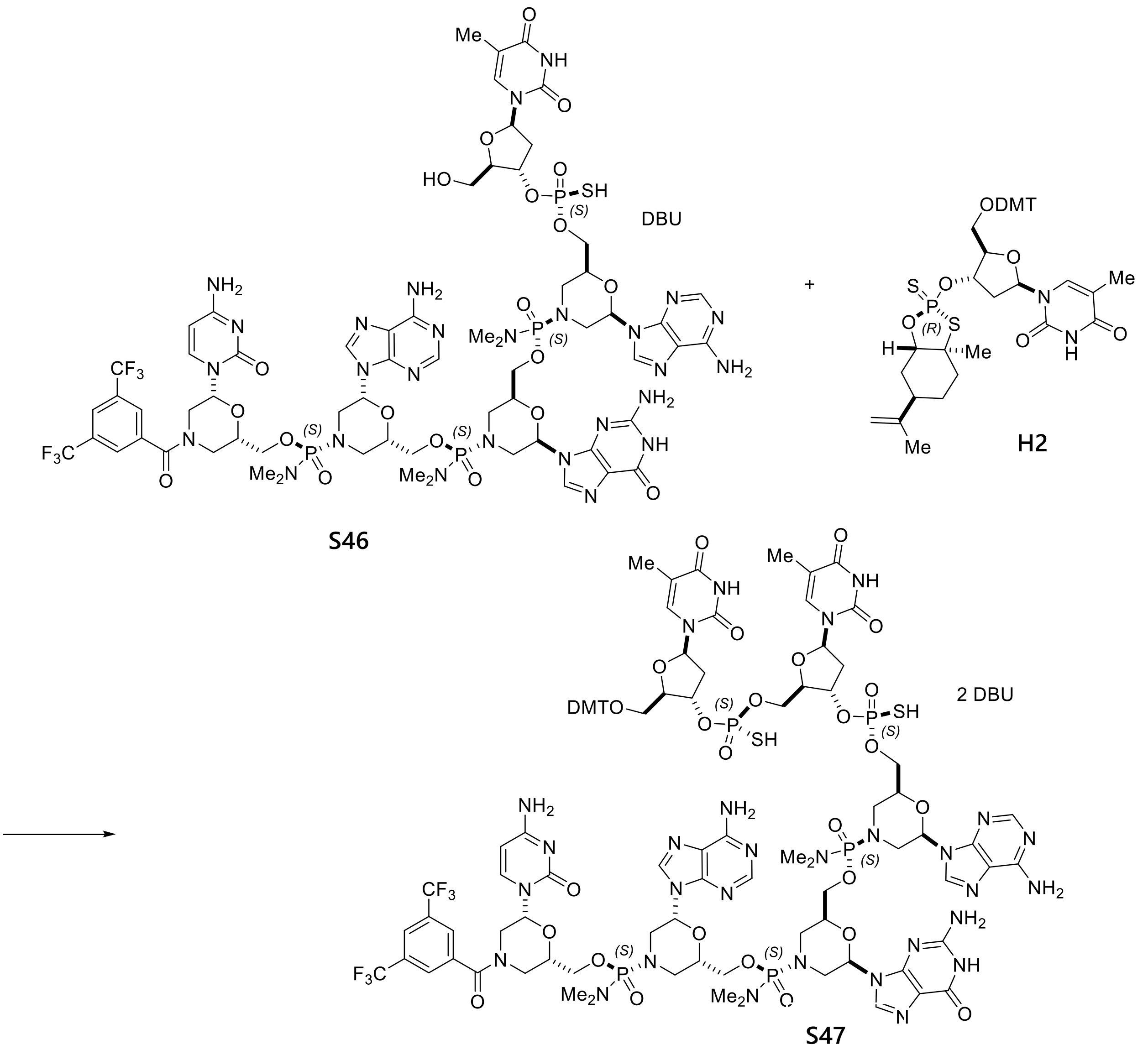

Starting material **S46** (646 mg, 0.327 mmol) and reactant **H2** (777 mg, 0.982 mmol) were dissolved in 1,3-dimethyl-2-imidazolidinone (16 mL). To the resulting solution was added 4 Å molecular sieves (2 g). The reaction flask was applied to vacuum and filled with nitrogen. The process was repeated two more times. After being stirred for 30 min, the resulting mixture was treated with DBU (0.25 mL, 1.64 mmol). The reaction mixture was stirred for 2 h at room temperature and then filtered through a syringe filter. The filtrate was added into EtOAc (35 mL), rinsing with 1,3-dimethyl-2-imidazolidinone (4 mL). To the resulting slurry was added additional EtOAc (40 mL). The precipitate was isolated by filtration and rinsed with MeCN/EtOAc (5 mL/ 5

mL). The resulting solid was treated with MeCN (20 mL) followed by EtOAc (20 mL). The resulting slurry was filtered through a glass filter and rinsed with EtOAc/MeCN (5 mL/5 mL). Drying the filter cake in vacuo overnight provided 0.90 g of target product **S47**.
MS (ESI) m/z: $[M-2H]^{2-}$ Calcd for $C_{95}H_{113}F_6N_{29}O_{28}P_5S_2$ 1220.32; Found 1220.47.

6-mer: deprotection

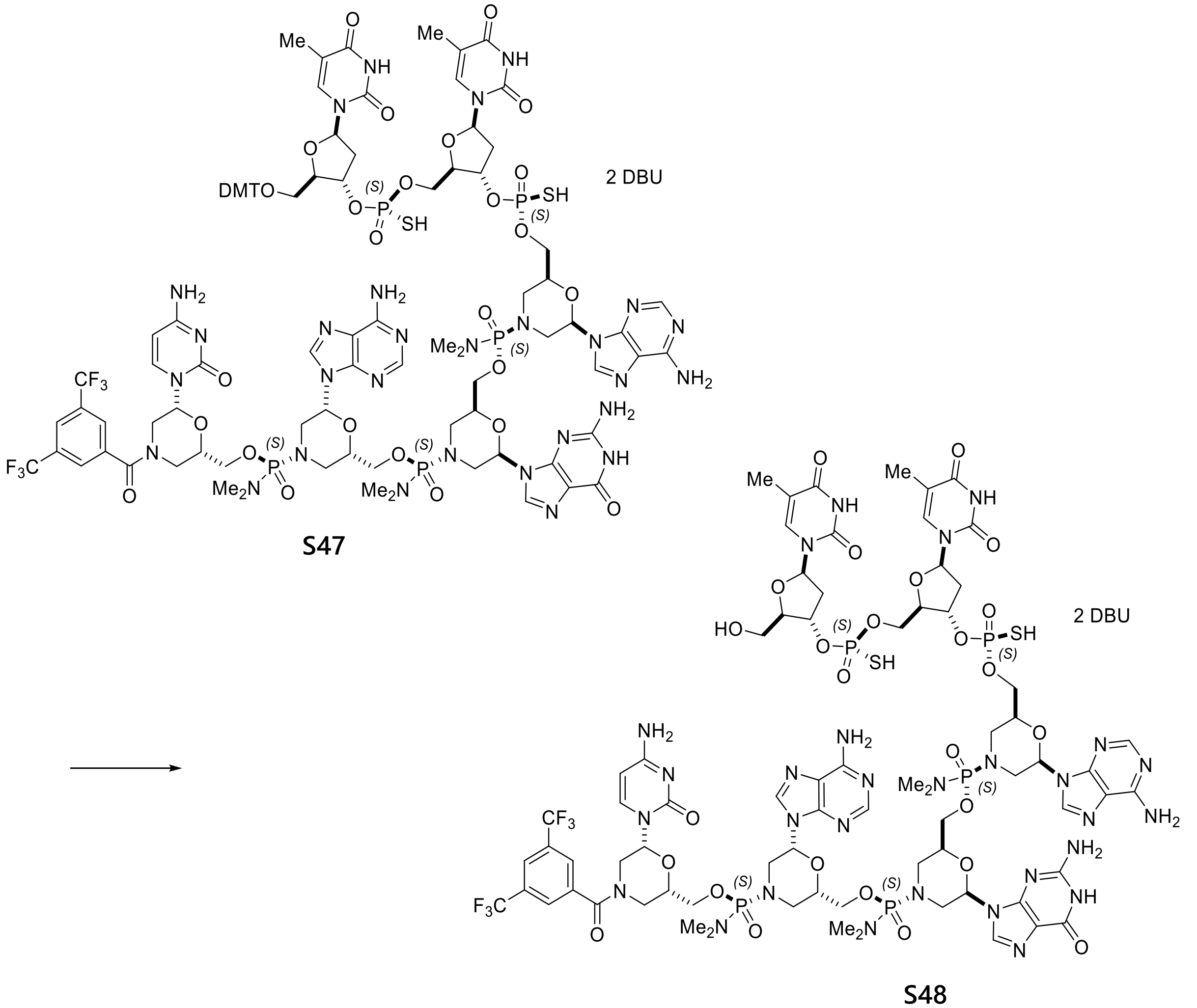

To starting material **S47** (0.90 g, 0.328 mmol) was added a mixture of 1,1,1,3,3,3-hexafluoro-2-propanol (10.8 mL), 2,2,2-trifluoroethanol (2.7 mL), triethylsilane (8.1 mL) and $CH_2Cl_2$ (13.5 mL). After being stirred at room temperature overnight, the reaction mixture was treated with EtOAc (150 mL) followed by MTBE (100 mL). The resulting precipitate was isolated by filtration and rinsed with a mixture of EtOAc/MeCN (10 mL/10 mL). The filter cake was treated with MeCN (25 mL) to make a slurry. After 5 min stirring, EtOAc (25 mL) was added. The resulting slurry was filtered through a glass filter and rinsed with MeCN/EtOAc (10 mL/10 mL). Drying the filter cake in vacuo for 1 h provided 800 mg of target product **S48**.

MS (ESI) m/z: $[M+2H]^{2+}$ Calcd for $C_{74}H_{98}F_6N_{29}O_{26}P_5S_2$ 1070.76; Found 1070.66.

<u>7-mer: coupling</u>

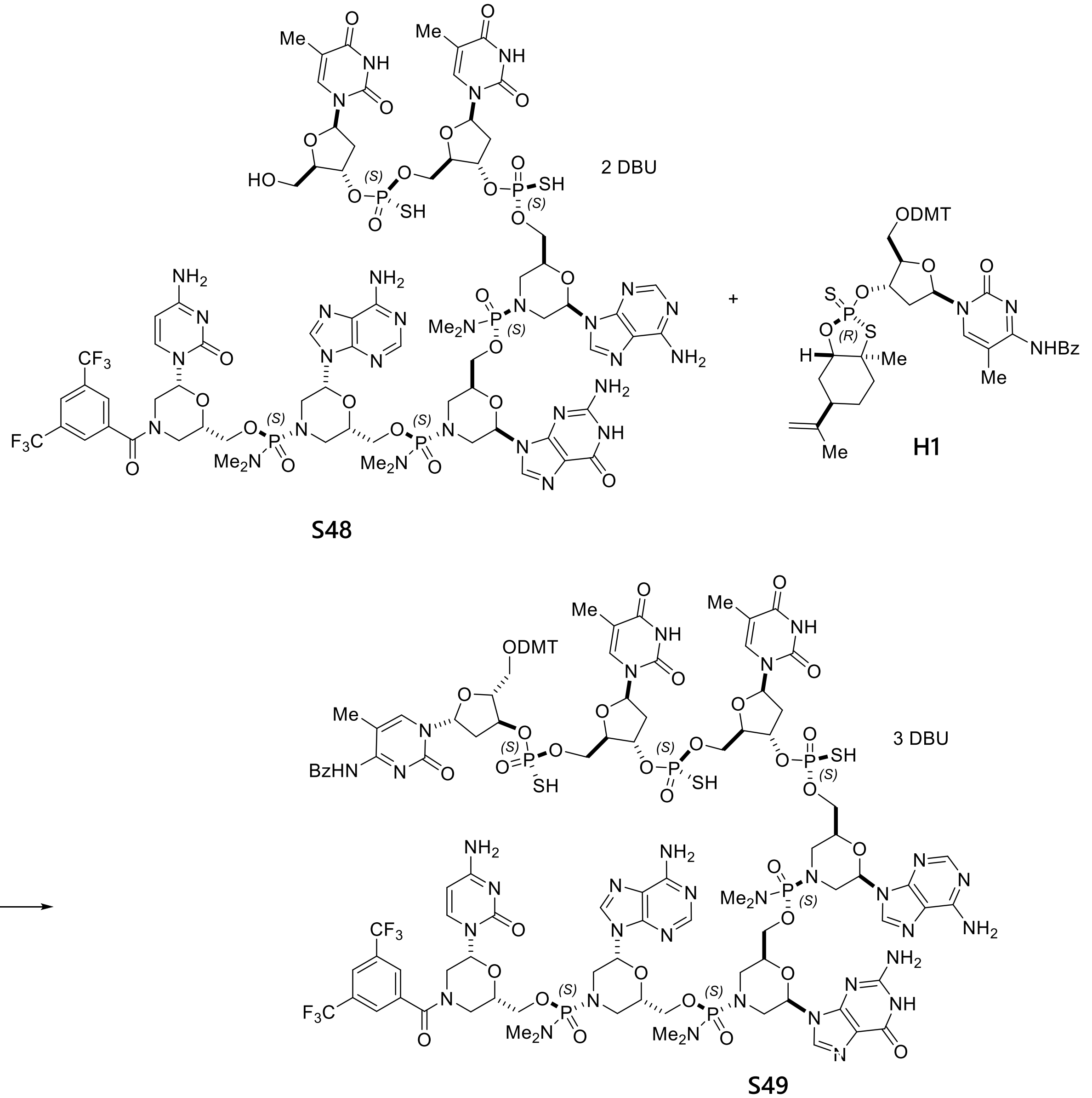

**S49**

Starting material **S48** (950 mg, 0.389 mmol) and reactant **H1** (1042 mg, 1.17 mmol) were dissolved in 1,3-dimethyl-2-imidazolidinone (23.8 mL). To the resulting solution was added 4 Å molecular sieves (1 g). The reaction flask was applied to vacuum and filled with nitrogen. The process was repeated two more times. After being stirred for 30 min, the resulting mixture was treated with DBU (0.35 mL, 2.33 mmol). The reaction mixture was stirred for 16 h at room temperature and then filtered through a syringe filter. The filtrate was added into EtOAc (40 mL), rinsing with 1,3-dimethyl-2-imidazolidinone (5 mL). To the resulting slurry was added additional EtOAc (35 mL). The precipitate was isolated by filtration and rinsed with MeCN/EtOAc (10 mL/10 mL). The resulting solid was treated with MeCN (20 mL) followed by EtOAc (20 mL). The resulting slurry was filtered through a glass filter and rinsed with EtOAc/MeCN (7.5 mL/7.5 mL). Drying the filter cake in vacuo for 4 h provided 1.20 g of target product **S49**.

$^{31}$P NMR (162 MHz, METHANOL-d4) δ = 57.13 (s, 1P), 56.94 (s, 2P), 17.05 (s, 1P), 16.98 (s, 1P), 16.79 (s, 1P).

MS (ESI) m/z: $[M-2H]^{2-}$ Calcd for $C_{112}H_{130}F_6N_{32}O_{34}P_6S_3$ 1431.35; Found 1431.26.

7-mer: deprotection

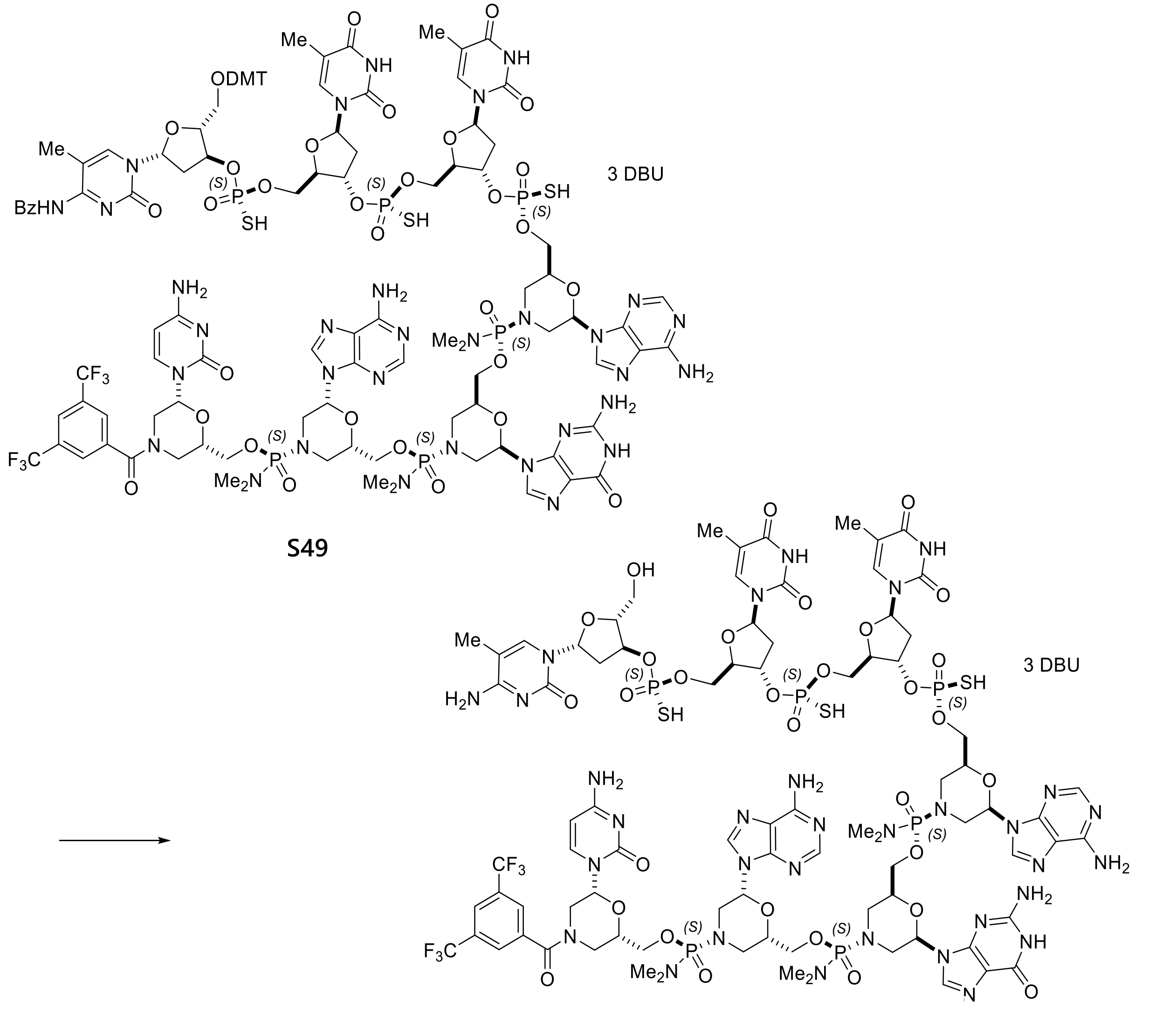

**S49** → **S50**

To starting material **S49** (1.20 g, 0.361 mmol) was added a mixture of 1,1,1,3,3,3-hexafluoro-2-propanol (14.4 mL), 2,2,2-trifluoroethanol (3.6 mL), triethylsilane (10.8 mL) and $CH_2Cl_2$ (18 mL). After being stirred at room temperature overnight, the resulting solution was treated with EtOAc (100 mL) followed by MTBE (50 mL). The resulting precipitate was collected by filtration and rinsed with a mixture of EtOAc/MeCN (10 mL/10 mL). The filter cake was treated with MeCN (25 mL) followed by EtOAc (25 mL). The resulting slurry was filtered through a glass filter and rinsed with MeCN/EtOAc (10 mL/10 mL). Drying the filter cake in vacuo for 3 h provided 1.0 g of target product **S50**.

MS (ESI) m/z: $[M-2H]^{2-}$ Calcd for $C_{84}H_{108}F_6N_{32}O_{31}P_6S_3$ 1228.27; Found 1228.50.

8-mer: coupling

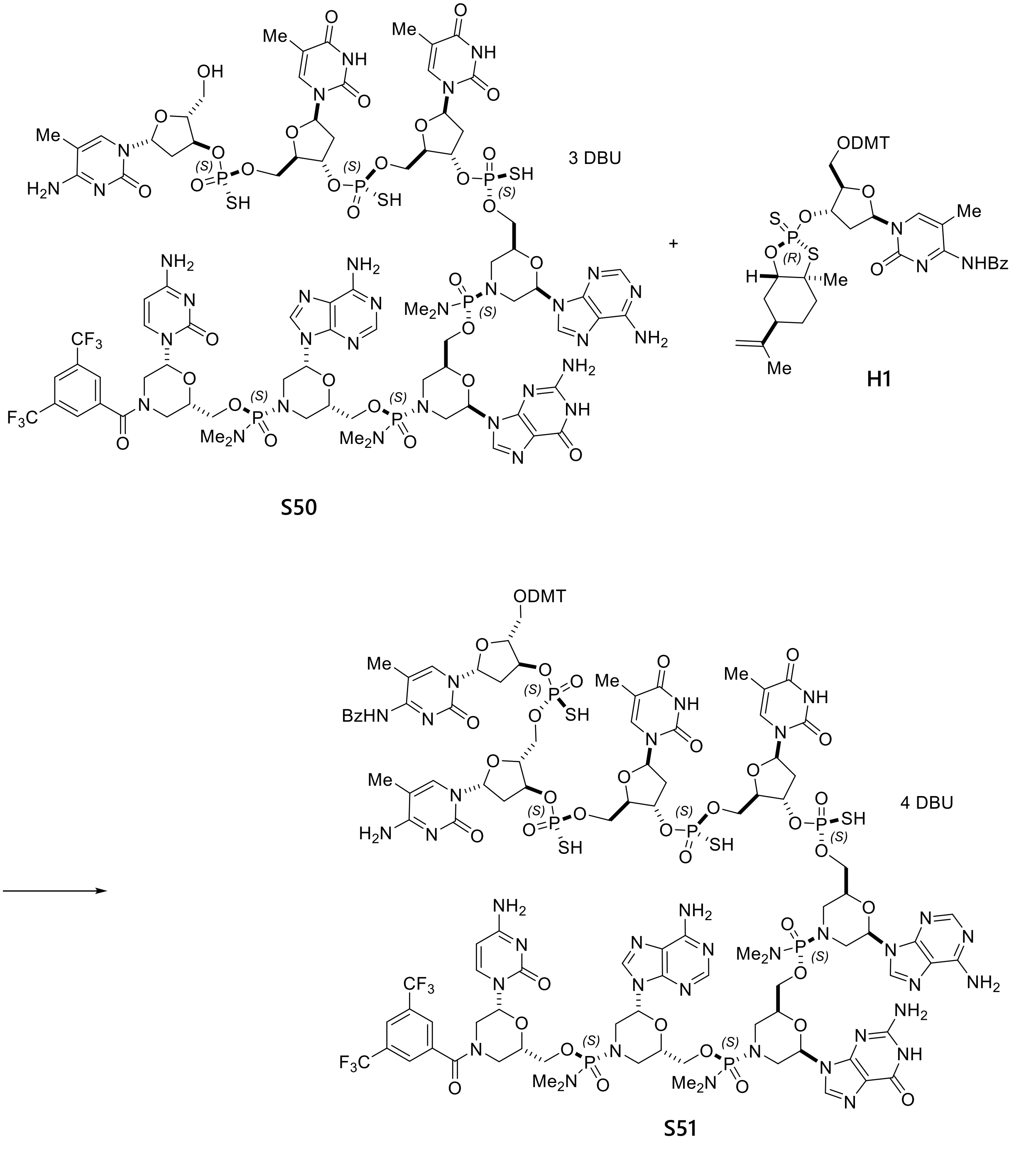

To a solution of starting material **S50** (300 mg, 0.103 mmol) in 1,3-dimethyl-2-imidazolidinone (9.0 mL) was added reactant **H1** (276 mg, 0.309 mmol). To the resulting solution was added 4 Å molecular sieves (1.0 g). The reaction flask was applied to vacuum and filled with nitrogen and the process was repeated two more times. After being stirred for 30 min, the resulting mixture was treated with DBU (0.11 mL, 0.72 mmol). The reaction mixture was stirred for 4 h at room temperature and then filtered through a syringe filter. The filtrate was added into EtOAc (25 mL), rinsing with 1,3-dimethyl-2-imidazolidinone (4.5 mL). To the resulting slurry was added additional EtOAc (20 mL). The precipitate was isolated by filtration and rinsed with MeCN/EtOAc (7.5 mL/7.5 mL). The resulting solid was treated with MeCN (10 mL) followed by EtOAc (10 mL). The resulting slurry was filtered through a glass filter and rinsed with EtOAc/MeCN (5 mL/5 mL). Drying the filter cake in vacuo overnight provided 0.36 g of target product **S51**.

$^{31}$P NMR (162 MHz, METHANOL-d4) δ = 57.36 (s, 1P), 57.31 (s, 1P), 56.90 (s, 1P), 56.27 (s, 1P) 16.96 (s, 1P), 16.94 (s, 1P), 16.67 (s, 1P).

MS (ESI) m/z: $[M-2H]^{2-}$ Calcd for $C_{122}H_{144}F_6N_{35}O_{39}P_7S_4$ 1591.37; Found 1591.35.

8-mer: deprotection

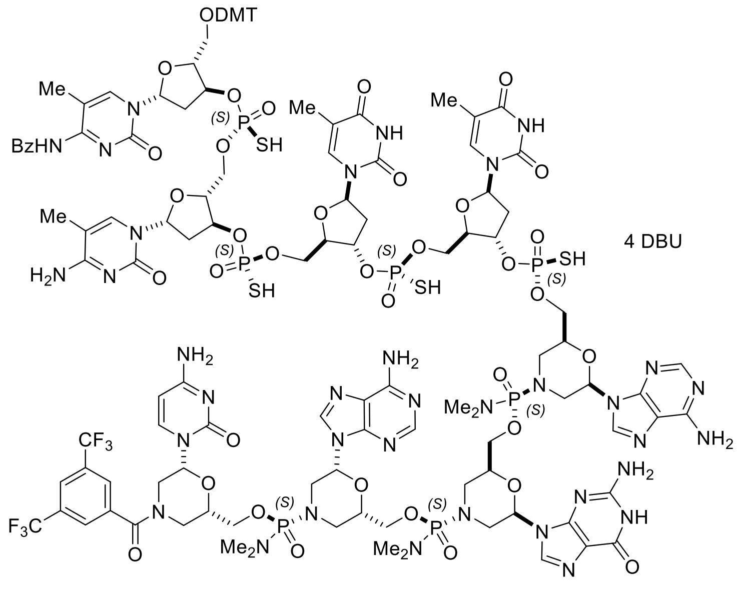

**S51**

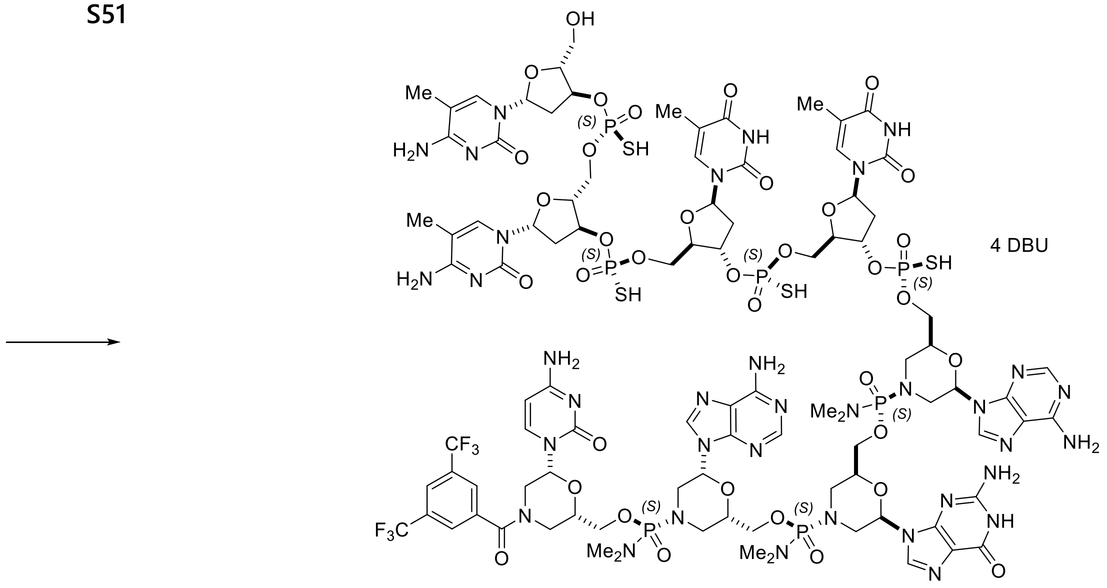

**S52**

To starting material **S51** (360 mg, 0.095 mmol) was added a mixture of 1,1,1,3,3,3-hexafluoro-2-propanol (4.3 mL), 2,2,2-trifluoroethanol (1.1 mL), triethylsilane (3.2 mL) and $CH_2Cl_2$ (5.4 mL). The resulting solution was stirred at room temperature for 17 h and treated with EtOAc (75 mL) followed by MTBE (15 mL). The resulting precipitate was collected by filtration and rinsed with a mixture of EtOAc/MeCN (5 mL/5 mL). The filter cake was treated with MeCN (15 mL) followed by EtOAc (15 mL). The resulting slurry was filtered through a glass filter and rinsed with MeCN/EtOAc (5 mL/5 mL). Drying the filter cake in vacuo for 2 h provided 0.305 g of target product **S52**.

MS (ESI) m/z: $[M-2H]^{2-}$ Calcd for $C_{94}H_{122}F_6N_{35}O_{36}P_7S_4$ 1388.29; Found 1388.26.

9-mer: coupling

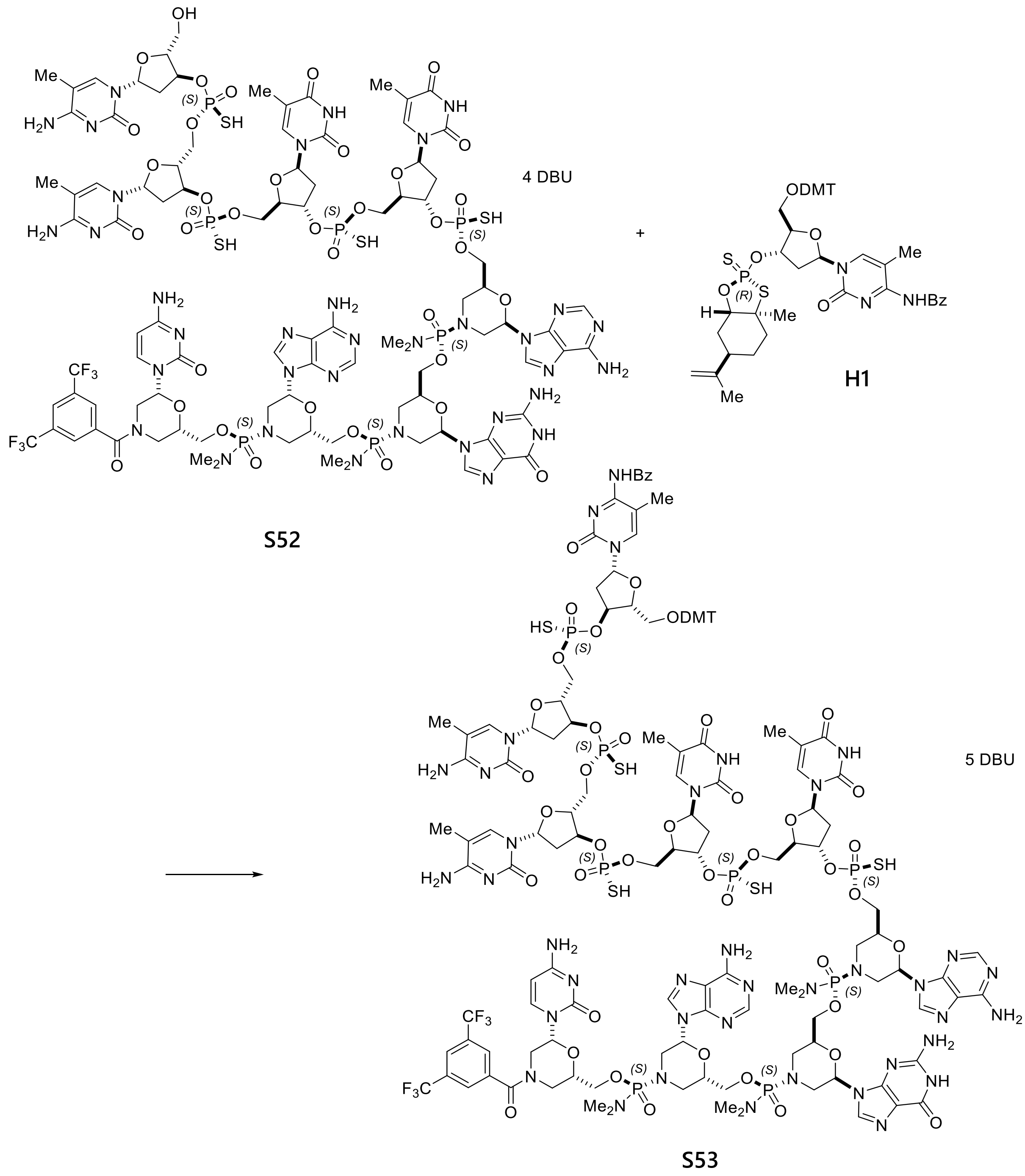

To a solution of starting material **S52** (305 mg, 0.090 mmol) in 1,3-dimethyl-2-

imidazolidinone (12 mL) was added reactant **H1** (241 mg, 0.270 mmol). To the resulting solution was added 4 Å molecular sieves (1 g). The reaction flask was applied to vacuum and filled with nitrogen. The process was repeated two more times. After being stirred for 30 min, the resulting mixture was treated with DBU (0.11 mL, 0.72 mmol). The reaction mixture was stirred for 2.5 days at room temperature and then filtered through a syringe filter. The filtrate was added into EtOAc (20 mL), rinsing with 1,3-dimethyl-2-imidazolidinone (4 mL). To the resulting slurry was added additional EtOAc (20 mL). The resulting precipitate was collected by centrifuge (3500 rpm, 30 min). The resulting pellet was rinsed with a mixture of MeCN/EtOAc (5 mL/ 5 mL), and treated with MeCN (15 mL) followed by EtOAc (15 mL). The resulting slurry was subjected to centrifuge (3500 rpm, 10 min). The pellet was rinsed with a mixture of MeCN/EtOAc (5 mL/5 mL), and dried in vacuo for 1h. 385 mg of target product **S53** was obtained.

$^{31}$P NMR (162 MHz, METHANOL-d4) δ = 57.44 (s, 1P), 57.35 (s, 1P), 56.88 (s, 2P), 56.17 (s, 1P) 16.95 (s, 1P), 16.92 (s, 1P), 16.74 (s, 1P).

MS (ESI) m/z: $[M-2H]^{2-}$ Calcd for $C_{132}H_{158}F_6N_{38}O_{44}P_8S_5$ 1751.89; Found 1751.73.

<u>9-mer: deprotection</u>

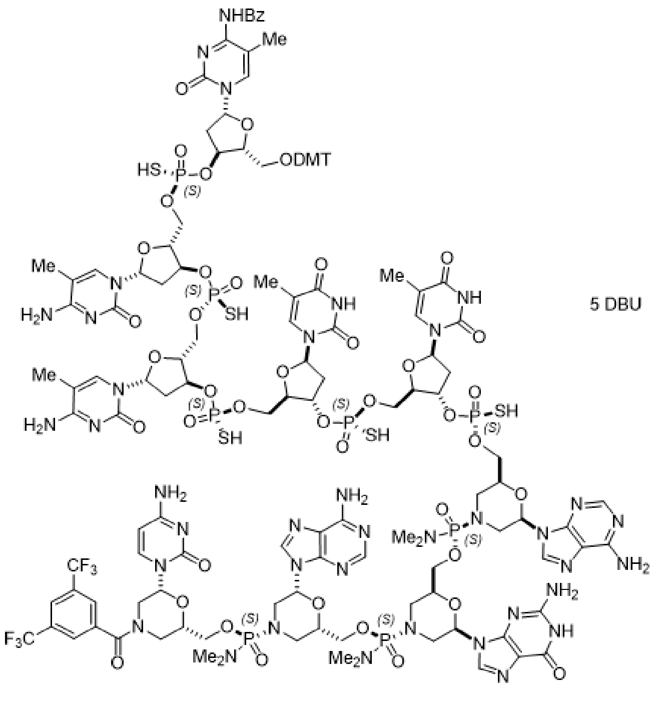

**S53**

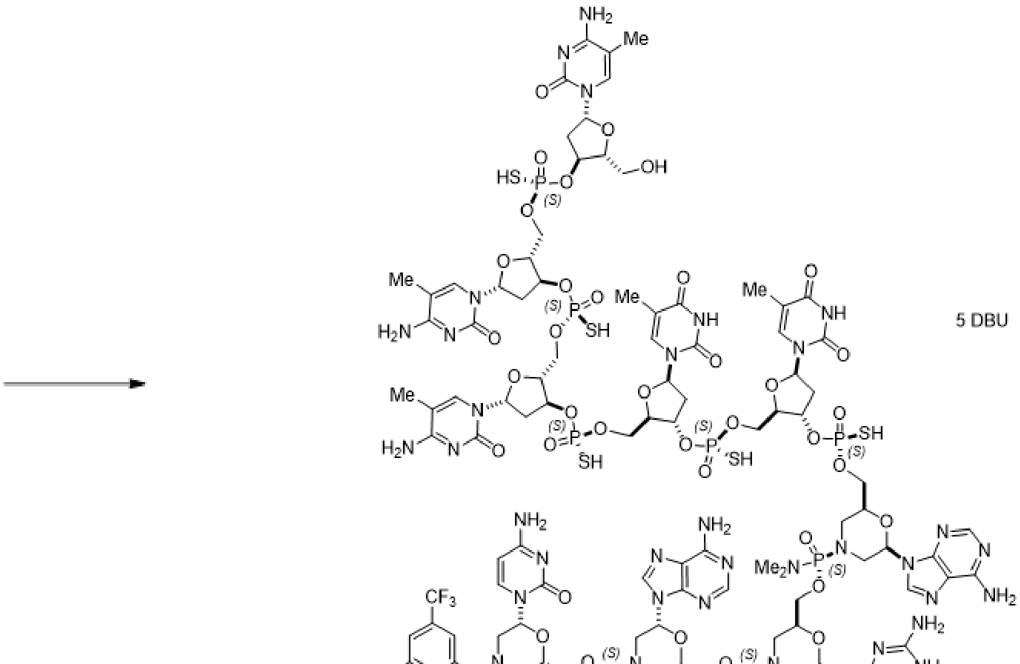

**S54**

To starting material **S53** (385 mg, 0.090 mmol) was added a mixture of 1,1,1,3,3,3-hexafluoro-2-propanol (4.6 mL), 2,2,2-trifluoroethanol (1.2 mL), triethylsilane (3.5 mL) and $CH_2Cl_2$ (5.8 mL). The resulting solution was stirred at room temperature overnight, and treat with EtOAc (90 mL). The resulting precipitate was collected by filtration and rinsed with a mixture of EtOAc/MeCN (10 mL/10 mL). Drying the filter cake in vacuo for 5 h provided 320 mg of target product **S54**.

MS (ESI) m/z: $[M-2H]^{2-}$ Calcd for $C_{104}H_{136}F_6N_{38}O_{41}P_8S_5$ 1547.81; Found 1547.81.

10-mer: coupling

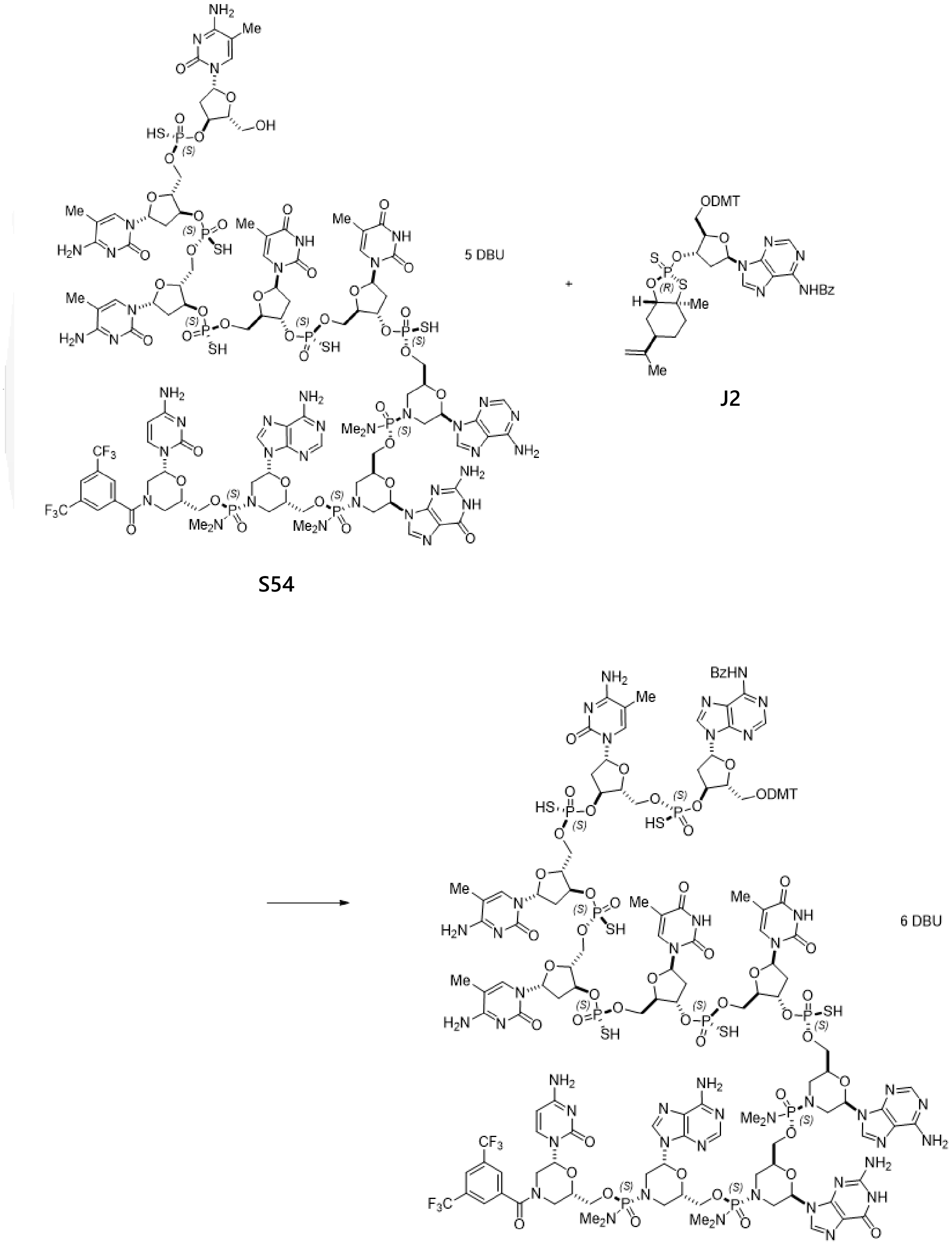

S54

J2

S55

To a solution of starting material **S54** (320 mg, 0.083 mmol) in 1,3-dimethyl-2-imidazolidinone (13 mL) was added reactant **J2** (225 mg, 0.249 mmol). To the resulting solution was added 4 Å molecular sieves (1.0 g). The reaction flask was applied to vacuum and filled with nitrogen. The process was repeated two more times. After being stirred for 30 min, the resulting mixture was treated with DBU (0.112 mL, 0.746 mmol). The reaction mixture was stirred for 17 hr at room temperature and then filtered through a syringe filter. The filtrate was added into EtOAc (20 mL), rinsing with 1,3-dimethyl-2-imidazolidinone (5 mL). To the resulting slurry was added additional EtOAc (20 mL). The resulting slurry was centrifuged (3500 rpm, 30 min). To the pellet was added MeCN (20 mL) followed by EtOAc (20 mL). The resulting slurry was centrifuged (3500 rpm, 20 min). The pellet was rinsed with EtOAc/MeCN (5 mL/5 mL) and dried in vacuo for 1 h. 420 mg of target product **S55** was obtained and used in next step without further purification.

$^{31}$P NMR (162 MHz, METHANOL-d4) δ = 57.29 (s, 1P), 56.99 (s, 1P), 56.95 (s, 1P), 56.78 (s, 2P), 56.23 (s, 1P), 16.95 (s, 2P), 16.72 (s, 1P).

MS (ESI) m/z: $[M-2H]^{2-}$ Calcd for $C_{142}H_{170}F_6N_{43}O_{48}P_9S_6$ 1915.41; Found 1915.21.

10-mer: deprotection

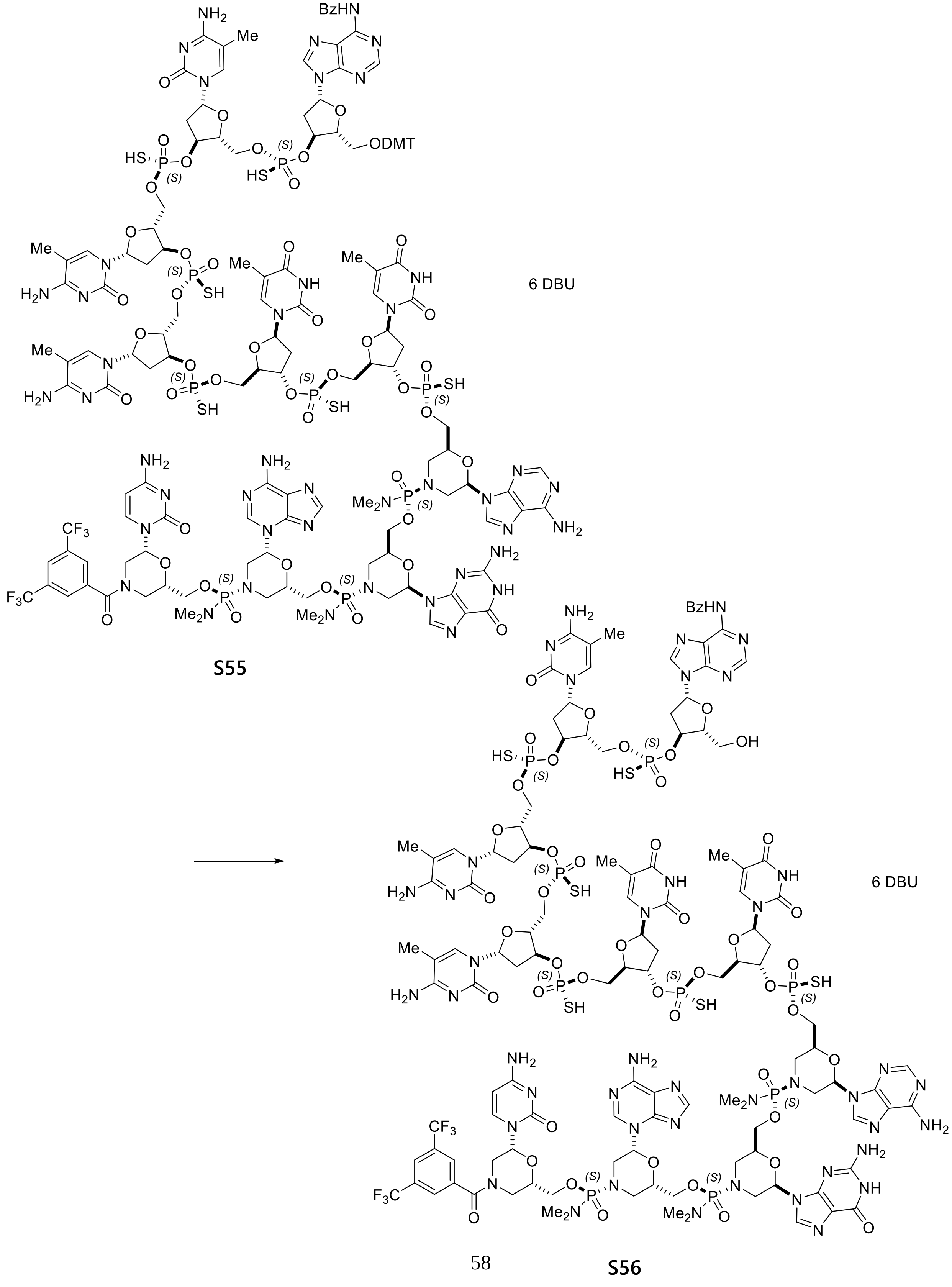

**S56**

To starting material **S55** (430 mg, 0.84 mmol in theory) was added a mixture of 1,1,1,3,3,3-hexafluoro-2-propanol (4.8 mL), 2,2,2-trifluoroethanol (1.2 mL), triethylsilane (3.6 mL) and $CH_2Cl_2$ (6.0 mL). The resulting solution was stirred at room temperature for 30 min and treated with EtOAc (90 mL). The resulting precipitate was collected by filtration and rinsed with a mixture of EtOAc/MeCN (10 mL/10 mL). The filter cake was treated with MeCN (20 mL) followed by EtOAc (20 mL). The resulting slurry was filtered through a glass filter and rinsed with a mixture of EtOAc/MeCN (10 mL/10 mL). Drying the filter cake in vacuo overnight provided 316 mg of target product **S56**.

MS (ESI) m/z: $[M-2H]^{2-}$ Calcd for $C_{121}H_{152}F_6N_{43}O_{46}P_9S_6$ 1764.34; Found 1764.19.

11-mer: coupling

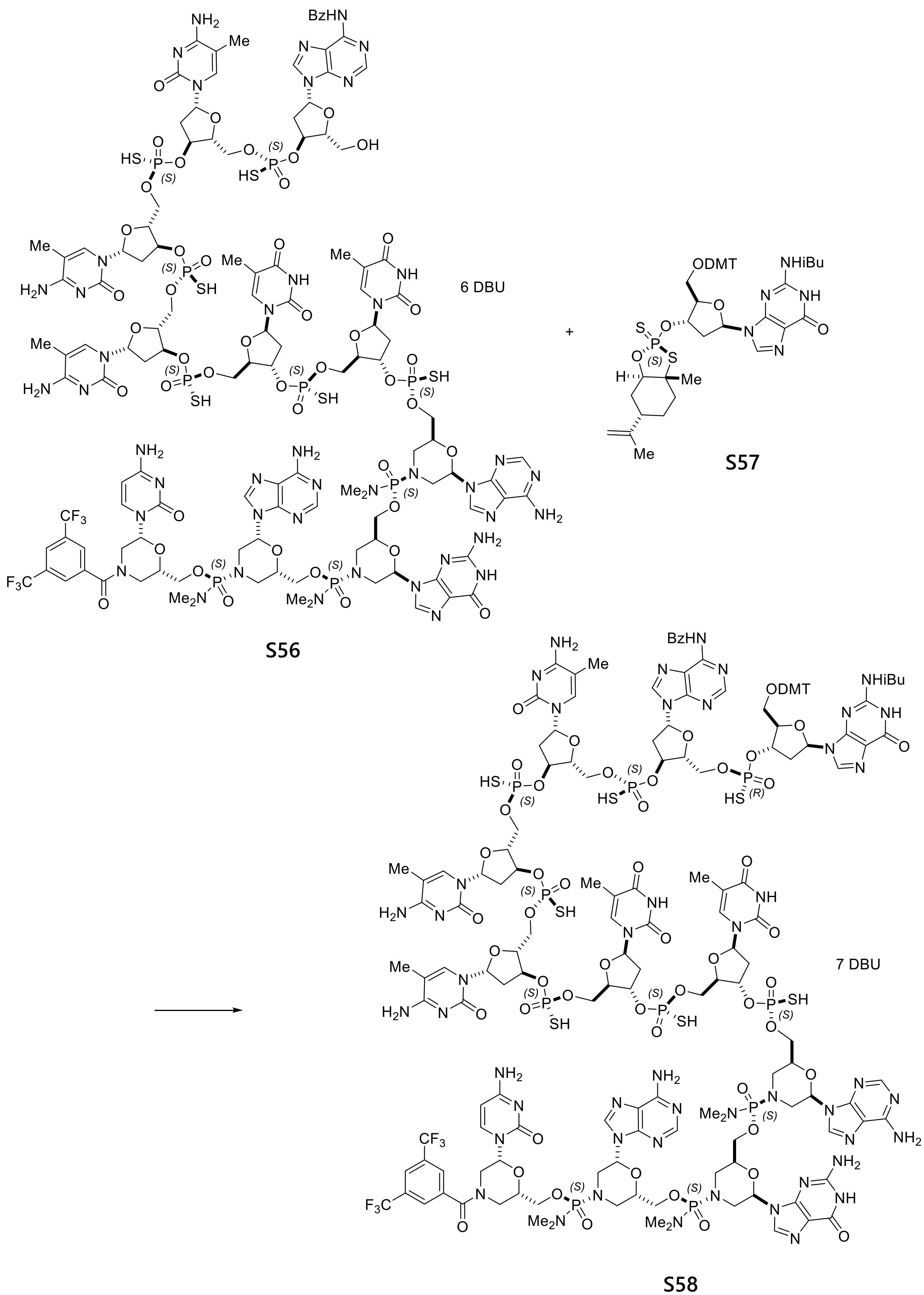

S56 S57 S58

To a solution of starting material **S56** (316 mg, 0.071 mmol) in 1,3-dimethyl-2-imidazolidinone (12.6 mL) was added reactant **S57** (189 mg, 0.213 mmol). To the resulting solution was added 4 Å molecular sieves (1.4 g). The reaction flask was applied to vacuum and filled with nitrogen. The process was repeated two more times. After being stirred for 30 min, the resulting mixture was treated with DBU (0.11 mL, 0.71 mmol). The reaction mixture was stirred at room temperature overnight and additional reactant **79** (92 mg) was added. After being stirred for 2 days, the reaction mixture was filtered through a syringe filter and the resulting filtrate was added into EtOAc (20 mL), rinsing with 1,3-dimethyl-2-imidazolidinone (3 mL). The resulting slurry mixture was centrifuged (3500 rpm, 30 min). The resulting pellet was treated with MeCN (20 mL) followed by EtOAc (20 mL). The resulting slurry was filtered through a glass filter and rinsed with MeCN/EtOAc (5 mL/ 5 mL). Drying the filter cake in vacuo at room temperature for 4 h provided 375 mg of target product **S58**.

$^{31}$P NMR (162 MHz, METHANOL-d4) δ = 57.27 (s, 1P), 56.95 (s, 1P), 56.91 (s, 1P), 56.83 (s, 1P), 56.81 (s, 1P), 56.75 (s, 1P), 56.24 (s, 1P), 16.95 (s, 2P), 16.71 (s, 1P).

MS (ESI) m/z: $[M-3H]^{3-}$ Calcd for $C_{156}H_{187}F_6N_{48}O_{54}P_{10}S_7$ 1414.96; Found 1414.94

11-mer: deprotection

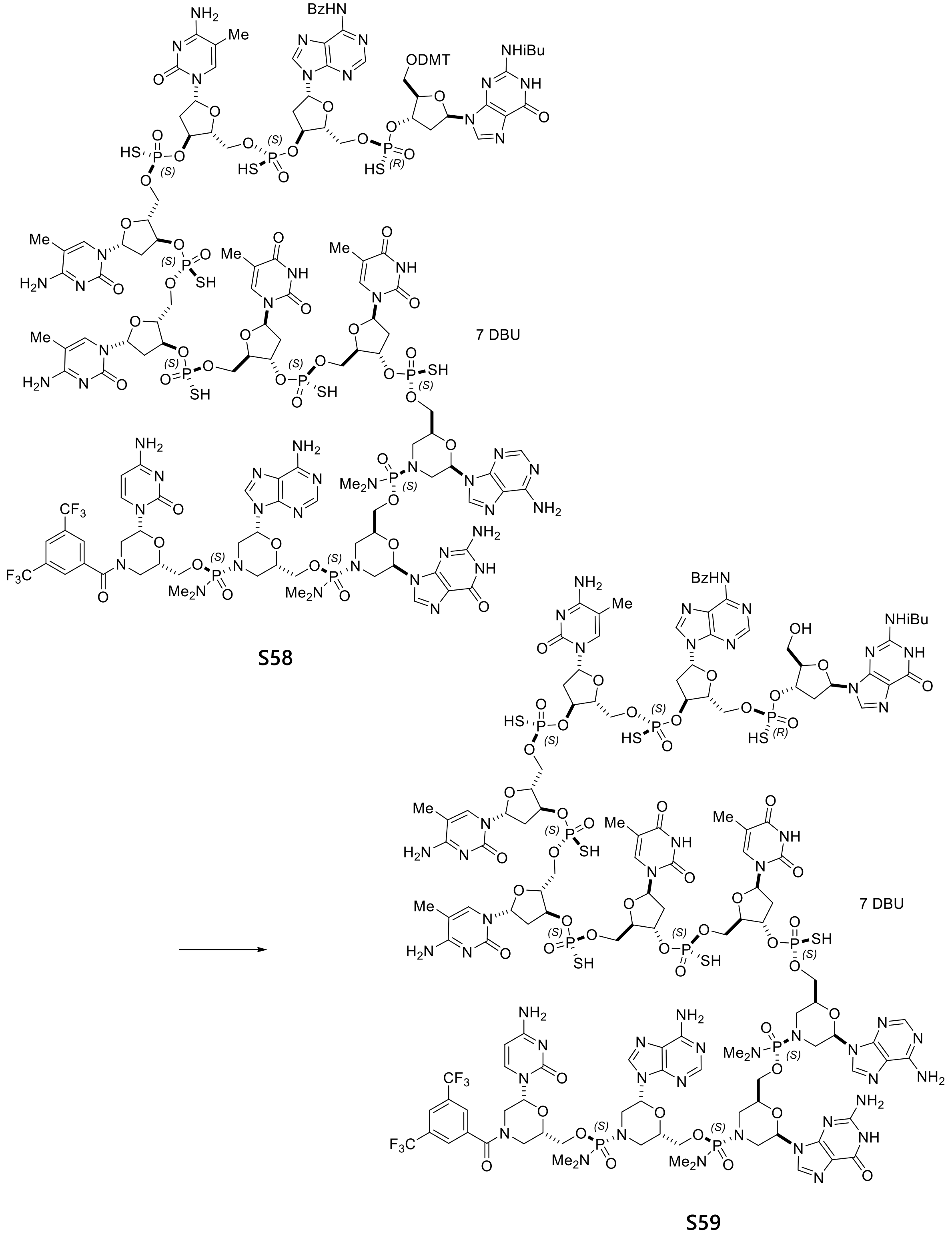

Starting material **S58** (375 mg, 0.071 mmol) was dissolved in a mixture of 1,1,1,3,3,3-hexafluoro-2-propanol (4.5 mL), 2,2,2-trifluoroethanol (1.1 mL), triethylsilane (3.4 mL) and $CH_2Cl_2$ (5.6 mL). The resulting solution was stirred at room temperature for 40 min and treated with EtOAc (75 mL) followed MTBE (25 mL). The resulting precipitate was collected by filtration and rinsed with a mixture of EtOAc/MeCN (10 mL/10 mL). The filter cake was treated with MeCN (20 mL) followed by EtOAc (20 mL). The resulting slurry was filtered through a filter and rinsed with MeCN/EtOAc (5 mL/5 mL). Drying the filter cake in vacuo overnight provided 343 mg of target product **S59**.

MS (ESI) m/z: $[M-2H]^{2-}$ Calcd for $C_{135}H_{170}F_6N_{48}O_{52}P_{10}S_7$ 1971.88; Found 1971.73.

12-mer: coupling

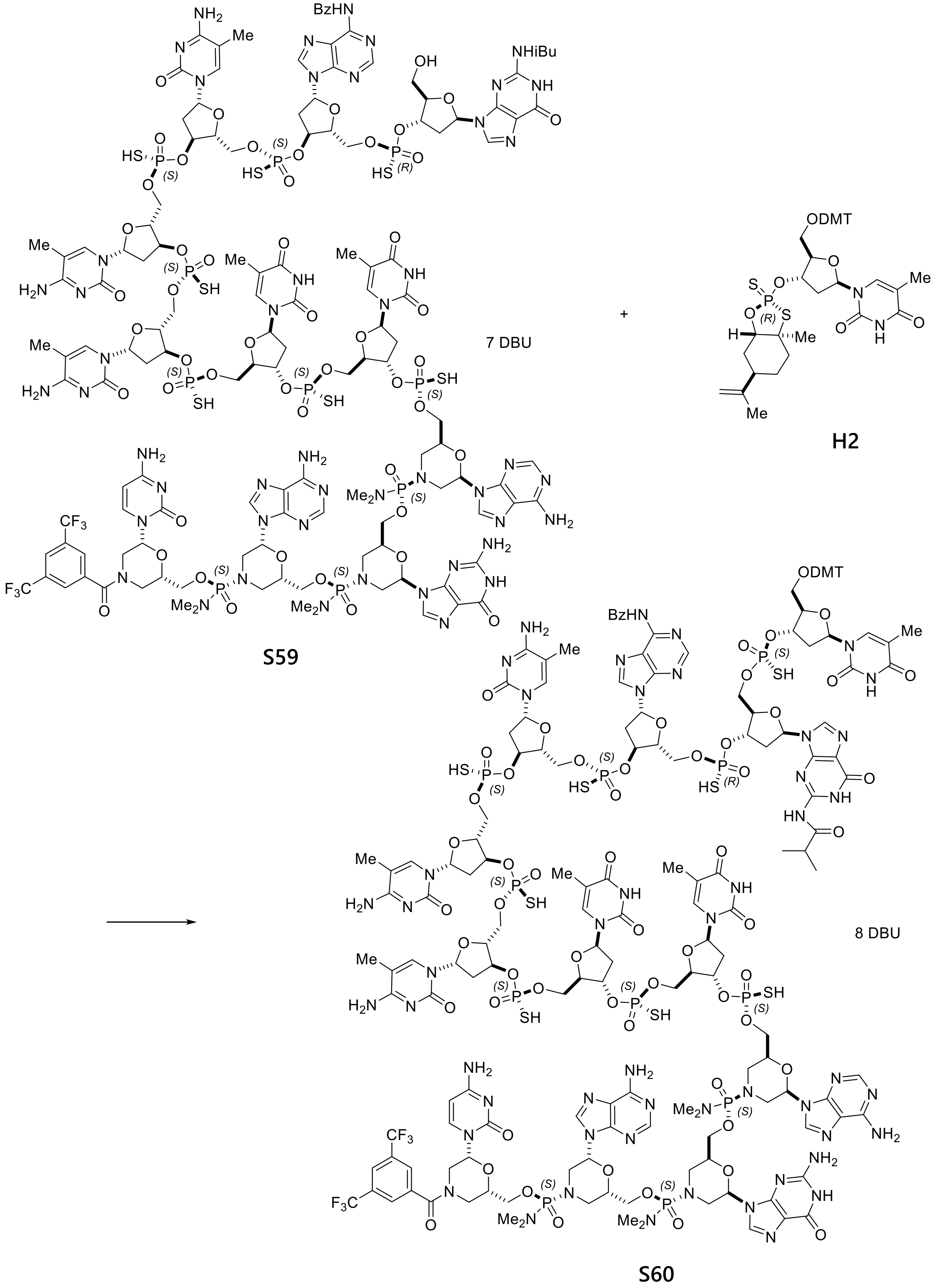

To a solution of starting material **S59** (343 mg, 0.068 mmol) in 1,3-dimethyl-2-imidazolidinone (12 mL) was added reactant **H2** (189 mg, 0.239 mmol). To the resulting solution was added 4 Å molecular sieves (1.5 g). The reaction flask was applied to vacuum and filled with nitrogen. The process was repeated two more times. After being stirred for 30 min, the resulting mixture was treated with DBU (0.113 mL, 0.753 mmol). The reaction mixture was stirred for 23 h at room temperature and then filtered through a syringe filter. The filtrate was added into EtOAc (20 mL), rinsing with 1,3-dimethyl-2-imidazolidinone (5 mL). Additional EtOAc (20 mL) was added. The resulting slurry was centrifuged (3500 rpm, 30 min). The resulting pellet was treated with MeCN (20 mL) followed by EtOAc (20 mL). The resulting slurry was filtered through a glass filter and rinsed with MeCN/EtOAc (5 mL/5 mL). Drying the filter cake in vacuo at room temperature for 3 h provided target product **S60**.

$^{31}$P NMR (162 MHz, METHANOL-d4) δ = 57.28 (s, 1P), 57.24 (s, 1P), 56.94 (s, 1P), 56.81 (s, 2P), 56.74 (s, 2P), 56.22 (s, 1P), 16.95 (s, 2P), 16.70 (s, 1P)

MS (ESI) m/z: $[M-3H]^{3-}$ Calcd for $C_{166}H_{200}F_6N_{50}O_{60}P_{11}S_8$ 1521.63; Found 1521.41

12-mer: deprotection

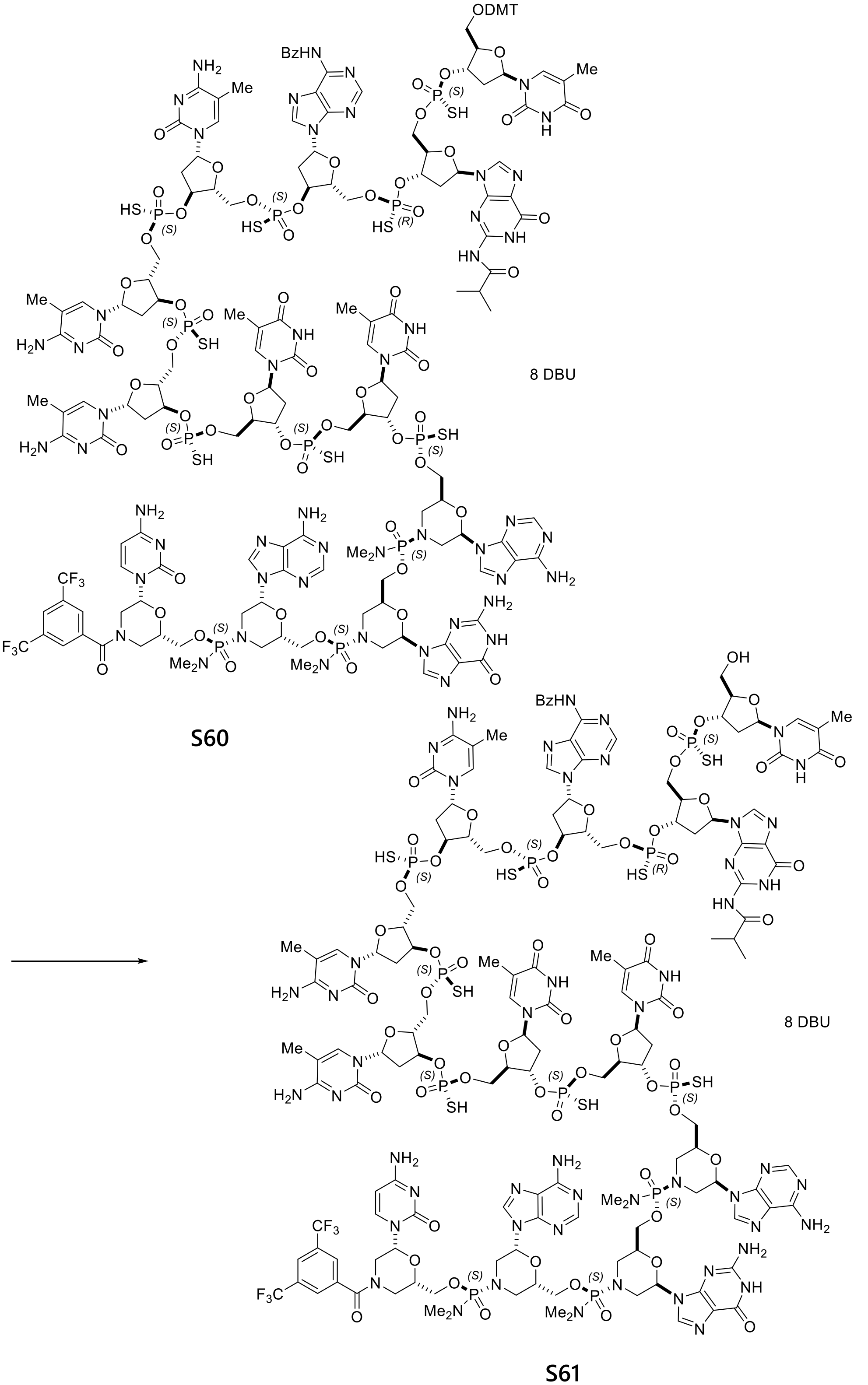

S60

S61

Starting material **S60** (396 mg, 0.068 mmol in theory) was dissolved in a mixture of 1,1,1,3,3,3-hexafluoro-2-propanol (4.8 mL), 2,2,2-trifluoroethanol (1.2 mL), triethylsilane (3.6 mL) and $CH_2Cl_2$ (6.0 mL). The resulting solution was stirred at room temperature for 16 h and treated with EtOAc (100 mL). The resulting precipitate was collected by filtration and rinsed with a mixture of EtOAc/MeCN (5 mL/5 mL). The filter cake was treated with MeCN (20 mL) followed by EtOAc (20 mL). The resulting slurry was filtered through a glass filter and rinsed with MeCN/EtOAc (5 mL/5 mL). Drying the filter cake in vacuo for 1h provided 310 mg of target product **S61**.

MS (ESI) m/z: $[M-3H]^{3-}$ Calcd for $C_{145}H_{182}F_6N_{50}O_{58}P_{11}S_8$ 1421.26; Found 1421.32.

#### 13-mer: coupling

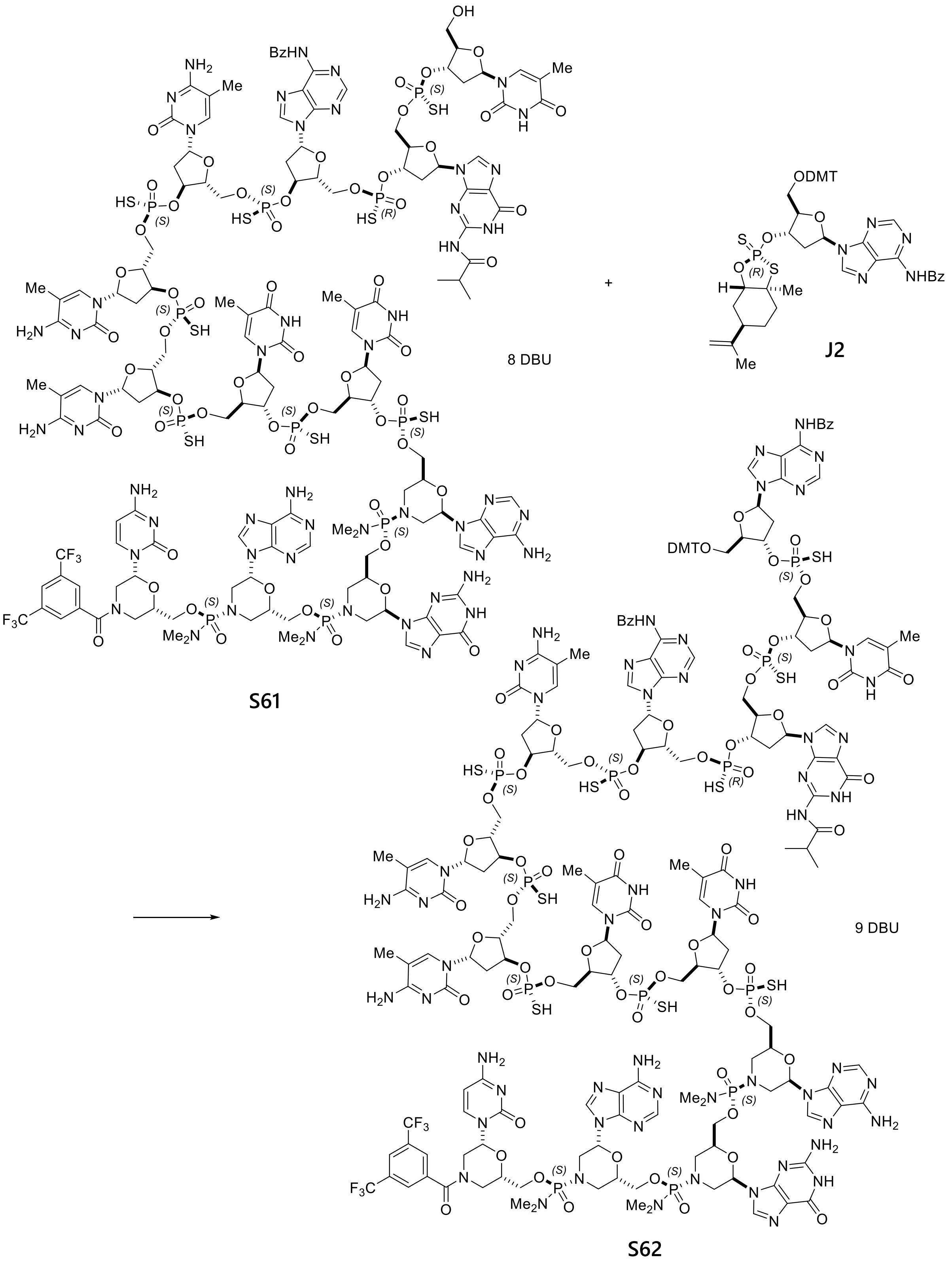

To a solution of starting material **S61** (310 mg, 0.057 mmol) in 1,3-dimethyl-2-imidazolidinone (11 mL) was added reactant **J2** (179 mg, 0.198 mmol). To the resulting solution was added 4 Å molecular sieves (1.2 g). The reaction flask was applied to vacuum and filled with nitrogen. The process was repeated two more times. fter being stirred for 30 min, the resulting mixture was treated with DBU (0.102 mL, 0.678 mmol). The reaction mixture was stirred overnight at room temperature and then filtered through a syringe filter. The filtrate was added into EtOAc (20 mL), rinsing with 1,3-dimethyl-2-imidazolidinone (5 mL). Additional EtOAc (20 mL) was added. The resulting slurry was centrifuged (3500 rpm, 30 min). The resulting pellet was treated with MeCN (20 mL) followed by EtOAc (20 mL). The resulting slurry was filtered through a glass filter and rinsed with MeCN/EtOAc (5 mL/5 mL). The filter cake was dried in vacuo at room temperature for 3 days, and then treated with 25 mL MeCN to make a slurry. After being stirred for 30 min, the resulting slurry was filtered through a glass filter and rinsed with MeCN/EtOAc (5 mL/5 mL). Drying the filter cake in vacuo for 1 h provided 365 mg of target product **S62**.

$^{31}$P NMR (162 MHz, METHANOL-d4) δ = 57.22 (s, 1P), 56.96 (s, 2P), 56.89 (s, 1P), 56.78 (s, 2P), 56.74 (s, 2P), 56.27 (s, 1P), 16.96 (s, 2P), 16.72 (s, 1P).

MS (ESI) m/z: $[M-3H]^{3-}$ Calcd for $C_{183}H_{216}F_6N_{55}O_{65}P_{12}S_9$ 1666.32; Found 1666.24.

13-mer: deprotection

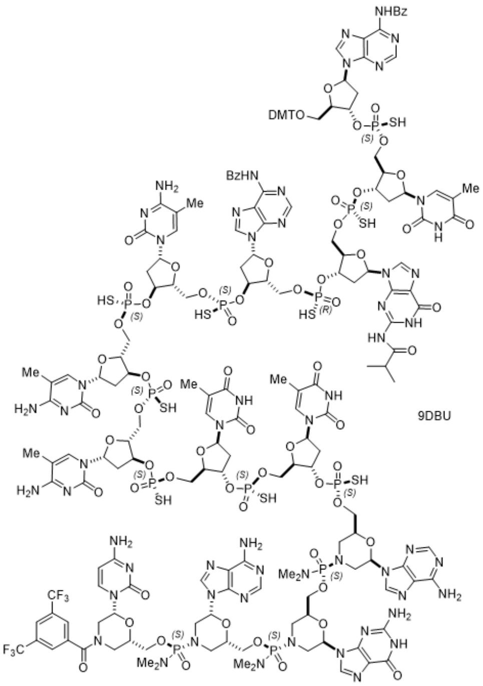

**S62**

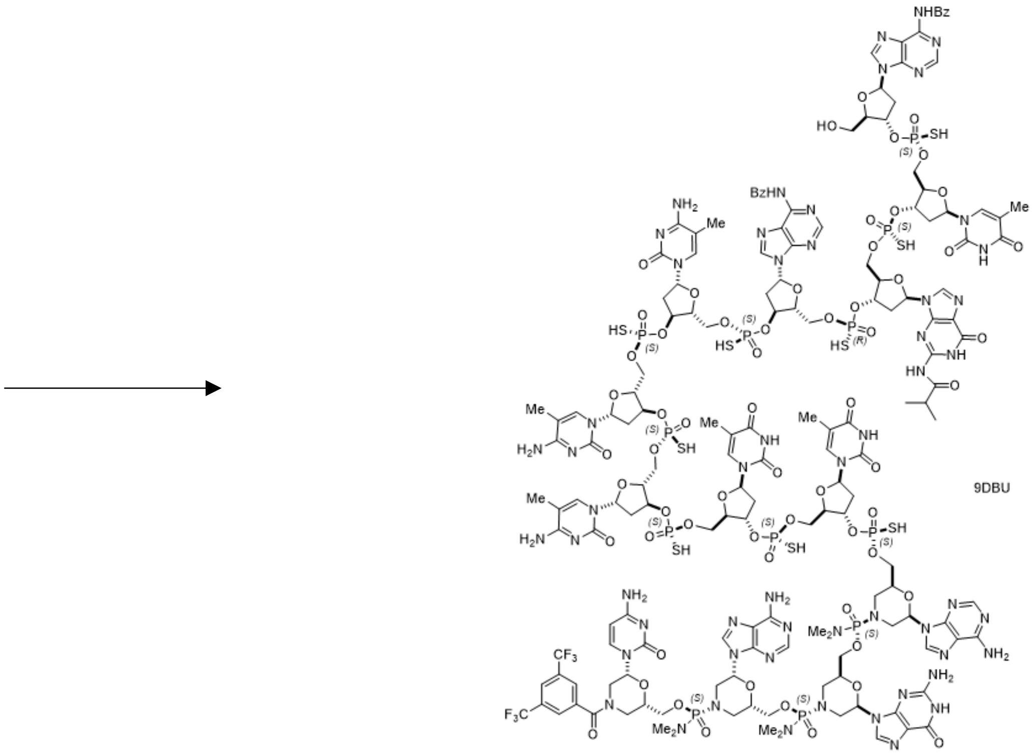

**S63**

Starting material **S62** (365 mg, 0.057 mmol) was dissolved in a mixture of 1,1,1,3,3,3-hexafluoro-2-propanol (4.4 mL), 2,2,2-trifluoroethanol (1.1 mL), triethylsilane (3.3 mL) and $CH_2Cl_2$ (5.5 mL). The resulting solution was stirred at room temperature for 20 min and treated with 125 mL EtOAc. The resulting precipitate was collected by filtration and rinsed with a mixture of EtOAc/MeCN (10 mL/10 mL). The filter cake was treated with MeCN (20 mL) followed by EtOAc (10 mL). The resulting slurry was centrifuged (4000 rpm, 60 min). The resulting pellet was isolated by decantation and rinsed with MeCN/EtOAc (5 mL/5 mL). Drying in vacuo overnight provided 328 mg of target product **S63**.

MS (ESI) m/z: $[M-3H]^{3-}$ Calcd for $C_{162}H_{198}F_6N_{55}O_{63}P_{12}S_9$ 1565.61; Found 1565.65.

#### 13+5 Block coupling

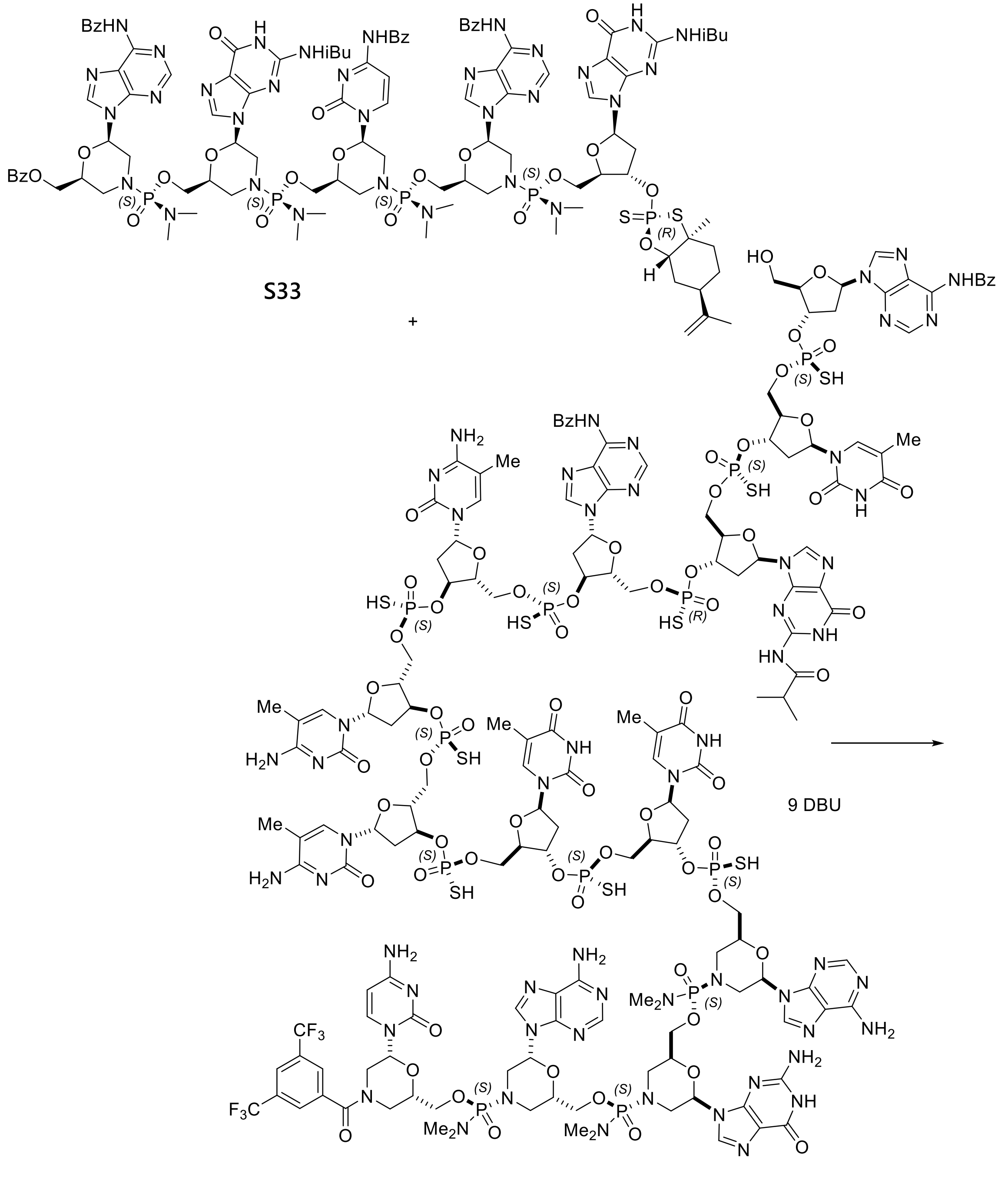

S63

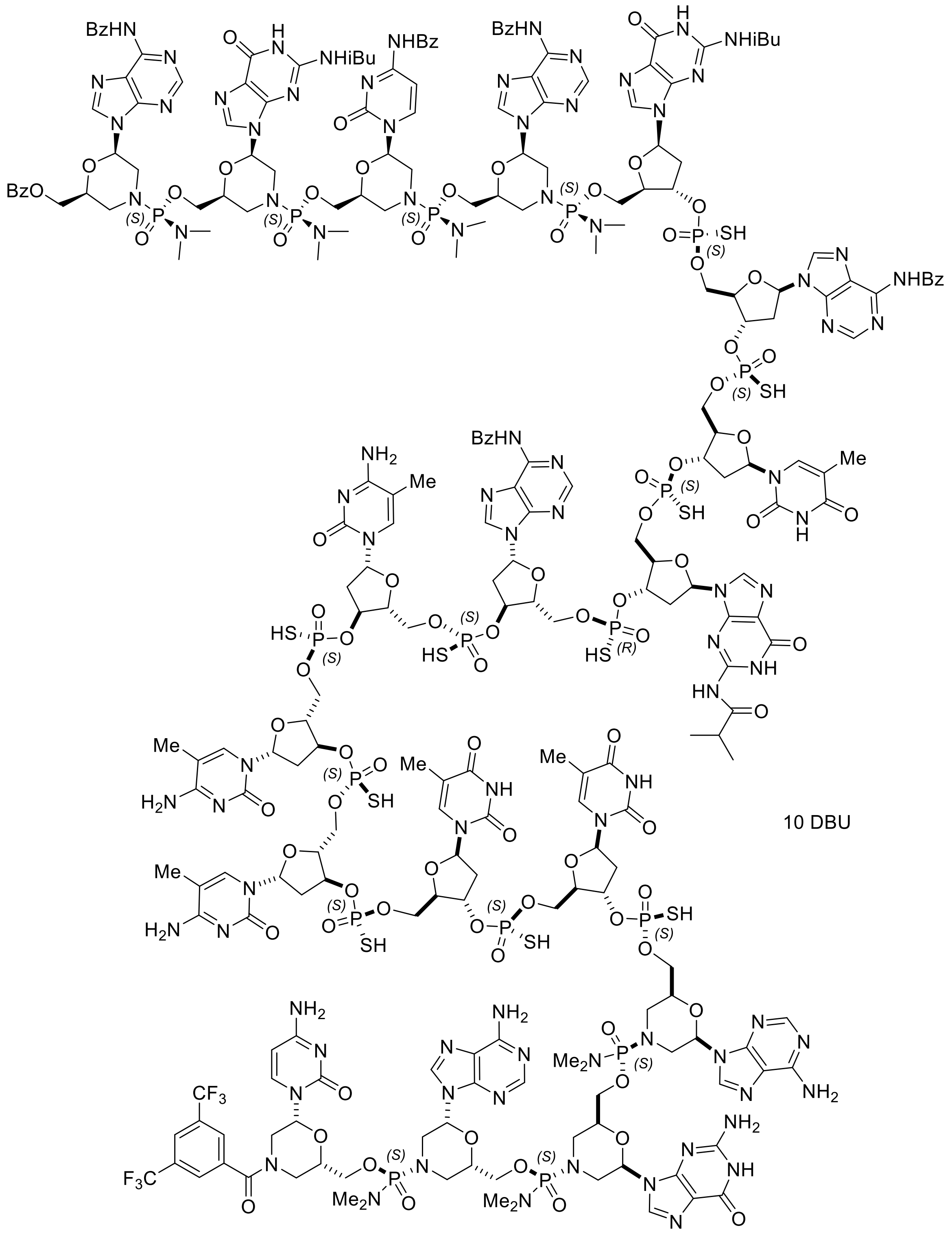

**S65**

To a mixture of starting material **S63** (100 mg, 0.016 mmol) and reactant **S33** (139 mg, 0.058 mmol) was added 1,3-dimethyl-2-imidazolidinone (3.5 mL). The resulting mixture was azeotroped with toluene (2 mL each time) three times at 30-33 ºC. To the resulting solution was added 4 Å molecular sieves (0.40 g). The reaction flask was applied to vacuum and filled with nitrogen. The process was repeated two more times. After being stirred for 30 min, the resulting mixture was treated with DBU (0.032 mL, 0.21 mmol). The reaction mixture was stirred for 3 days at room temperature and then filtered through a syringe filter. The filtrate was added into EtOAc (15 mL), rinsing with 1,3-dimethyl-2-imidazolidinone (2.5 mL). The resulting slurry was centrifuged (3500 rpm, 20 min). The pellet was dissolved in EtOH (3 mL) and $CH_2Cl_2$ (6 mL). To the resulting solution was added EtOAc (20 mL). The resulting slurry was filtered through a glass filter and rinsed with MeCN (10 mL). Drying the filter cake in vacuo at room temperature for 0.5 h provided 0.13 g of target product **S65.**

$^{31}P$ NMR (162 MHz, METHANOL-d4) δ = 57.30 (s, 1P), 57.19 (s, 1P), 56.91 (s, 2P), 56.80 (s, 2P), 56.73 (s, 2P), 56.62 (s, 1P), 56.18 (s, 1P), 17.07 (s, 2P), 16.94 (s, 2P), 16.91 (s, 1P), 16.85 (s, 1P), 16.67 (s, 1P)

MS (ESI) m/z: $[M-4H]^{4-}$ Calcd for $C_{255}H_{309}F_6N_{86}O_{88}P_{17}S_{10}$ 1736.38; Found 1736.31.

**Final deprotection**

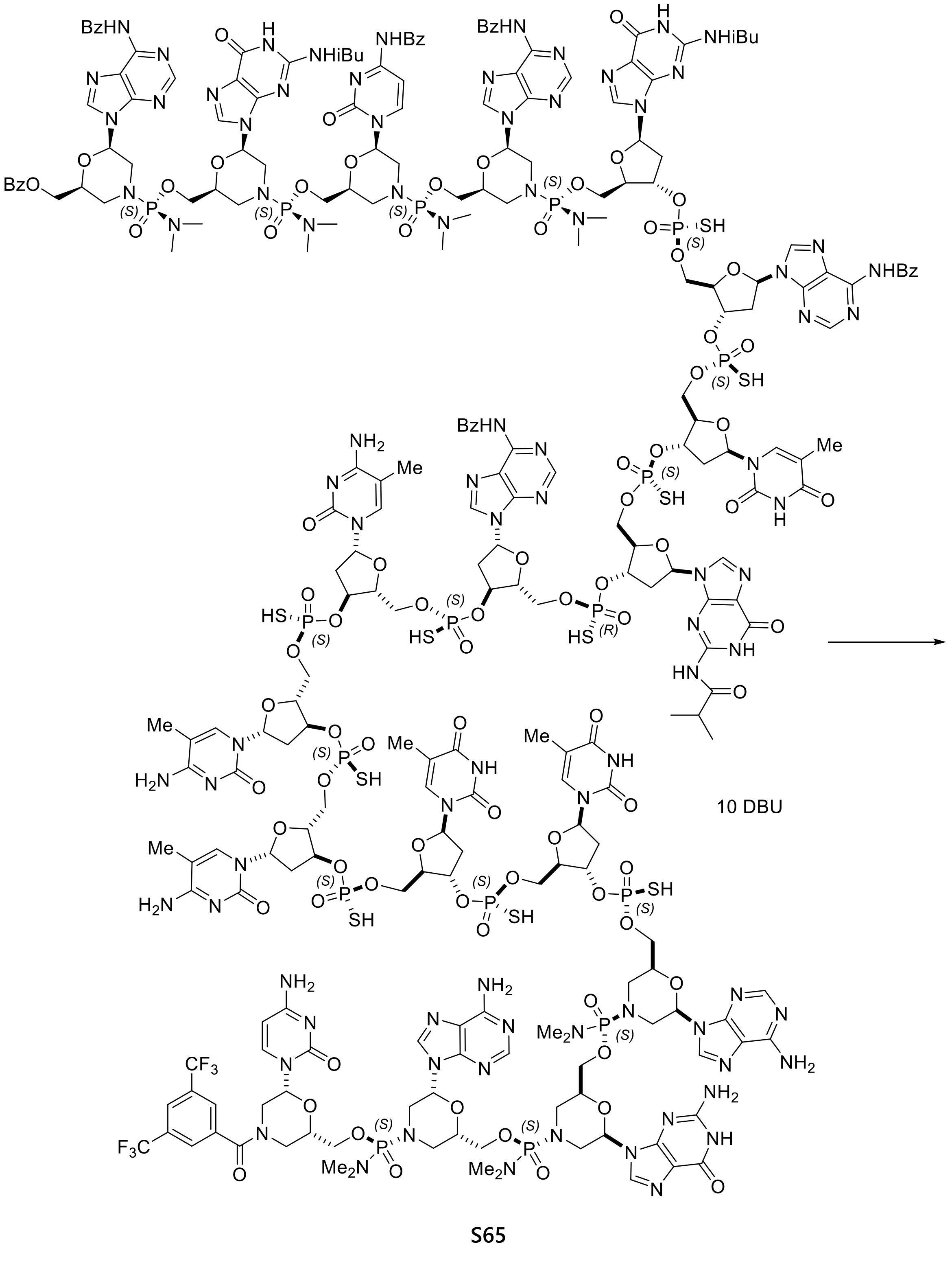

**S65**

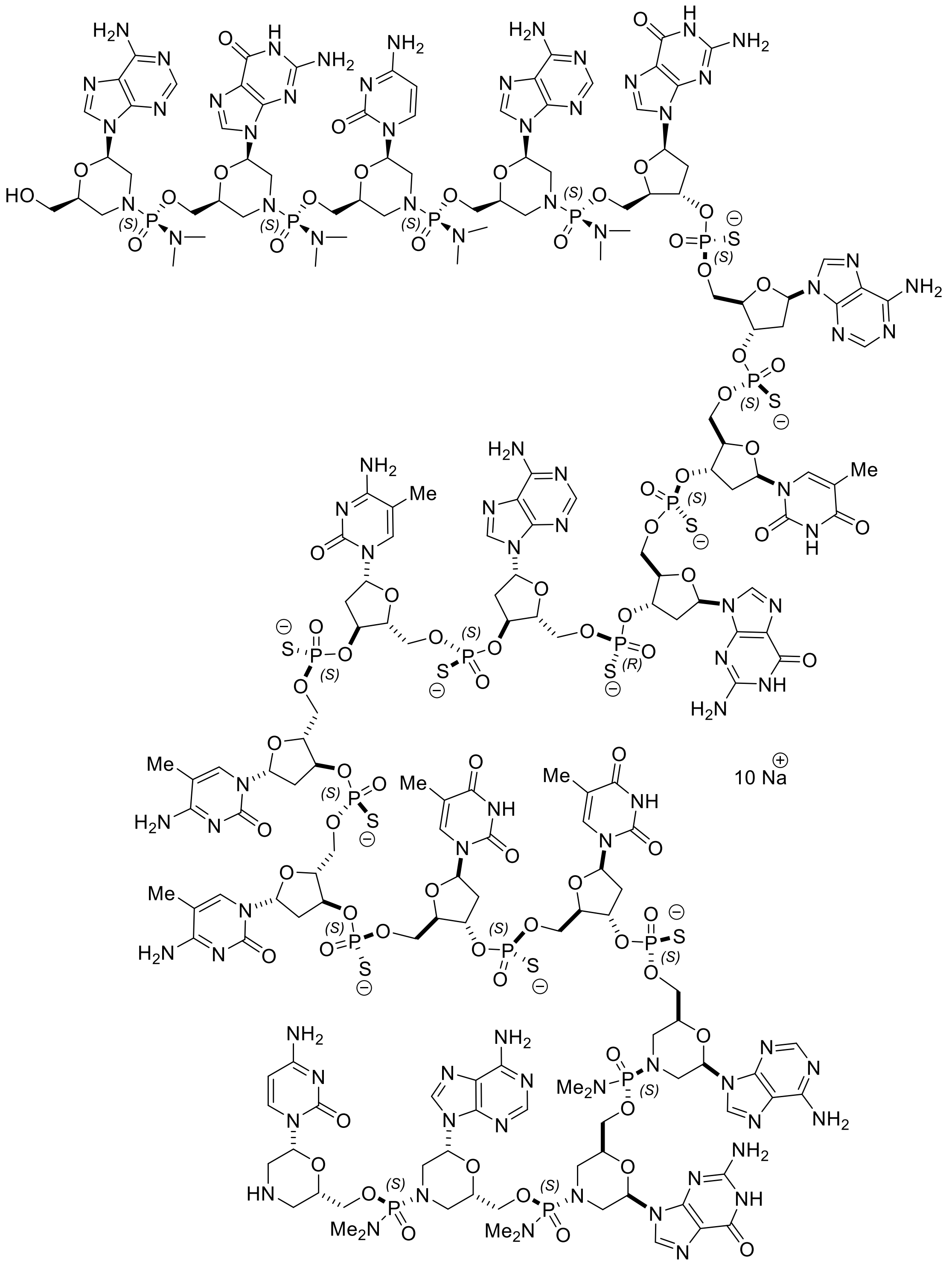

**ASO-486-R5-S**

To a solution of starting material **S65** (0.130 mg, 0.015 mmol) in a mixture of methanol (4.6 mL) and 28% ammonium hydroxide (4.6 mL) was added DL-dithiothreitol (0.024 g, 0.15 mmol). The resulting mixture was stirred at 53-55 ºC for 23 h and cooled to room temperature. A mixture of MeCN/EtOAc (20 mL/20 mL) was added and the resulting slurry was subjected to centrifuge (4000 rpm, 90 min). The resulting pellet was isolated and dissolved in water (30 mL). The aqueous solution was subjected to ultrafiltration (Amicon Ultra-15, ultracel 3K, 3500 rpm, 35 min). The remaining solution was diluted with water (30 mL) and subjected to ultrafiltration (Amicon Ultra-15, ultracel 3K, 3500 rpm, 35 min). The remaining solution was filtered through a syringe filter and rinsed with water. The filtrate (ca. 5 mL) was subjected to centrifuge (4000 rpm, 30 min) and the supernatant was purified by prep-HPLC using the conditions in **Table S5** and then the conditions in **Table S6**.

**Table S5**: RP-HPLC conditions

| | | | | | |
|---|---|---|---|---|---|
| Column | Waters, XBridge Prep C18 5µm OBD, 19x100mm (Part Number: 186002978) | | | | |
| Instrument | Waters 2545 Binary Gradient Module, Waters 3100 Mass Detector | | | | |
| Mobile phase A | 100 mM HFIP (Hexafluoroisopropanol) + 8.6 mM TEA (Triethylamine) in water | | | | |
| Mobile phase B | Methanol 100% | | | | |
| Column Temperature (ºC) | 60 | | | | |
| Gradient | | | | | |
| | TIME (min) | A% | B% | Flow rate (mL/min) | comments |
| | 0 | 90 | 10 | 25 | Initial |
| | 2.2 | 90 | 10 | 25 | |
| | 4.4 | 80 | 20 | 30 | Elution Gradient |
| | 11.1 | 50 | 50 | 30 | |
| | 11.2 | 0 | 100 | 30 | Wash |
| | 17.9 | 0 | 100 | 30 | |
| | 18.0 | 90 | 10 | 30 | Reset Conditions |
| | 20.2 | 90 | 10 | 30 | |
| Flow Rate (mL/min) | See the table | | | | |
| Wavelength (nm) | 260 | | | | |

**Table S6**: IEX-HPLC conditions

| | | | | |
|---|---|---|---|---|
| Column | TOSOH Bioscience, TSKgel SuperQ-5PW, 7.5mm ID x 7.5cm, 10µm (Part No: 0018257) | | | |
| Instrument | Agilent 1200 | | | |
| Mobile phase A | 10 mM NaOH in water | | | |
| Mobile phase B | 10 mM NaOH + 1M NaCl in water | | | |
| Column Temperature (°C) | 45 | | | |
| Gradient: | | | | |
| | TIME (min) | A% | B% | comments |
| | 0 | 50 | 50 | Initial |
| | 1.7 | 30 | 70 | Elution Gradient |
| | 11.6 | 0 | 100 | Wash |
| | 13.3 | 0 | 100 | |
| | 13.4 | 50 | 50 | Reset Conditions |
| | 15.1 | 50 | 50 | |
| Flow Rate (mL/min) | 2.0 | | | |
| Wavelength (nm) | 260 | | | |

Desalting of the purified product was conducted 4 times with Amicon Ultra-15, Ultracel-3K (3500 rpm, 45 min). Freeze-drying of the resulting solution (12.5 mL) for 2 days provided 18 mg of target product **ASO-486-R5-S**.

HRMS (ESI) m/z: $[M-3H]^{3-}$ Calcd for $C_{192}H_{266}N_{86}O_{78}P_{17}S_{10}$ 1957.7415; Found 1957.7418.

**Preparation of Compound ASO-486-R5-R**

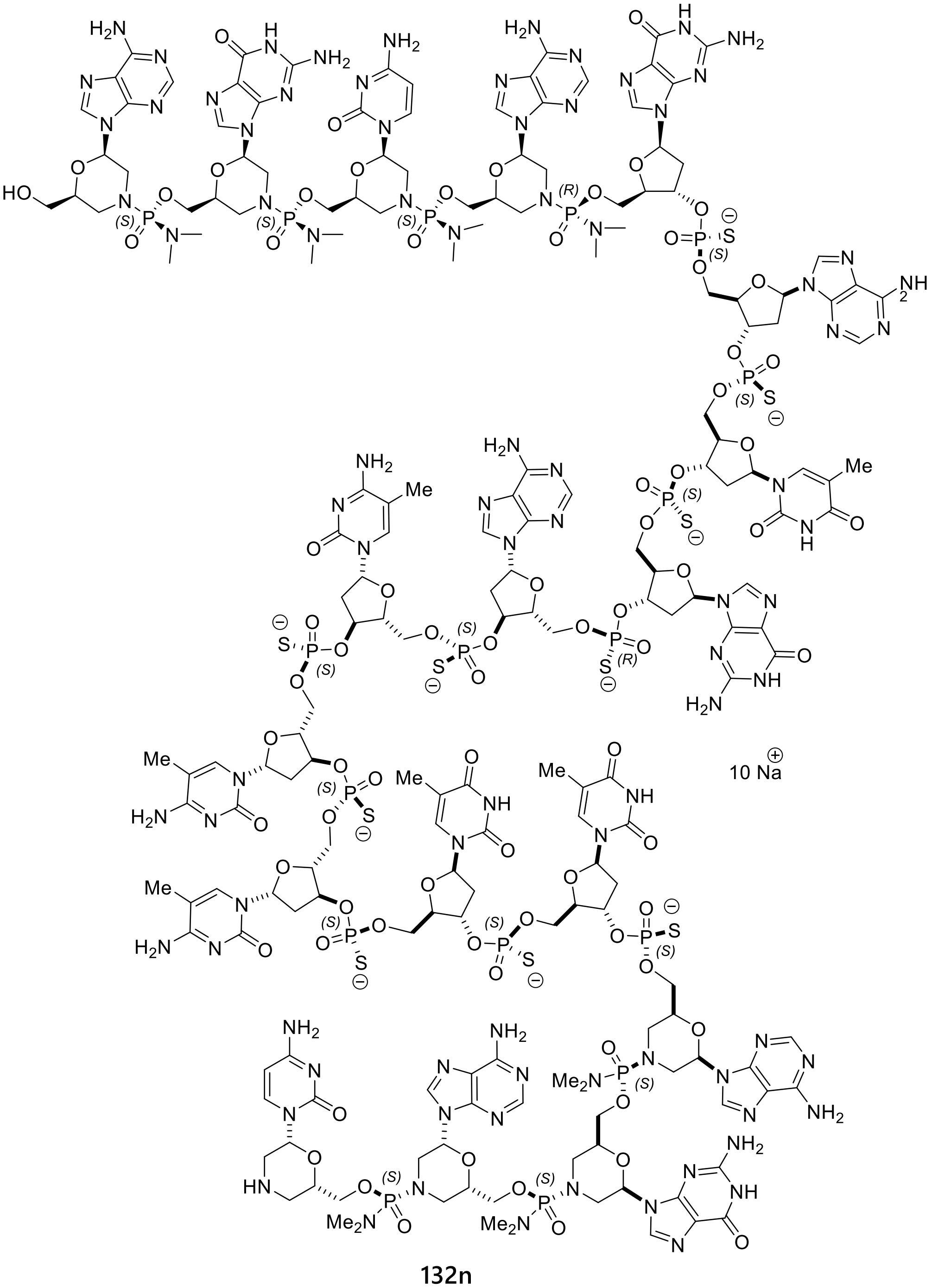

**132n**

With compound **S30b** instead of compound **S30a** in the preparation of the 5' wing 5-mer (compound **S31**), **ASO-486-R5-R** was prepared via the same reaction sequences as described for **ASO-486-R5-S**.

HRMS (ESI) m/z: $[M-3H]^{3-}$ Calcd for $C_{192}H_{266}N_{86}O_{78}P_{17}S_{10}$ 1957.7415; Found 1957.7422.

#### Preparation of Compound ASO-486-R5-M

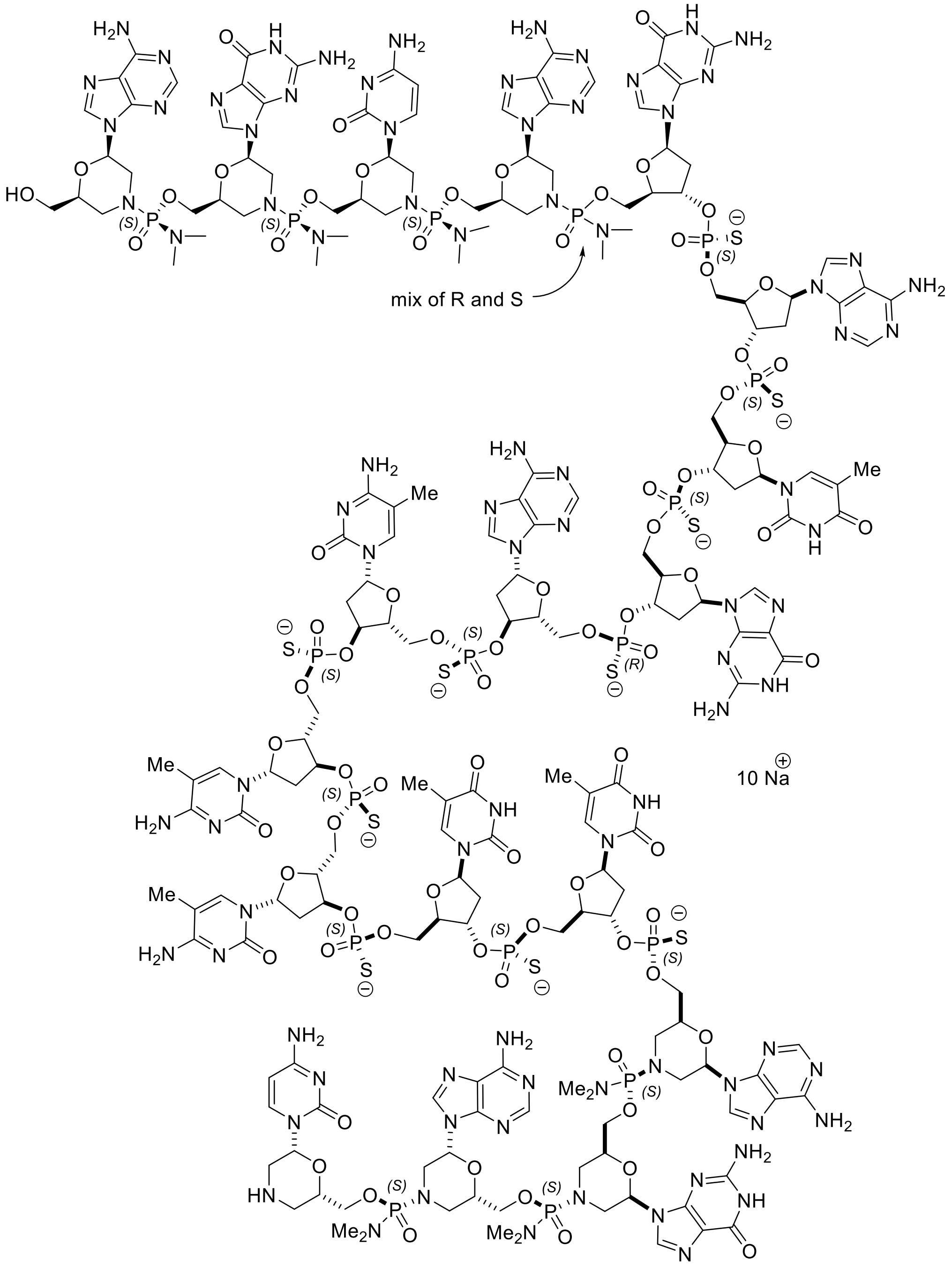

**132f**

With ((2R,3S,5R)-3-(bis(4-methoxyphenyl)(phenyl)methoxy)-5-(2-isobutyramido-6-oxo-1,6-dihydro-9H-purin-9-yl)tetrahydrofuran-2-yl)methyl dimethylphosphoramidochloridate (**S30**) instead of compound **S30a** in the preparation of the 5' wing 5-mer (compound **S31**), **ASO-**

**486-R5-M** was prepared via the same reaction sequences as described for Compound **ASO-486-R5-S**.

HRMS (ESI) m/z: $[M-3H]^{3-}$ Calcd for $C_{192}H_{266}N_{86}O_{78}P_{17}S_{10}$ 1957.7415; Found 1957.7439.

#### 5. Cited Documents

1. WO2018057430A1.
2. U.S. Patent No. 10,457,698.
3. U.S. Patent No. 10,836,784.
4. Bennett,C.F. (2019) Therapeutic Antisense Oligonucleotides Are Coming of Age. *Annual Review of Medicine*, **70**, 307–321.

#### 6. $^{31}$P NMR spectra of selected compounds (162 MHz, Methanol-d4)

**Compound S43 (4-mer)**

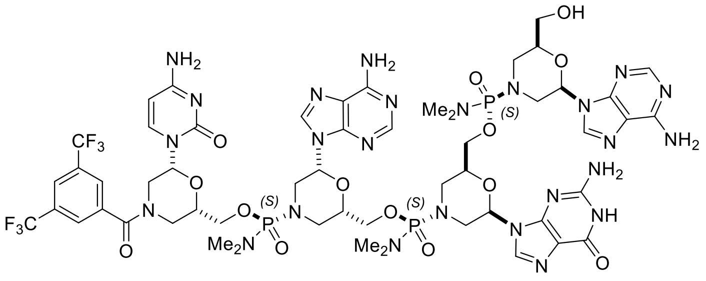

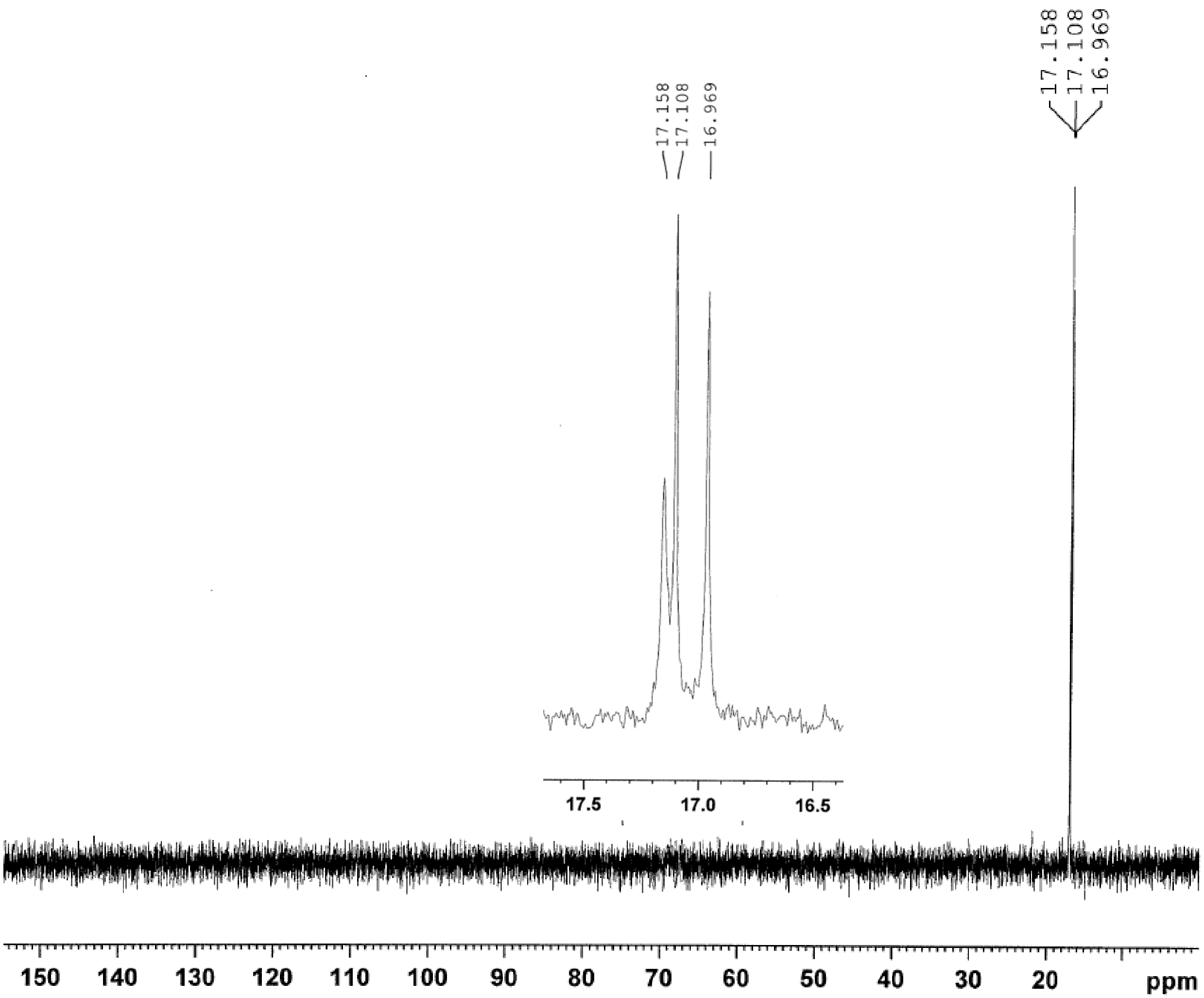

**Compound S45 (5-mer)**

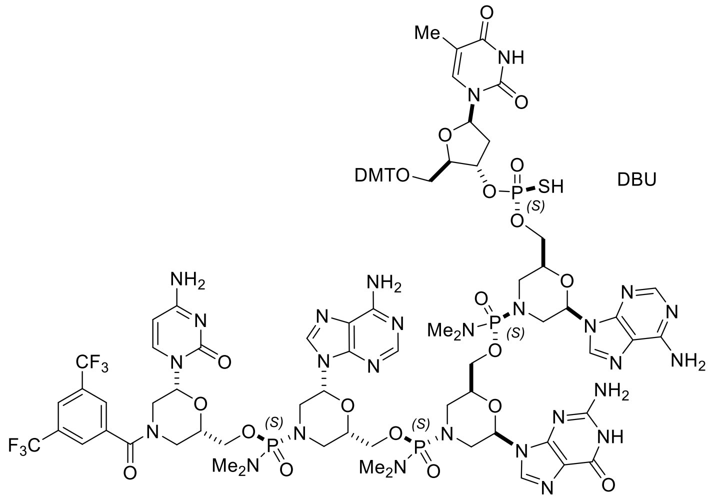

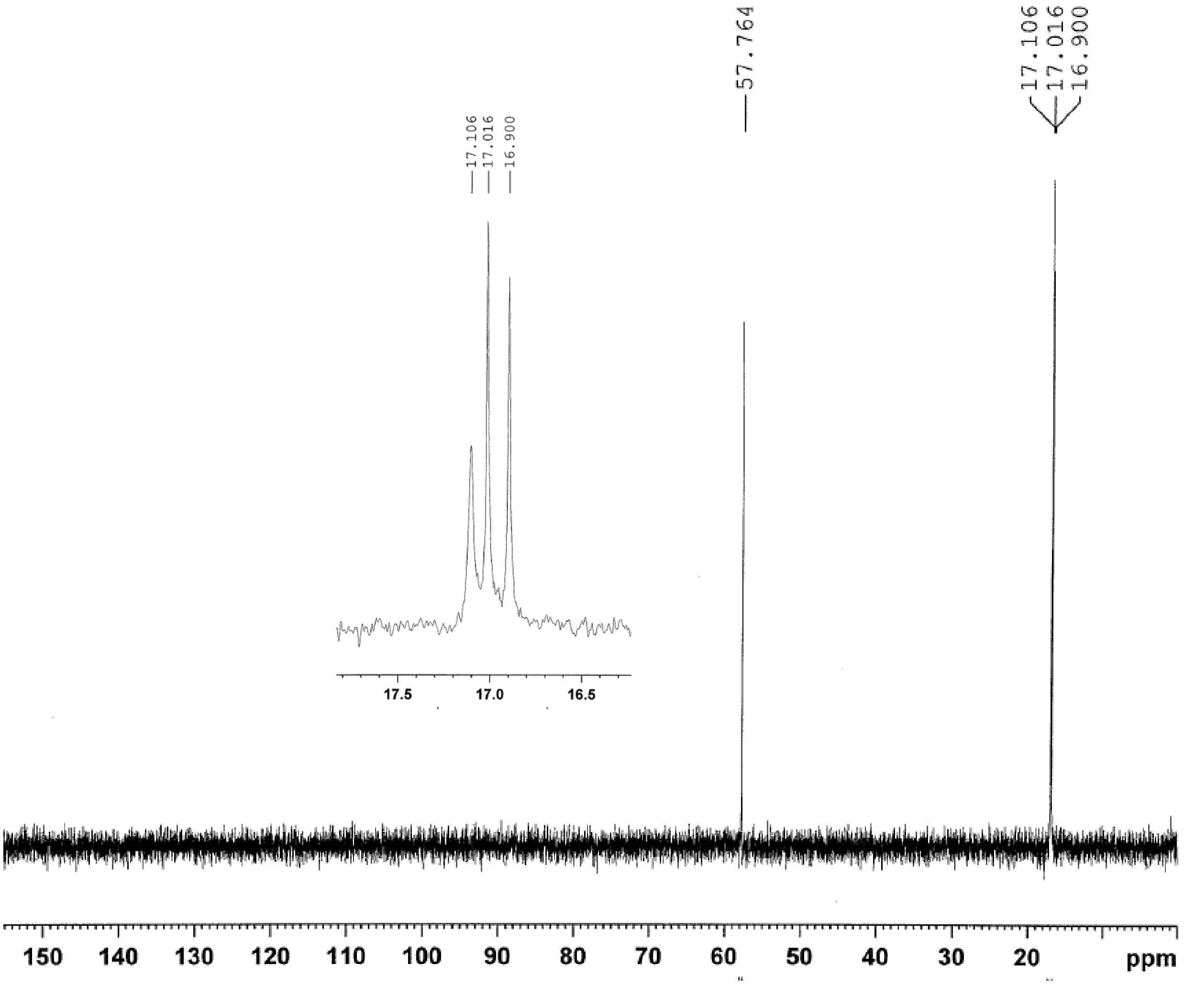

**Compound S49 (7-mer)**

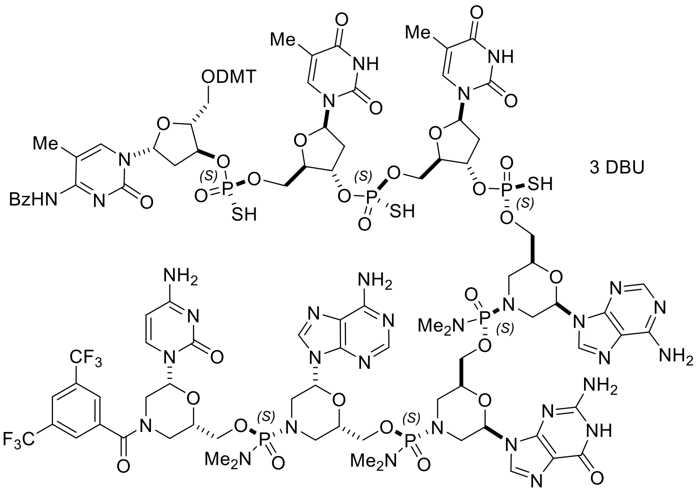

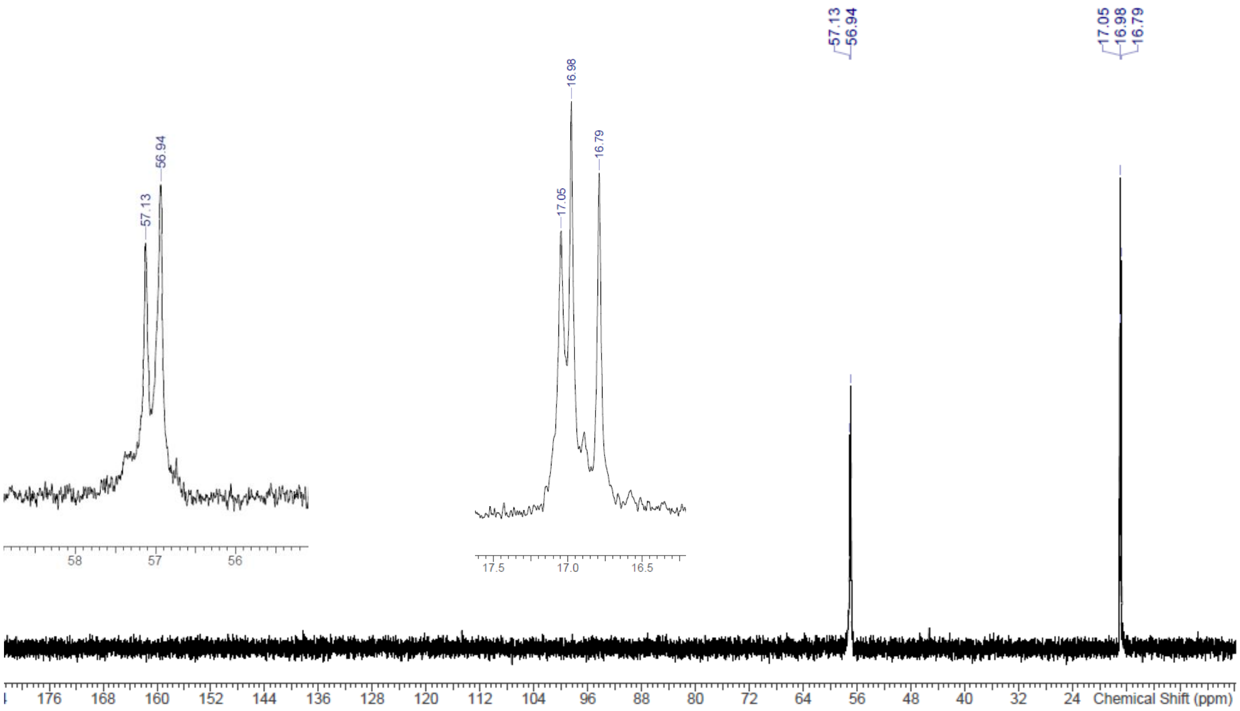

#### Compound S51 (8-mer)

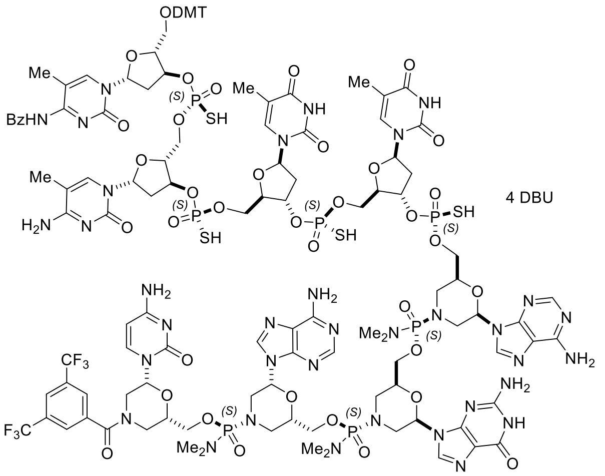

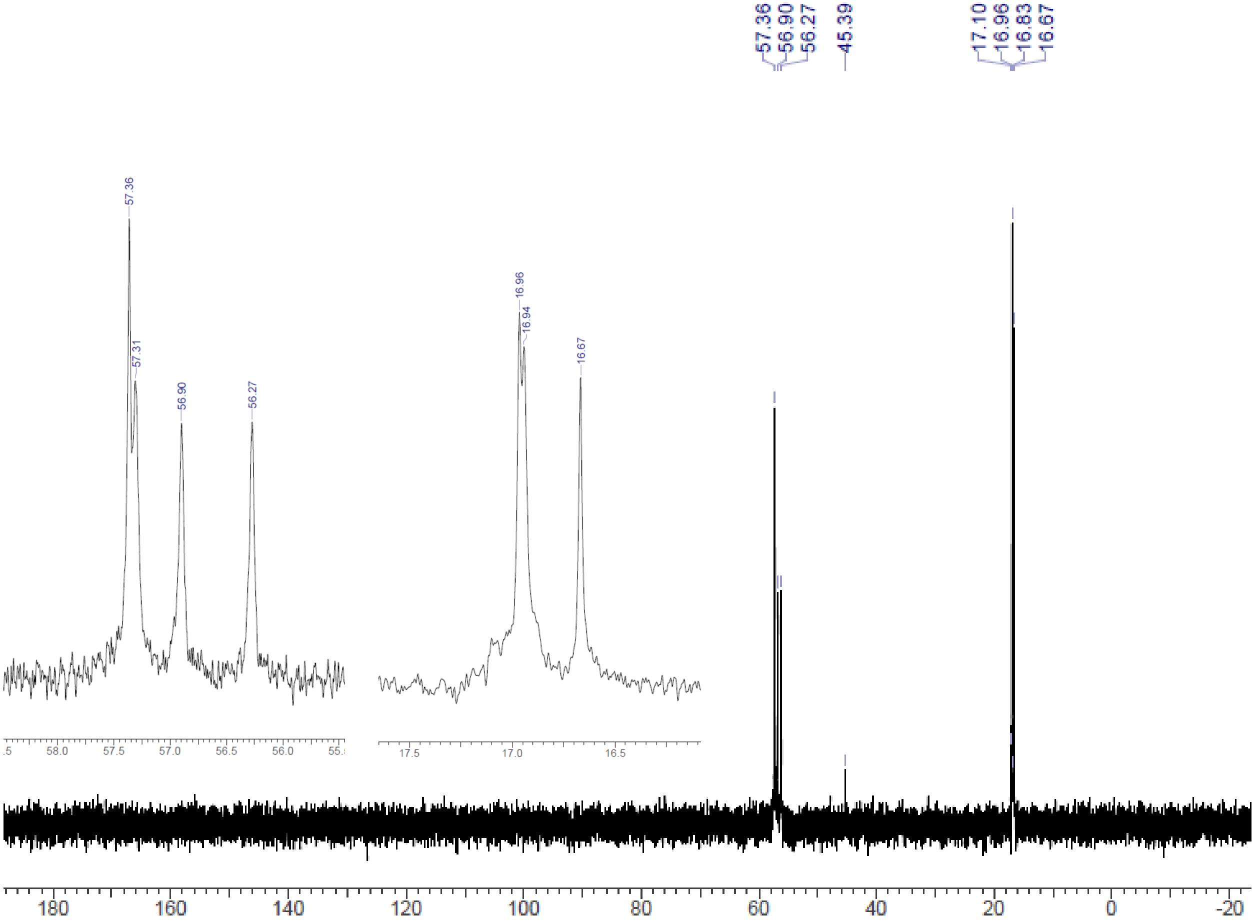

#### Compound S53 (9-mer)

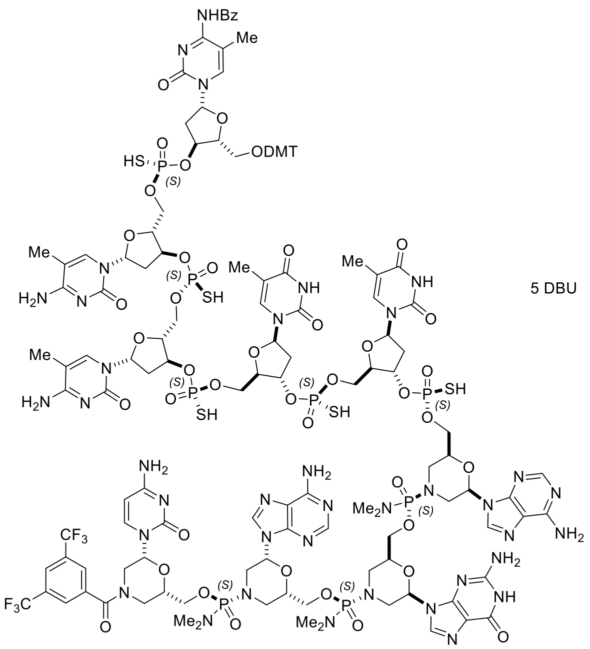

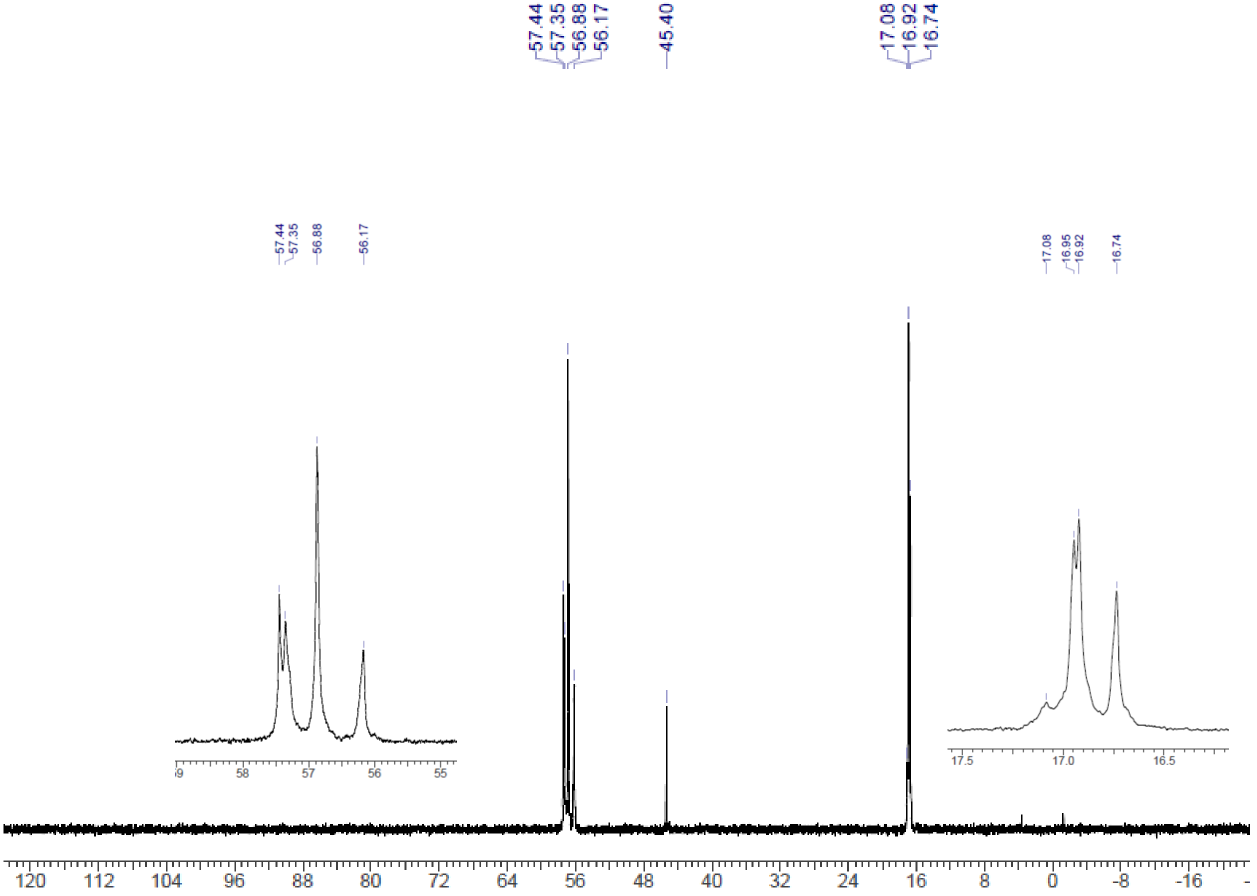

#### Compound S55 (10-mer)

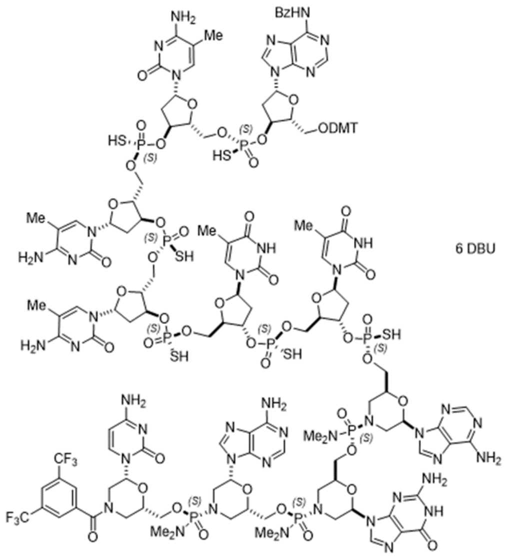

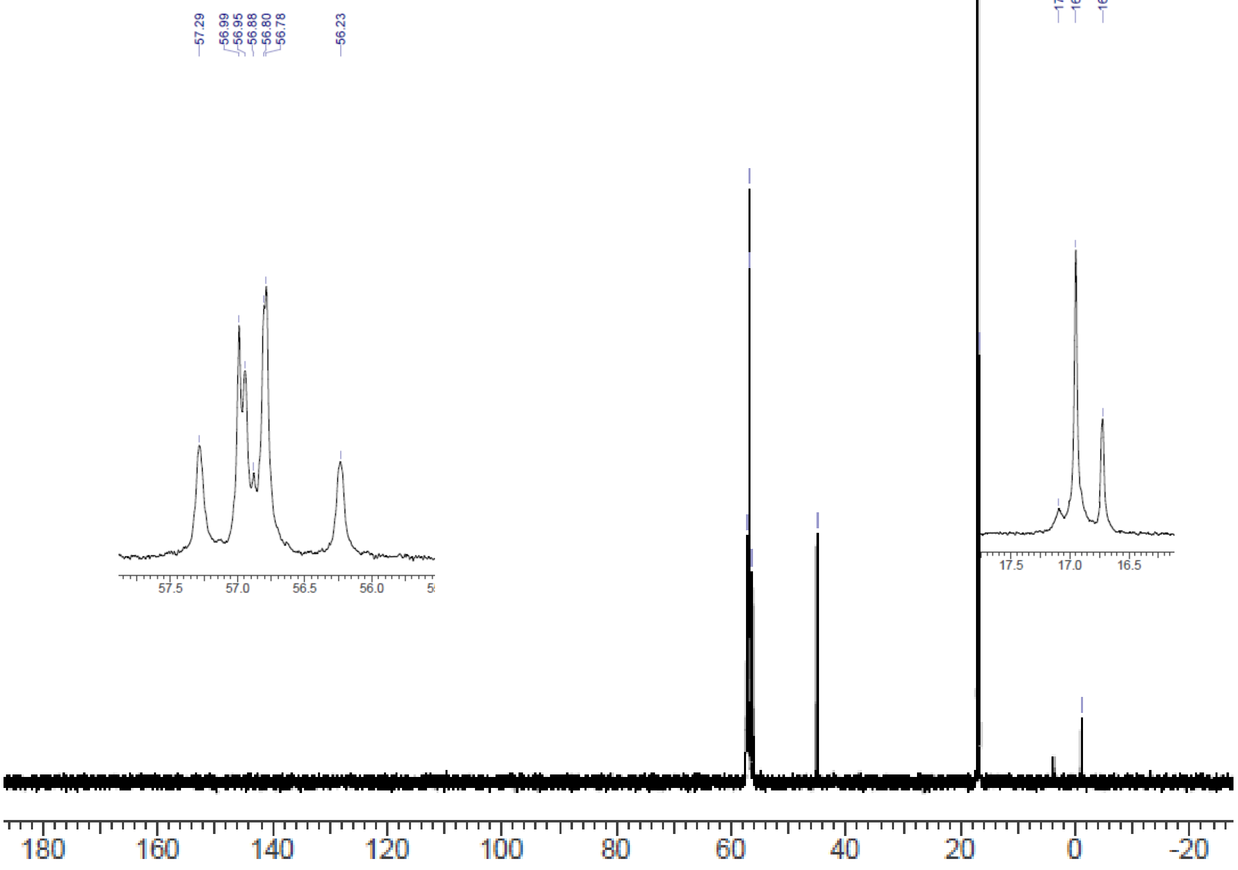

#### Compound S58 (11-mer)

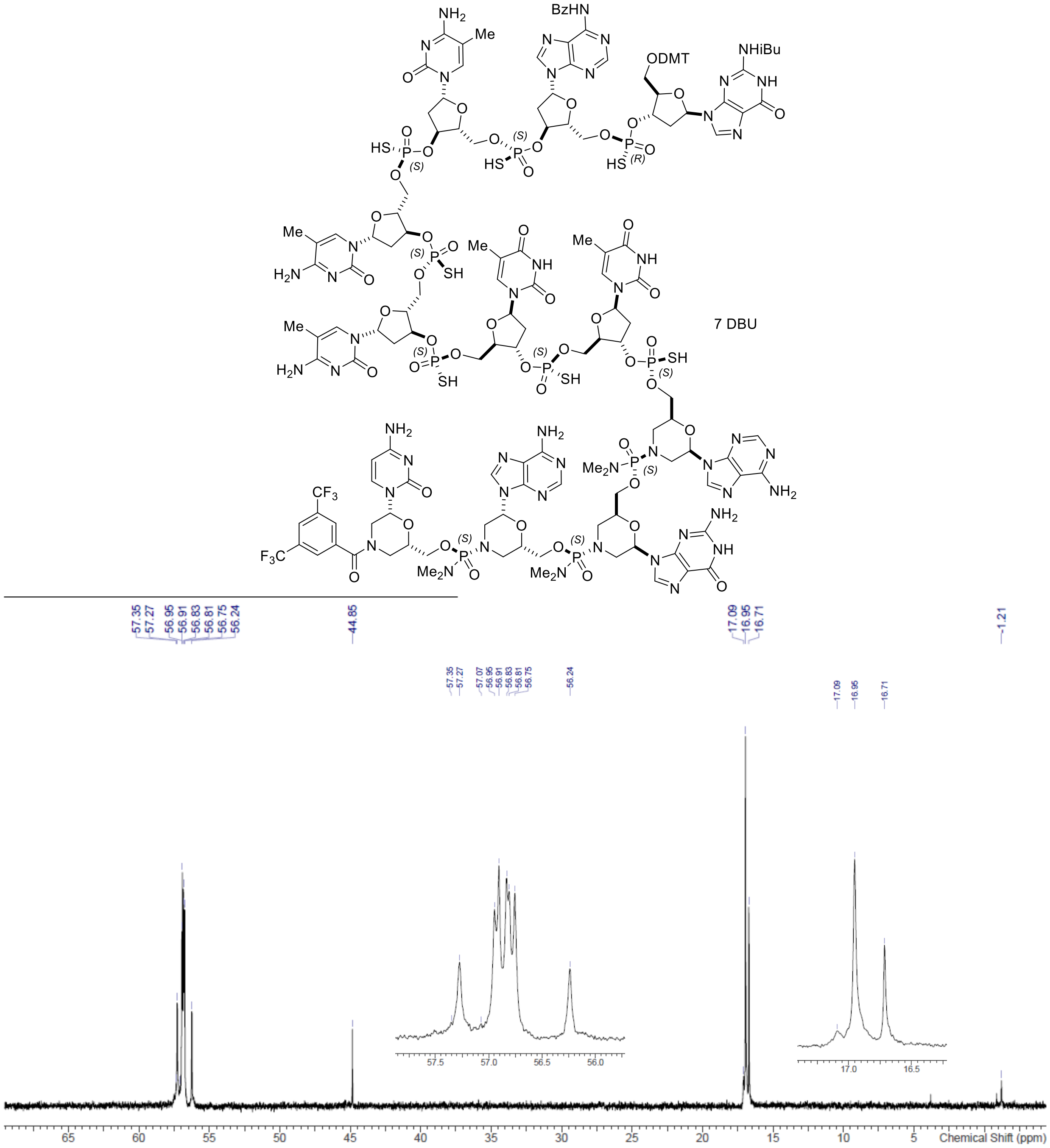

#### Compound S60 (12-mer)

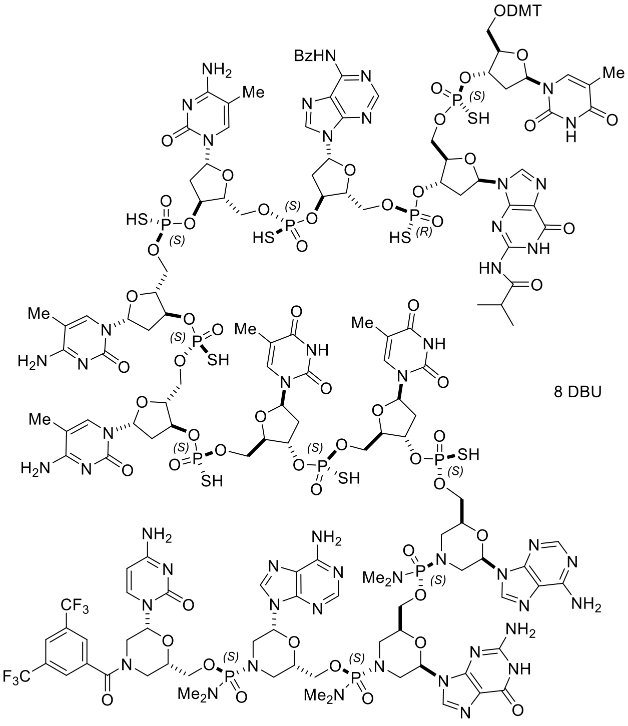

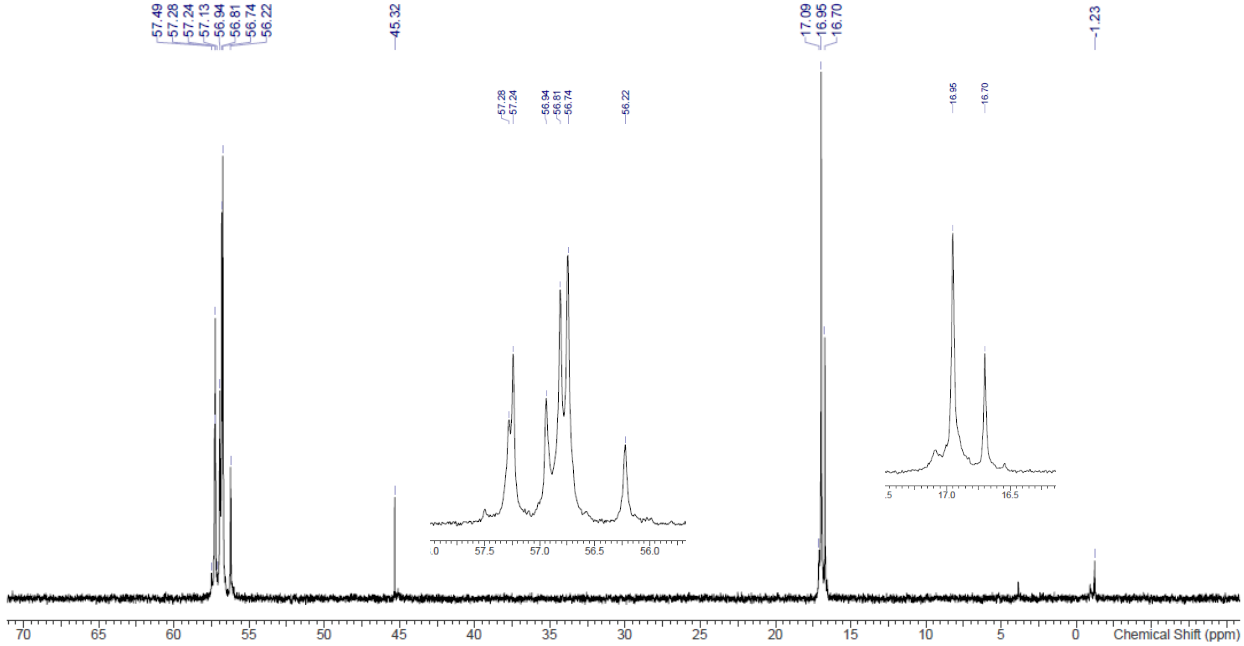

#### Compound S62 (13-mer)

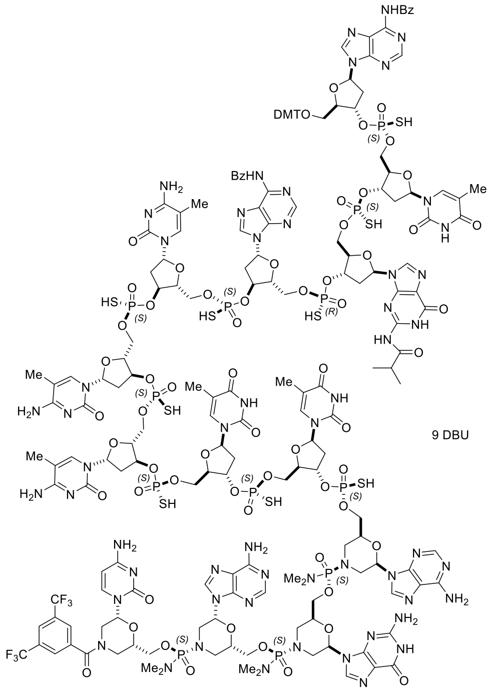

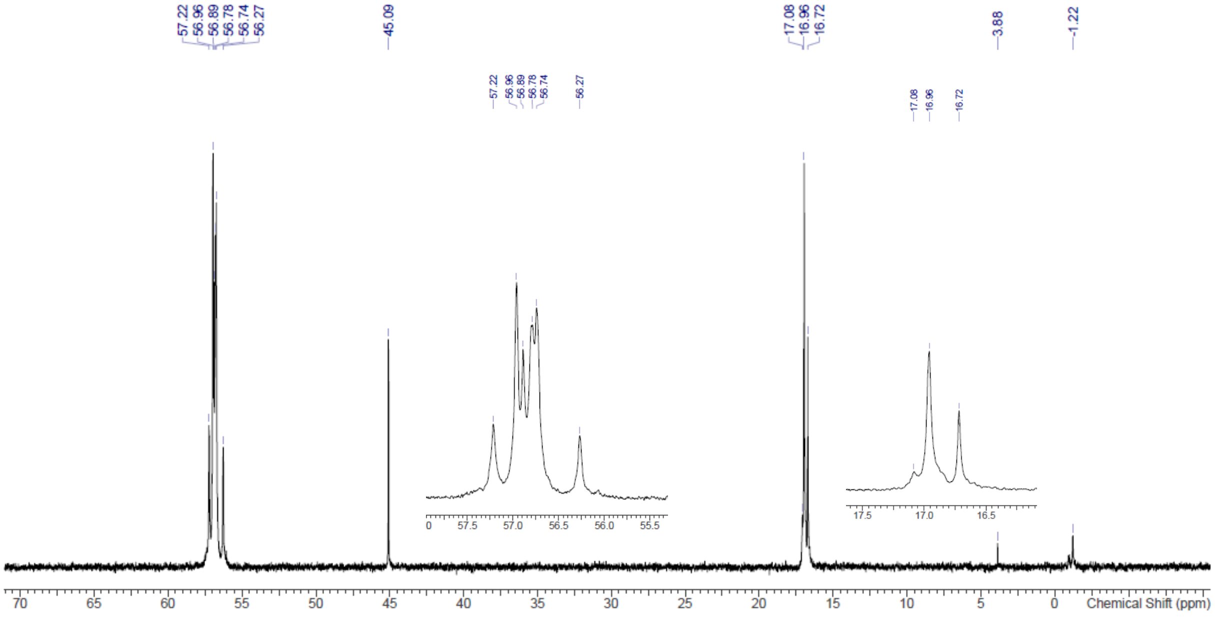

#### Compound S65 (18-mer)

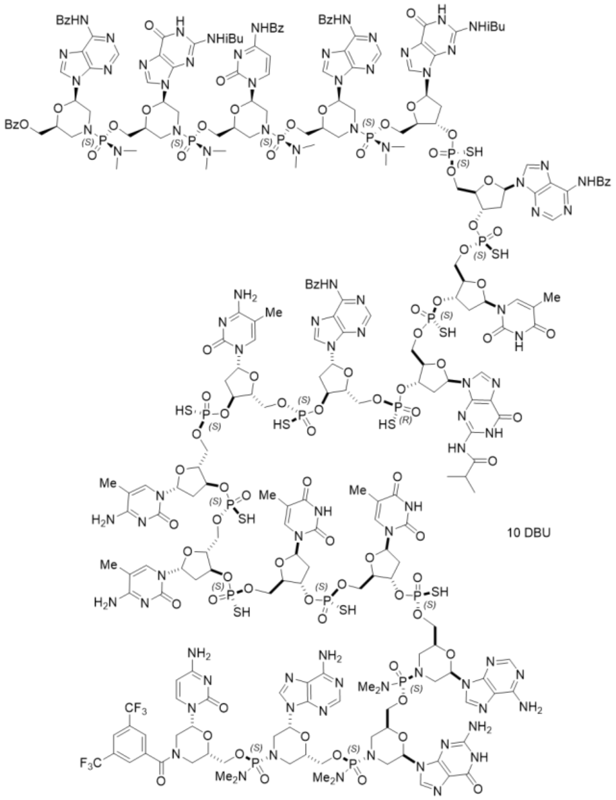

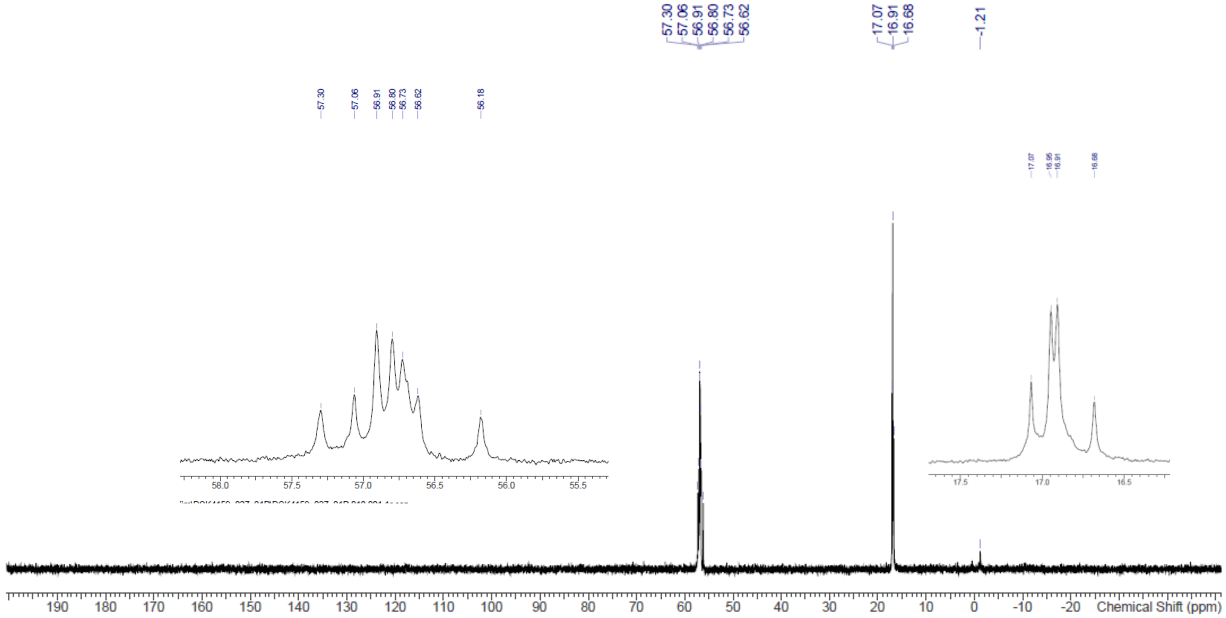

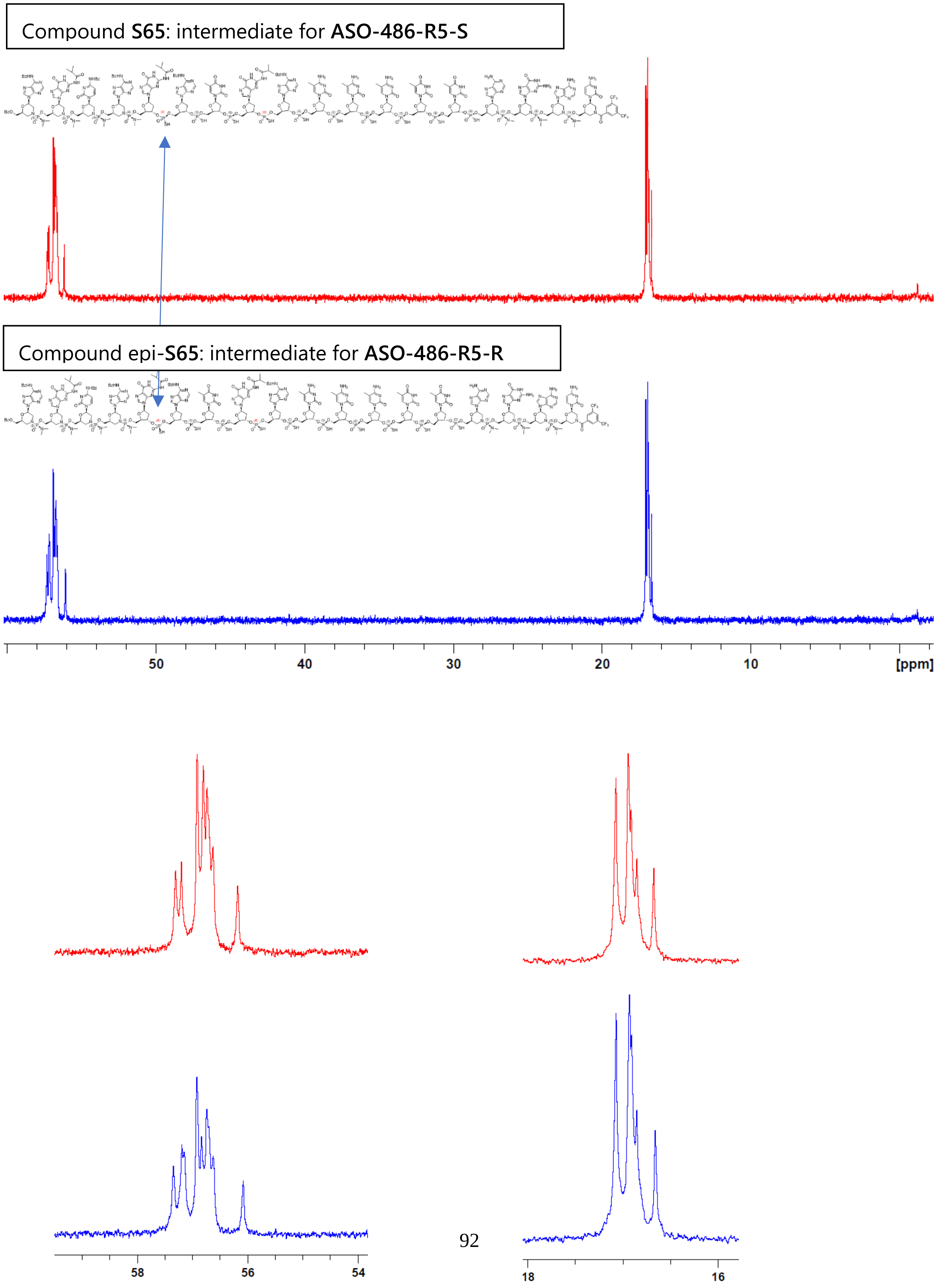
Compound S65: intermediate for ASO-486-R5-S
Compound epi-S65: intermediate for ASO-486-R5-R
50
40
30
20
10
[ppm]
58
56
54
18
16

### 7. HPLC chromatograms of selected compounds

#### HPLC chromatograms of ASO-486-R5-M, ASO-486-R5-R and ASO-486-R5-S

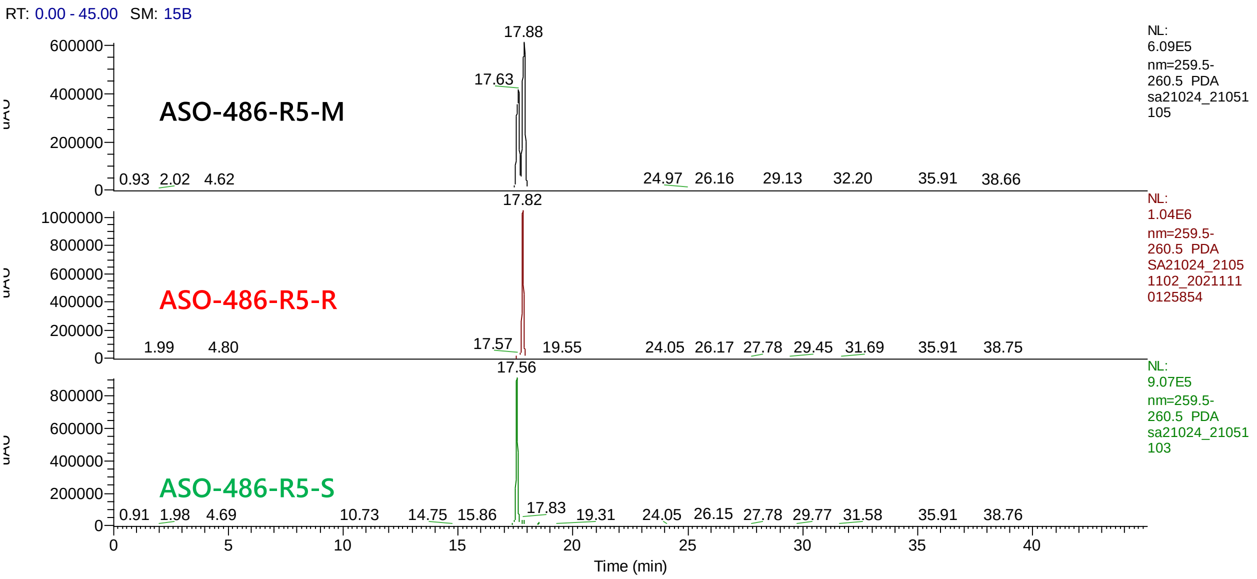

#### Overlay of HPLC chromatograms

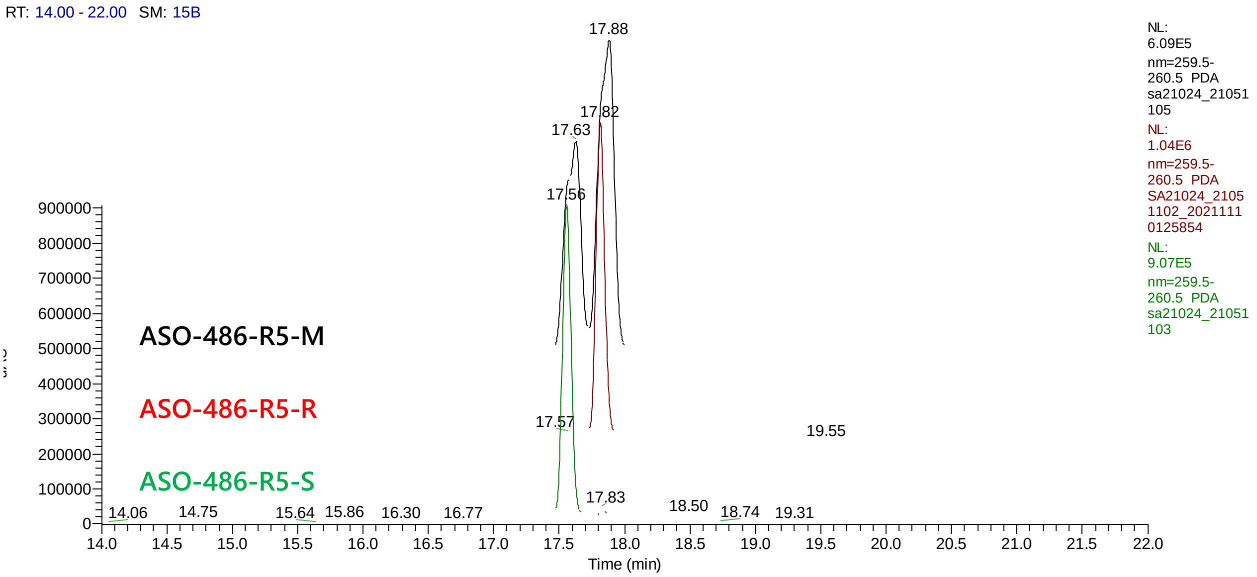
